# Building pangenomes for domesticated and wild tree species: genomic complexity and strategies

**DOI:** 10.1101/2025.07.22.665893

**Authors:** Ismael Blanchard, Quynh Trang Bui, Alexis Mergez, Sukanya Denni, Amandine Cornille, Alexis Groppi, Johann Confais, Ludovic Duvaux, Véronique Decroocq, Benjamin Linard

## Abstract

Long-read sequencing and pangenomics are revolutionizing crop research by providing more complete genome information and revealing crucial structural variations linked to important agricultural traits. Building on recent advances in intraspecific pangenome construction, this study addresses the challenge of creating broader, cross-taxon pangenomes, using the *Armeniaca* taxonomic section as a model. Leveraging a diverse panel of genome assemblies, we constructed a pangenome graph and cataloged associated single nucleotide polymorphisms (SNPs) and structural variants. We characterized the diversity of these variants and assessed the extent to which different taxa contribute to overall pangenome expansion. Additionally, we evaluated the performance of low-depth sample mapping to the graph-based reference, highlighting key technical limitations that may affect the quality of downstream analyses. We further identified specific subsets of SVs that exhibit associations with particular classes of transposable elements. As a case study illustrating the potential functional and phenotypic relevance of graph-derived SVs, we examined the genomic configuration of the *DAM* locus within the *Armeniaca* pangenome.

## Introduction

Recent advances in long-read sequencing technologies have enabled the assembly of more complete and accurate plant genomes, advancing crop biology and breeding. However, single-reference assemblies represent only a fraction of genomic diversity, limiting our understanding of species-wide and taxa-wide genetic variation (Li *et al*., 2022*a*). Pangenome studies often involve assembling genomes from multiple individuals from related and inter-fertile species, and reveal complex structural variations that play a critical role in adaptation and domestication **(Lye and Purugganan, 2019)**. Structural variations affect gene content and repetitive elements, influencing agronomically important traits like stress resistance **(Dolatabadian *et al*., 2020)**. Despite challenges in computational infrastructure, emerging approaches aim to shift from single-reference genomes to pangenomes in crop improvement and ecological studies **(Schreiber *et al*., 2024)**. Most pangenome construction tools developed to date have been designed for intraspecific pangenomes **(Gong *et al*., 2023)**. Transitioning from species-level pangenomes to taxa-wide pangenomes presents a challenge, which we have addressed using the *Armeniaca* taxonomic section as a proof-of-concept.

Among the *Armeniaca* section, *Prunus armeniaca L.* (Rosaceae) (Li and Bartholomew, 2003) corresponds to both cultivated apricots and its wild forms which are still growing in natural populations in Central Asia (Decroocq *et al*., 2016). The initial domestication event occurred around 2,900 years ago, originating from a natural population of southern Central Asian apricots and spreading towards China. A subsequent domestication event took place approximately 2,200 years ago, originating from a northern Central Asian population and expanding into the Mediterranean basin (Groppi *et al*., 2021). Several other species within the genus Prunus, such as *P. sibirica L., P. mandshurica,* and *P. mume,* are considered close relatives to the cultivated apricots (Rieger, 2006).

The *Armeniaca* section is characterized by its high diversity and significant phenotypic variation, which has evolved since its divergence from *Amygdalus* species around 7 million years ago (Groppi et al, 2021). Despite the fact that *Armeniaca* species are inter-fertile and possess collinear genomes, population studies based on a single reference genome often fail to capture critical phenotype-modulating genetic variants such as single nucleotide variants (SNVs), structural variations (SVs), and copy number variations (CNVs). This limitation, known as reference bias, becomes more pronounced as genetic diversity increases, hindering the development of effective breeding strategies (Golicz *et al*., 2020). Pangenome approaches have been introduced to overcome this challenge, with variation graphs representing genetic diversity across multiple genomes (Golicz *et al*., 2020). A pangenome aggregates all genetic variations, including SNPs and large SVs, within a species or species complex. This aggregation can extend beyond the species level to include closely related species, often referred to as super-pangenomes or meta-pangenomes (Li *et al*., 2023). Pangenome graphs have proven effective in characterizing intra and interspecific genetic variability, enhancing the extraction and analysis of complex genetic traits (Liao *et al*., 2023). Several plants of agronomic importance, including rice, cabbage, grapevine, and tomato, have been studied using pangenome graphs to capture complex SVs, some of which are restricted to wild types and absent in cultivars (Bayer *et al*., 2020; Cochetel *et al*., 2023; Li *et al*., 2023). These studies have revealed valuable genetic diversity that can be used in breeding programs. Recently, the first *Prunus* pangenome graph was published for *P. persica* L., which identified key transposon variations linked to fruit coloration (Chen *et al*., 2025). These advancements demonstrate the potential of pangenome graphs to uncover genetic traits that are vital for crop improvement and adaptation.

We present here an Armeniaca metapangenome comprising 25 chromosome-scale assemblies of three *Armeniaca* species, including European and Chinese common apricots (*P. armeniaca)*, Siberian (*P. sibirica*) and Mandchourian (*P. mandschurica*) apricots. SVs in the *Armeniaca* pangenome were identified using publicly available whole-genome short and long read data panels. The expanded variation catalog reveals structurally complex loci, and the pangenome’s accuracy was validated through the analysis of specific loci. We assessed the integration of new variations into the model, confirming its potential to enhance read mappings of novel sequence datasets. This approach reduces the bias associated with using a single genome as a reference. Finally, variations related to a cluster of dormancy-associated genes (DAM genes) were explored, demonstrating the pangenome graph’s potential for future studies.

## Material and methods

### Genomic dataset

Long-read assemblies for 30 Armeniaca genomes were compiled : 15 of *Prunus armeniaca*, 5 of *P. sibirica*, one of *P. mandschurica*, one *P. hongpingensis*, one *P. zhenghensis* (Suppl.File_S1 : table 1). One long read and one short-read assemblies were also retrieved for *P. mume*. Moreover, we recovered a set of related Prunus species as outgroups : three *P. salicina* (Japanese plum). Out of these, for this study we *de novo* assembled : two wild Central asian *P. armeniaca* (KZ_150_8 and KR_091_1 from W2 and W1 genetic clusters, respectively, Groppi et al, 2021) and two *P. sibirica* genomes (CH240 and CH250. All data sources and sequencing approaches for the 32 assemblies are detailed in Suppl Table T1. Details about the new assemblies can be found in Suppl. File S1, section 1.

**Table 1.** : Global SNP and Indel Metrics per Genomic Group.

|  | # SNP /<br>assembly | # SNP<br>density per<br>kb | # Indels<br>/ assembly |
| --- | --- | --- | --- |
| EU | 781833 | 4.24 | 16601 |
| CH | 974612 | 4.25 | 20501 |
| W | 1095549 | 4.53 | 23177 |
| Sib | 1260631 | 5.85 | 26909 |
| Man | 1393252 | 6.23 | 28766 |
| Total |  |  | 115954 |

### Data curation

For each chromosome of each genome assembly, we retained the longest scaffolds, for a total of 8 chromosome sequences per genome. Prior to the pangenome construction, we verified the integrity of each sequence using two different steps. First, we clustered all sequences using their similarity and via the method implemented in the PGGB sequence partitioning script (Garrison *et al*., 2024). Briefly, we compressed and indexed our files using samtools v1.10 (Li *et al*., 2009), fastix v0.1.0 (github.com/ekg/fastix) and used mash v2.2.2(Ondov *et al*., 2016) to obtain a distance matrix, using the option ‘-s 10000’ to specify the sketch size (the amount of representative k-mers summarising each genome). We then inspected the kmer distance matrix to evaluate the similarity between scaffolds from all chromosome sets, as a means to confirm the correctness of their labelling, as well as potential unexpected proximities. Distance heatmaps and dendrograms were generated using R (R Core Team, 2023) and the package pheatmap v1.0.12. Second, we verified global synteny between chromosomes with the D-genies v1.5.0 tool (Cabanettes and Klopp, 2018), using the web version (dgenies.toulouse.inra.fr). When needed, we renamed the mislabelled chromosomes following the standard nomenclature for *Prunus armeniaca* genetic map (Genome Database for Rosaceae, www.rosaceae.org) and we harmonized chromosome orientations with seqkt v1.3(github.com/lh3/seqtk) using the -r option.

### Strategy for iterative chromosome graph construction

We build chromosome pangenome graphs using minigraph-cactus (Hickey *et al*., 2023), which introduces some key choices related to graph construction. This tool relies on an iterative approach in which a guide tree defines the order of graph augmentations, each assembly being iteratively aligned and introduced to the graph. By design, each ordering can potentially end in a different final graph (see (Garrison and Guarracino, 2023)). As a consequence, two criteria will impact the graph construction process: a) the choice of the “reference” (starting) assembly, recommended to be of high quality and/or richly annotated, and b) the order of assemblies for graph augmentation, which should ideally represent phylogenetic proximities, so that potential genome alignment artefacts may not be introduced.

For the first criteria, the Rojo_HCUR haplotype was used as our initial backbone for pangenome construction. It was chosen due to its method of genome assembly, which provides a haplotype-resolved and chromosome-level genome (Campoy *et al*., 2020). It represents the ‘Currot’ haplotype of the Rojo Pasión cultivar, a hybrid between the European apricot ‘Currot’ and the North-American Orangered ^®^ cultivars (Campoy *et al*., 2020).

To define the order of following assemblies, we followed their degree of genetic divergence relative to Rojo_HCUR, by computing a kmer-distances and building a neighbour-joining phylogeny for all assemblies, using the pantools method (Jonkheer *et al*., 2022). Briefly, for each chromosome, the KMC (Kokot *et al*., 2017) and the pantools ‘kmer classification’ commands were used to obtain a Mash distance matrix between all chromosomes (kmer_size = 19). From this matrix, a Neighbor-Joining phylogenetic tree was built with the phangorn v2.11.1(Schliep, 2011) and ape v5.1-7 R packages(Paradis and Schliep, 2019). Each chromosome tree was visually inspected with Dendroscope v3.8.10 (Huson *et al*., 2007) while the Robinson-Foulds (RF) distance (Robinson and Foulds, 1981) was used to verify topological consistency between trees. The online tool iTol (Letunic and Bork, 2007) was used to generate tree figures and taxonomic relationships were compared with the Rehder taxonomic classification of *Armeniaca* (Rehder, 1986). This also helped to approximate taxonomic relationships for several assemblies that were not listed by this source.

### Pangenome graph construction

We used the singularity image v3.9.9 (Kurtzer *et al*., 2017), retrieved from github.com/sylabs/singularity), that included the minigraph-cactus pipeline v2.6.13. For each run per chromosome, the NJ phylogeny (with outgroups pruned) was given as input to the pipeline, ensuring controlled ordering during construction and Rojo_HCUR was set as the starting assembly. Two important parameters were manually set : --filter 1 allowed all nodes to be retained in the graph, including those that are unique to a single haplotype and --reference was set to a list containing all haplotypes in their phylogenetic order, ensuring that all bases are represented in the final graph. This is crucial for population-level analyses where genetic diversity across individuals needs to be retained. Other noticeable options used are --all-snarls to include all complex variations in indexes that will be associated with the graph and --gbz-translation to pack the graph in gbz format, where all haplotypes can be translated into our reference assembly coordinates. Finally, Minigraph-Cactus produces three output graphs by default (‘full’, ‘clip’, and ‘filter, see tool documentation). Every result from our study is based on the ‘full’ graph, where all bases from input assemblies can be retained.

Simultaneously to its construction, VCF (Variant Calling Format) files were generated. Because a VCF will project graph variants to sequence coordinates of a single selected assembly, we produced as many VCF as assemblies in the graph. While one could directly call the options --vcf and --vcfReference to do so, we experienced issues with the singularity image (the vg deconstruct command was failing in some cases). Consequently, we ran these extractions manually after construction, using a more recent version of the vg toolkit (compared to the version included in the minigraph-cactus singularity imag) (Garrison *et al*., 2018). First we extracted the paths with command ‘vg paths’ to obtain the normalised names of all graphs paths (e.g. normalized assembly names) then we used vg v1.57 to perform the ‘vg deconstruct’ command and obtain the targeted VCFs.

### Variants extraction and statistics

Graph and variant statistics were generated with four approaches. First we used the Pan1c workflow (forgemia.inra.fr/alexis.mergez/pan1c) using all chromosome graphs as inputs. This workflow contains a module to generate general and pan genome-oriented graph statistics, as well as providing visualisation tools to estimate the quality of the graph.

Second, we ran the Pansel tool (Zytnicki, 2025), which aims to extract graph regions holding significantly conserved (conversely diverging) regions, relatively to the full content of a graph. We ran the tool using Rojo_HCUR as the reference and used the default window size of 1000. As in the original paper of the method, only the extreme 5% of the fitted log-normal distribution were retained to define conserved and divergent regions. We ran the method once per chromosome graph (to detect conserved/divergent regions within) and once for the full pagneome (to detect these regions among chromosomes).

Third, we ran vcftools v0.1.16 (Danecek *et al*., 2011). using the VCFs projected to ROJO_HCUR assembly to obtain insertions/deletions (indels) statistics (option --keep-only-indels) and SNP statistics (--remove-indels option). Results were analysed using the R dplyr (Wickham *et al*., 2025) and ggplot2 (Wickham, 2016) packages. We analyzed indels within a size bin of 50 bp to 100,000 bp, this upper bound being the maximum length that the pan1c workflow will consider, thus allowing direct comparison between the tools.

Finally, Panacus plots (Parmigiani *et al*., 2024) were generated to estimate the variant enrichments brought to the graph by each assembly. Assembly order was set manually using the -O option and followed the ordering used at construction. The -q parameter was used to define quorum thresholds, which correspond to the minimum proportion of haplotypes in which a genomic segment must be present to be included in the output. A quorum of 1.0 (i.e., 100%) identifies the core genome, comprising regions shared by all haplotypes, while a quorum of 0.0 includes any sequence present in at least one haplotype, thus representing the entire cumulative amount of new bases brought by each haplotype.

### Transposon content

To identify potential transposons, the extracted indels were binned by size, and each subset was then independently analyzed using BLAST version 2.2.28 (Altschul, 1997). This analysis was complemented by a custom transposon library built from 11 assemblies selected to maximize genetic divergence. Briefly, for each of these assemblies, we used the TEdenovo pipeline (Flutre et al., 2011) from the REPET package v3.0 (urgi.versailles.inrae.fr/Tools/REPET) to generate a new transposon library, followed by three rounds of curation based on TEannot (Quesneville et al., 2005; Ahmed et al., 2011) where any consensus not supported by at least one full-length copy in the assembly was discarded (Jamilloux *et al*., 2017). The 11 curated libraries were then concatenated into a pan-library of apricot species, and a final redundancy filtering step was performed using PASTEC (Hoede *et al*., 2014) to retain a single representative for each transposon definition sharing ≥98% of length and ≥95% of sequence identity. Using this custom library, we used BLAST version 2.2.28 (Altschul, 1997) to obtain the best transposon alignments for each indel. To select the most robust and relevant transposons for each indel, we applied a set of hierarchical criteria designed to minimize redundancy and enhance result interpretability. First, only transposon hits with a minimum sequence identity of 90% were retained to ensure high-confidence matches. Then, priority was given to transposon size (larger elements being more likely to represent complete and informative annotations), followed by lower e-values, higher sequence identity, and longer alignment lengths. In cases of equal ranking, alignments with fewer mismatches and fewer gaps were preferred. Hits without an assigned order or class were also excluded from the dataset to maintain classification consistency across the analysis.

To allow meaningful comparisons of transposon density across both indel size bins and chromosomes, independently of differences in available sequence space, we applied two complementary normalization strategies. First, the number of transposons detected within each indel size bin was normalized by the total aligned base pairs in that bin (`align_len`), accounting for the fact that larger bins naturally contain more sequence and may therefore accumulate more hits. Second, transposon counts per chromosome were normalized by the average chromosome length, using size data derived from high-quality haplotype assemblies.

These normalization steps enhanced the biological interpretability of the density patterns observed. Variability and confidence intervals of the proportion of transposons within indels and chromosomes were estimated through a bootstrap procedure of 1000 replicates each. To test for statistically significant differences, we performed Pearson’s chi-square tests, which confirmed a significant pattern between TE order distributions and both indel size bin and chromosome. In addition, pairwise proportion tests were carried out for the top five TE orders between different bins and chromosomes.

### Short read mappings

To evaluate the potential of our pangenome graph in thoroughly detecting SVs, we compared the mapping of a panel of 325 Illumina short-read accessions to the graph versus to the reference assembly Rojo_HCUR. This panel, described in Groppi et al. (2021) and representing the diversity of the Armeniaca collection maintained at the French CRB at Bordeaux, was sequenced at depths of up to 20X. To allow reads to map preferentially to the chromosome holding the most similar sequence, in particular when homologous sequences (or translocations) are present on more than one chromosome, all chromosomes were combined. We concatenated all Rojo_HCUR chromosomes into a single FASTA subsequently indexed with samtools faidx and minimap2 -d. Read mapping on the reference was performed using minimap2 v2.28 (Li, 2018). Mappings were computed using the short-read parameter preset (-ax sr) and the parameter (-R) keeping read group information for downstream processing. Sample metadata were introduced in the resulting BAM files using samtools view -bh, ensuring compatibility with following analyses. The mapping to the graph was performed using the VG toolkit v1.20 (Sirén *et al*., 2021). Before mapping, chromosome graphs were combined using vg combine -p. The combined graph was then indexed with vg autoindex and the giraffe pipeline (-w giraffe) to generate a GBZ binary. The read mapping was performed using vg giraffe with paired inputs (-p) and output alignments in GAM format. We also set the -N and -R options to assign sample identifiers to the results. To allow direct comparison between the two approaches, each GAM was then projected to Rojo_HUC sequence coordinates, by generating a BAM file (e.g. graph-based coordinates are translated into linear coordinates of a selected reference).

We used samtools 1.20 (Li *et al*., 2009) to compute general statistics from the BAM output of both methods. To achieve a fair comparison between Minimap2 and VG Giraffe results, we carefully set up the FLAG-based filtering by applying the ‘-F 2308’ filter to all BAM files (see Suppl. File S1, section 8 for more details). Mapped read counts and associated MAPQ score distributions were then extracted using ’samtools view‘ and ’samtools stats‘ commands.

### Structural variation over the DAM region in Armeniaca

To extract the tandem-arrayed DAM (Dormancy-Associated MADS-box) gene region from the graph, we performed a homology analysis based on DAM1 to DAM6 genes. We used Uniprot sequence identifiers from (Balogh *et al*., 2019), with DAM1 to DAM4 and DAM5 to DAM6 originating from *P. mume* and *P. armeniac*a, respectively. Candidate homologous regions were identified by aligning gene nucleotide sequence to all assemblies using BLAST+ v2.2.28+ (Camacho *et al*., 2009) as well as, for DAM5 and DAM6 only, translated proteins of the MADS box domains extracted from UniProt (identifiers: A0A4D6CKB8_PRUAR; A0A4D6CHU2_PRUAR). We checked the validity of these candidate regions by aligning their sequences to known DAM coordinates a) from the Marouch assembly, as well as b) annotated DAM regions for all assemblies referenced in the Genome Database for Rosaceae (GDR). We retained alignments with the highest identity and length to define putative DAM coordinates. As Rojo_HCUR is not available in GDR, we used BLAST coordinates from the Marouch assembly, easier to inspect via JBrowse, to retrieve node IDs for each DAM gene. These were then mapped onto the subgraph previously extracted from Rojo_HCUR coordinates using the VG toolkit. Subgraphs were visualized with Bandage. (Wick *et al*., 2015).

The gene cluster structure was verified utilizing the JBrowse tool integrated within the Genome Database for Rosaceae (GDR), as described by Jung et al. (2019). This platform facilitated the comprehensive visualization of available transcriptomic data, enabling its direct correlation with BLAST outputs. Furthermore, JBrowse provided crucial insights into the architecture of the assemblies, specifically aiding in the identification and characterization of genomic rearrangements (e.g., loops) present in only a subset of the assemblies.

To unequivocally confirm the presence of the DAM region in the *P. mandshurica* CH 264 genome, despite its absence from its primary assembly, we used a targeted approach using PromethION long reads (ENA accession: ERR4656977). The raw ONT sequencing reads were used to build a BLAST database. Reads matching the DAM regions were then identified by searching the database using candidate DAM sequences as queries. The specific commands and methodological details for this analysis are comprehensively documented in https://forge.inrae.fr/benjamin.linard/armeniaca_pangenome_paper : `04_4_READ_ME_blast_and_methodology_DAM.md`.

## Results

### Genome assemblies and preliminary steps for graph construction

For this study, we built a comprehensive dataset of 32 Prunus genome assemblies, each obtained from diverse sequencing technologies and research groups (Suppl.File_S1, section 5). To facilitate the interpretability of our results, we categorized these assemblies into distinct groups based on their phylogenetic proximity, geographic origin, and agronomic utility. This classification strategy also broadly aligns with similar groupings established by Groppi et al. (2021). Our grouping scheme is as follows:

- **European (EU) Group:** This group comprises assemblies of European origin, including Rojo_HCUR, Marouch, RougeR_H1, RougeR_H2, Rojo_HORA, and Stella.
- **Wild (W) Group:** Representing wild Central Asian apricot genomes, this group includes five assemblies that correspond to the W1 and W2 genetic clusters identified by Groppi et al.: KZ150 H1, KZ150 H2, KR091 H1, KR091 H2, and CH320.
- **Chinese (CH) Group:** This group consists of three cultivated *P. armeniaca* genomes from China: Sungold, Meihua, and GSYX.
- **Siberian (Sib) Group:** This group encompasses five *P. sibirica* genomes (CH240 H1, CH240 H2, CH250 H1, CH250 H2 and F106) along with one phylogenetically close *P. armeniaca* accession (Longwongmao).
- **Manchurian (Man) Group:** The *P. mandshurica* genome (CH264) forms its own distinct group.
- **Hybrid (HYB) Group:** This group includes four *P. armeniaca x P. sibirica* F1 hybrids: RRxCH240A H1, RRxCH240A H2, RRxCH240B H1, and RRxCH240B H2.
- **Outgroup Assemblies (Mum, Sal, Other):** The remaining assemblies, serving as outgroups, are detailed in Suppl.File_S1, section 5.

Our dataset comprises both haplotype-resolved and unphased (mixed-haplotype) genomes. To streamline our analysis, we will uniformly refer to each distinct input sequence, whether it represents a single haplotype or a mixed assembly, as an independent "assembly." This approach is rooted in the operational mechanisms of pangenome graph tools, which interpret every input sequence as a unique "path" within the graph. For example, the two resolved haplotypes of KZ150 (H1 and H2) will be represented as two independent "assemblies" and, consequently, two distinct paths in the graph. Similarly, the unphased assembly of CH320 will be considered a single "assembly," corresponding to one path in the graph. This standardization ensures consistent processing and interpretation across all diverse assembly inputs.

Notably, the Rojo_HCUR assembly was selected as our “reference” genome for this study. It is based on the assembly of two high-quality, chromosome-level haplotype-resolved genomes of *P. armeniaca* (common apricot), using a technique called gamete binning. This method leverages high-throughput single-cell sequencing of pollen grains, allowing for the separation of reads into two haplotypes, which are then independently assembled. This approach results in highly accurate, haplotype-specific assemblies, with each haplotype (e.g., ’Currot/Rojo_HCUR’ and ’Orange Red/Rojo_HORA’) scaffolded to the chromosome level, demonstrating greater than 99% haplotyping precision and alignment accuracy with parental genomes. The separated haplotypes avoid the issues common to pseudo-haploid assemblies and resolves each haplotype distinctly, thus ensuring a higher quality reference for future population-level analyses and GWAS that rely on structural and sequence-level precision. With these applications in mind, Rojo_HCUR assembly is a reliable reference genome for apricot pangenome construction (as a reference is necessary for the graph construction tool, as described in next paragraphs). Combined with the fact that haplotype is a cultivated european and is known to be less diversified than its wild counterparts, one makes it a good candidate for reference.

Following an initial evaluation of assembly quality via k-mer distances, six assemblies showed pervasive inconsistencies in chromosomal k-mer distances and sizes for all of their chromosomes, indicating a lack of harmonized chromosome labeling. This necessitated the relabeling of corresponding sequences, a process thoroughly detailed in Suppl. File S1, section 2. In addition, the GSYX assembly showed multiple instances of reversed sequence orientations which were subsequently corrected manually, as also described in Suppl. File S1, section 2.

Following these preliminary assembly corrections, we constructed a comprehensive phylogeny using all available assemblies. This phylogenetic tree serves as a guiding framework for the subsequent pangenome graph construction, providing essential evolutionary context to inform the graph’s topology and interpretation (Figure 1).

**Figure 1:**
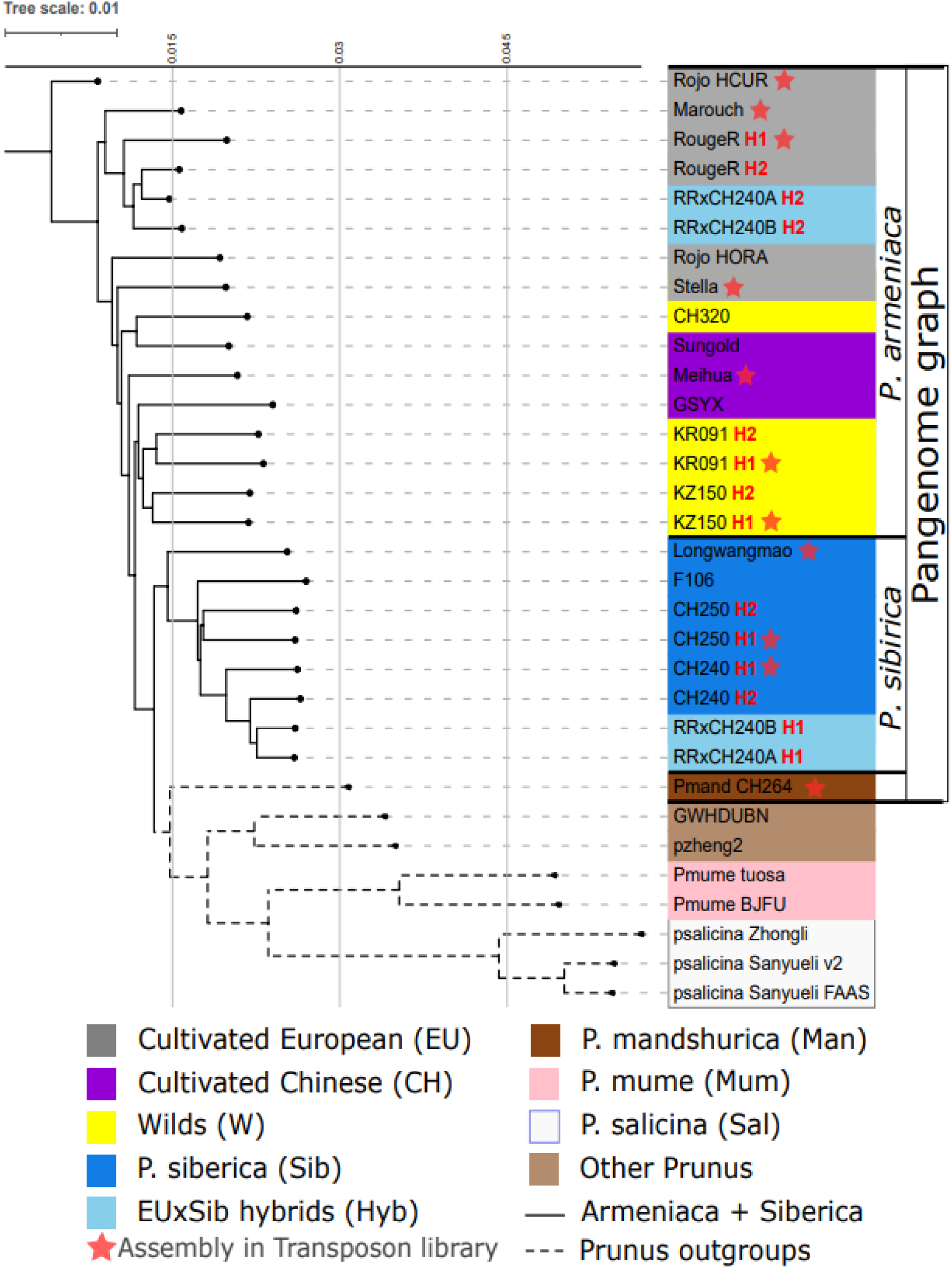
Tree of Armeniaca genome assemblies. This tree aims to define an order for pangenome graph construction (see methods), and as a consequence is rooted on the reference genome Rojo_HCUR and not outgroups. Wherever possible, individual haplotypes are maintained as distinct leaves within the tree, designated by red labels H1 and H2.

The tree (Figure 1) reveals several noteworthy placements. Firstly, As expected, haplotypes in hybrid assemblies segregate according to their parental origin (e.g., H2 within the European group and H1 within the Siberian group, as illustrated in Figure 1). Second, the phylogenetic position of Meihua and GSYX (both from the Chinese group) makes the Wild group non-monophyletic. Last, despite its classification as *P. armeniaca* in the Genome Database for Rosaceae, Longwangmao is positioned as basal to the *P. sibirica* lineage. To proceed further, we decided to proceed with this k-mer distance-based placement, given that the pangenome graph construction tool we employed utilizes a similar metric to determine assembly alignment order.

Based on our phylogenetic analysis and the substantial genetic distances observed relative to the reference assembly, Rojo_HCUR (detailed in Section 3 of Suppl. File S1), we opted to exclude most outgroup members from the pangenome graph construction. Specifically, *P. salicina*, *P. mume, P. zhengheensis* and *P. hongpingensis* were removed with *P. mandshurica* retained as the sole outgroup. This decision was primarily driven by the understanding that pangenome graph tools are fundamentally designed for species-centric analyses and may yield suboptimal alignments when incorporating distantly related genomes, which could impact graph quality and could generate erroneous genetic variants artifacts. Consequently, our final dataset for graph construction comprised 25 assemblies (14 phased and 11 unphased). We re-evaluated the phylogenetic relationships using this reduced assembly set; the resulting tree showed no significant topological changes compared to the full version, as evidenced by a Robinson-Foulds distance of 2, indicating only two differing splits. This refined tree was subsequently utilized as the guiding input for pangenome graph construction (Suppl. File S1, section 4).

### Armeniaca pangenome graph

Our pangenome graphs, constructed for each chromosome, exhibited a substantial complexity (see statistics in Suppl. File S1, section 5). The number of nodes per graph varied from 2.7 million to 6.7 million – with an average node length ranging from 27 to 41 base pairs – indicating SV occurrences every few dozen of base pairs approximately. As expected, the largest chromosomes contained the highest number of nodes (e.g. Chromosome 1 with 6,669,924), while the smallest, Chromosome 5, had only 2,697,166 nodes. This trend was also reflected in the mean path length (where each path represents one assembly), which ranged from 18.9 Mb (chromosome 5) to 46.1 Mb (chromosome 1). The compression factor, defined as the ratio between the total sequence length of all input assemblies and the total number of base pairs in the graph’s nodes, ranged from 4.01 (chromosome 2) to 6.38 (chromosome 5), reflecting differences in sequence redundancy and local structural complexity across the chromosomes.

The Pansel analysis of our pangenome graphs revealed a varied distribution of significantly conserved and divergent regions (under Pansel definition) across chromosomes. The number of conserved regions ranged from 124 on chromosome 8 to 296 on chromosome 1. Similarly, the number of divergent regions varied from 579 to 1524, also across chromosomes 8 and 1, respectively. When normalized by the reference chromosome lengths (Suppl. File S1, section 5, table_T2), chromosome 5 consistently showed a higher proportion of both conserved and divergent regions. Interestingly, chromosome 2, rather than the smallest chromosome (chromosome 5), exhibited the lowest number of both conserved and divergent regions. This suggests that genetic variations on chromosome 2 are more evenly distributed along its length, rather than concentrated in specific highly conserved or highly divergent loci.

#### Contribution of assemblies to Pangenome variation

The various assemblies and their respective phylogenetic groups contributed differentially to the genetic variations observed within the pangenome graph (Figure 2). As expected, incorporating additional assemblies led to a reduction of the core pangenome and an expansion of its other components. Focusing on the European (EU) group, we observed an initial increase in total genomic content from 45.5 Mb to 84.6 Mb, followed by a plateau. This saturation suggests that adding more assemblies may not significantly introduce novel genomic variations. Concurrently, within the same group, the core genome declined from 45.5 Mb to 37.8 Mb, stabilizing across the RRxCH240 to Stella additions. This observation further reinforces previous findings regarding the core genome’s stability within the same lineage (Jalil *et al*., 2022).

**Figure 2:**
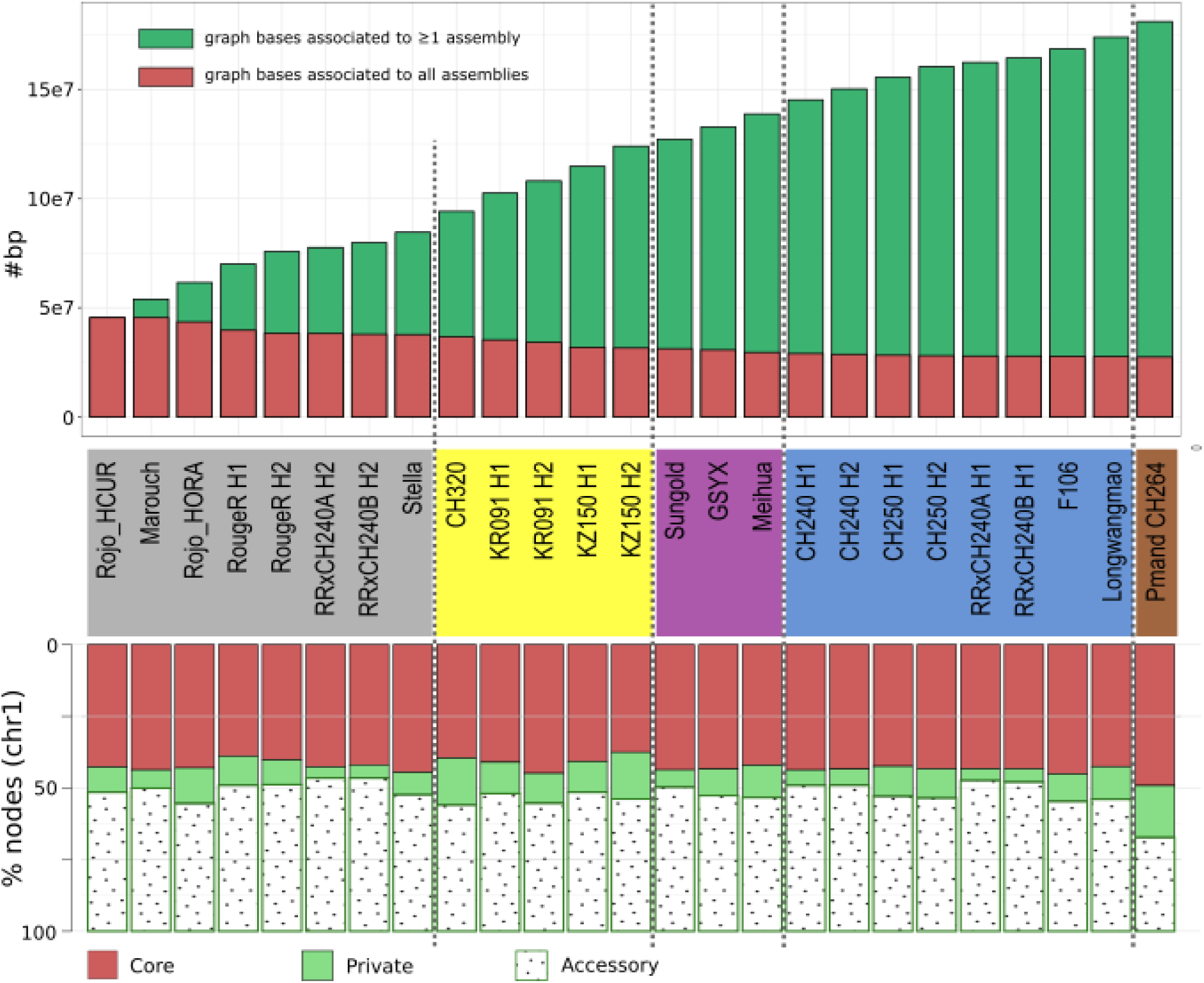
Assembly contribution to pangenome variant diversity (Example: chromosome 1). Assemblies and their associated groups are delineated along the X-axis, with distinct color coding: European (EU) group in grey, Wild (W) in yellow, Chinese (CH) in purple, Siberian (Sib) in blue, and Manchurian (Man) in brown. A. The upper histogram quantifies the increase in pangenome content, measured in added base pairs with each newly-added assembly (from left to right). Red bars represent the core genome bases, defined as sequences shared by all analyzed assemblies. Green bars denote the total number of bases present in at least one assembly, encompassing the core, accessory and private genome components, as per the definitions employed by the Panacus tool. B. Individual assembly composition per pangenome component. Red fraction represents the core genome, comprising graph nodes shared by all assemblies; Dotted fraction shows the accessory genome, encompassing graph nodes shared by more than two assemblies; Green fraction indicates the private genome, consisting of graph nodes unique to a specific assembly.

#### Pangenome dynamics and core genome stability

The progressive integration of assemblies from various geographic origins significantly influenced pangenome size and core genome composition (Figure 2). The addition of Wild (W) accessions triggered a substantial expansion of the pangenome, increasing its total size from approximately 94.2 Mb to 124.0 Mb, representing an average gain of ∼8 Mb per genome. Concurrently, the core genome observed a contraction from 37.7 to 31.6 Mb. Subsequent incorporation of Chinese (CH) assemblies continued this expansion, albeit with a reduced rate of accessory genome acquisition (∼4.9 Mb per genome). The core genome further diminished to 29.6 Mb. The addition of Sibirica (Sib) genotypes resulted in a more modest pangenome expansion, contributing approximately ∼4.3 Mb per genome with a gradual erosion of the core genome, which ultimately stabilized at 27.6 Mb. It is noteworthy that the RRxCH240A and RRxCH240B haplotypes contributed minimally to the overall pangenome increase, likely due to their hybrid nature derived from parental assemblies (RR and CH240) previously incorporated into earlier graph iterations. Finally, the Manchurian assembly contributed an almost exclusively accessory fraction of approximately ∼7.2 Mb. This observation supports the hypothesis that the pangenome retains the capacity to further expand with the inclusion of additional outgroup accessions. Strikingly, the core genome demonstrated remarkable stability across the Sibirian and Mandchourian groups. This suggests that a robust definition for the core genome within the *Armeniaca* section may have been achieved. Other chromosome ordered growth histograms values are shown in Suppl. Table T3 (Suppl.File_S1 section 5).

#### Dissecting assembly contributions to pangenome architecture

The contribution of each assembly to the pangenome was further analyzed by quantifying the proportional distribution of core, accessory, and private graph nodes (Figure 2, bottom histogram). Using chromosome 1 as an illustrative example (with proportions for all chromosomes detailed in Suppl. File S1, section 5, Table T4), core genome contributions ranged from 37% to 49%. The core genome variability across assemblies may be surprising but is attributable to the representation of repeated sequence patterns within the graph; a single node representing a repeat can be traversed multiple times by an assembly’s path, leading to repeated counting. Therefore, a higher core fraction in this representation may indicate potential duplications of core pangenome segments or ubiquitous genome repeats with variable copy numbers across assemblies. In contrast to the core genome, contributions to private and accessory fractions were highly variable. Private contribution reached up to 18% for *P. mandshurica,* with the Wild (W) group also exhibiting substantial contributions of private nodes. As expected, the *P. armeniaca × P. sibirica* hybrids consistently showed low private fractions, a consequence of their parental haplotypes already being present in the graph.

Analyses of other chromosomes (Suppl. File S1, section 5, Table_T4) revealed significant variations in the proportions of core, private and accessory genomes. For example, on chromosome 2, the core genome averaged a mere 11%, while private regions could constitute up to 36% (e.g. *RRxCH240BH2*, *P. mandschurica*). On chromosome 3, core fractions ranged from 17% to 38%, with accessory content dominating in certain haplotypes (e.g., *CH_320*, ∼49%). Conversely, the core genome exceeded 50% for some assemblies on chromosome 5, while private fractions spanned from negligible (<1%) to substantial (>30%). These diverse profiles underscore the chromosome-specific and lineage-specific nature of shared genomic variations within our pangenome.

#### Genomic variation within the pangenome

Our pangenome graph revealed significant genetic diversity across accessions. We observed a total of 25 million SNPs relative to the reference Rojo_HCUR (Table 3). SNP counts ranged from 1.4 million for Pmand_CH264 to over a total of 10 million for the Sibirica group ( Suppl. File S1, section 7, table_T8). While most groups showed consistent SNP density per assembly, it significantly increased for the Sibirica and Manchurian groups reflecting their divergences to the Armeniaca group.

The graph also contained over 537,000 **indels** relative to Rojo_HCUR ( Suppl. File S1, section 7, table_T8), with their number per assembly strongly correlated with phylogenetic distance – the *P. sibirica* and *P. mandshurica* groups showing nearly twice as many indels as the European group same observation can be said for the number of SNP per assembly (Table 1). Chromosomes 2 and 4 showed consistently lower SNP contributions, suggesting stronger evolutionary constraint. ( Suppl. File S1, section 7).

Number of variants detected in the graph for each group (Sib, W, EU, CH, Man) relative to the reference Rojo_HCUR. Raw number of indels and SNP, are normalized per assembly within groups. SNP density was calculated by normalizing the raw variant counts to both the number of assemblies and the total number of base pairs within each group, and is expressed per kilobase. The last row shows totals across all assemblies.

Indel length analysis indicated that the smallest bin (50-200 bp) accounted for 40% of all pangenome indels while the largest bin (5000-100000 bp) accounted for only 7.5% (Figure 3). Notably, two specific bins (200-400 bp and 800-1800 bp) displayed distinct peaks in indel counts across all chromosomes and phylogenetic groups (details for all bins in Suppl. File S1, section 7, table_T8).

**Figure 3.**
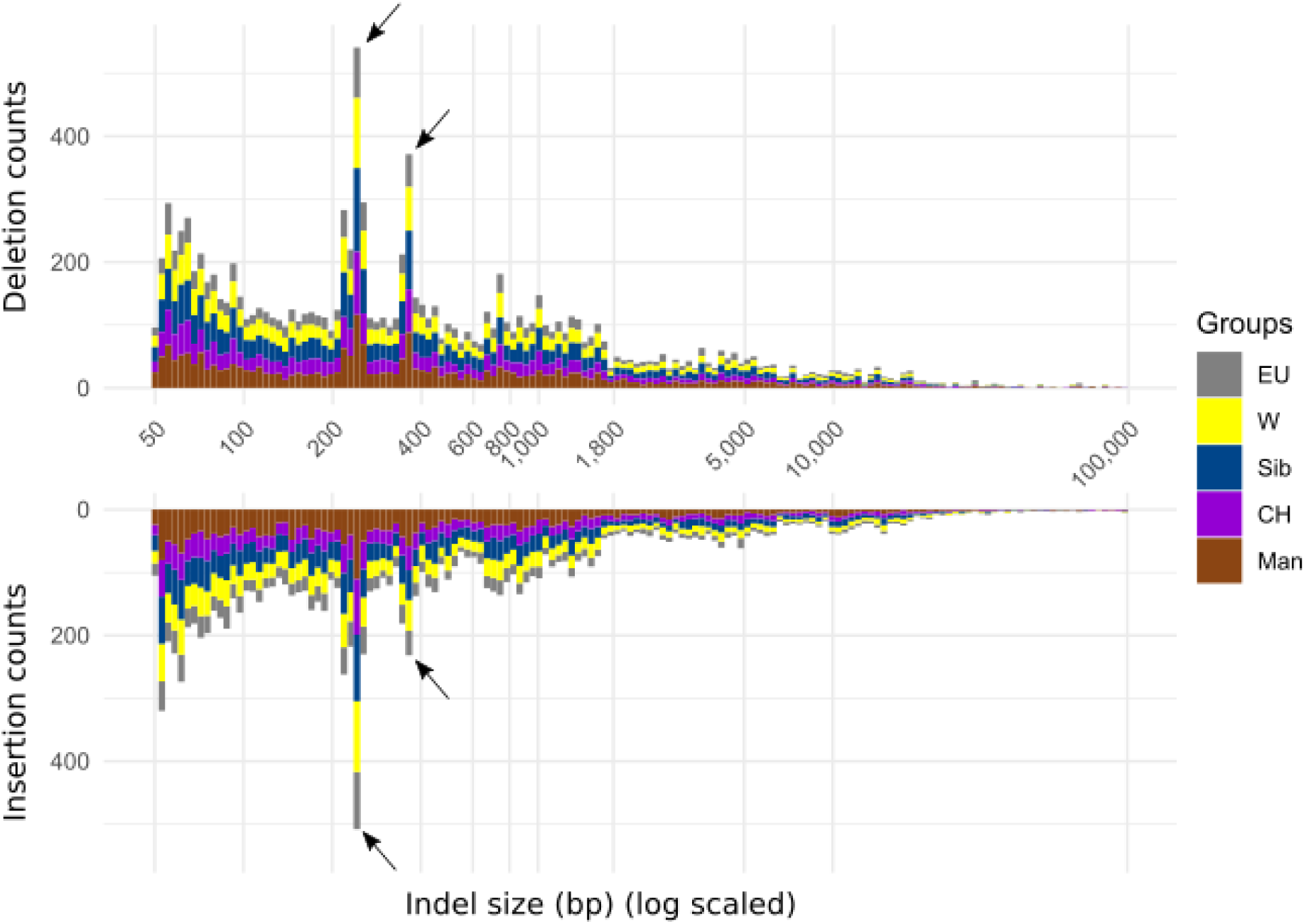
: Indel frequency and length polymorphism on chromosome 1. The figure presents the distribution of insertions (top histogram) and deletions (bottom histogram), relative to the Rojo_HCUR reference genome. The x-axis, on a log10 scale, represents the sorted lower bounds of indel length intervals (bins). The Y-axis indicates the mean number of indels observed within each phylogenetic group: European (EU), Wild (W), Chinese (CH), Sibirica (Sib), and Mandchourian (Man), arrows point out the highest peaks in the 200-400 bp bin.

#### Transposable element landscape within the pangenome

To investigate the transposable element landscape, we first created a pan-library of 19,136 consensus elements using the REPET package. This set was subsequently refined to 8,032 non-redundant definitions (Suppl. Data D1). Utilizing this pan-library, we identified a total of 42,679 TEs among all graph indels. Transposon density was largely consistent across chromosomes, averaging one detectable transposon every 18kb (Suppl. File S1, section 7). However, TE counts varied considerably based on the considered indel length bin. Notably, the largest number of TEs (8,557) was recovered from the 200–400 bp indel bin, while the 600–800 bp bin was associated with the lowest count of 1,940 TEs (Suppl.File_S1, section 8, Table T9) provides detailed statistics per bin).

Classifying TEs into five orders (LARD, LTR, TIR, TIR|LTR, Unclassified) revealed an uneven distribution across indel length bins. LARD elements were notably enriched in the 400–600 bp bin, comprising over 30% of transposons, compared to less than 10% in other bins. In contrast, LTR elements showed consistent proportions (over 20%) across bins (Figure 4 and Suppl. File S1, Table T9). The proportion of unclassified TEs increased in proportion for indel bins exceeding 1,800bp.

**Figure 4.**
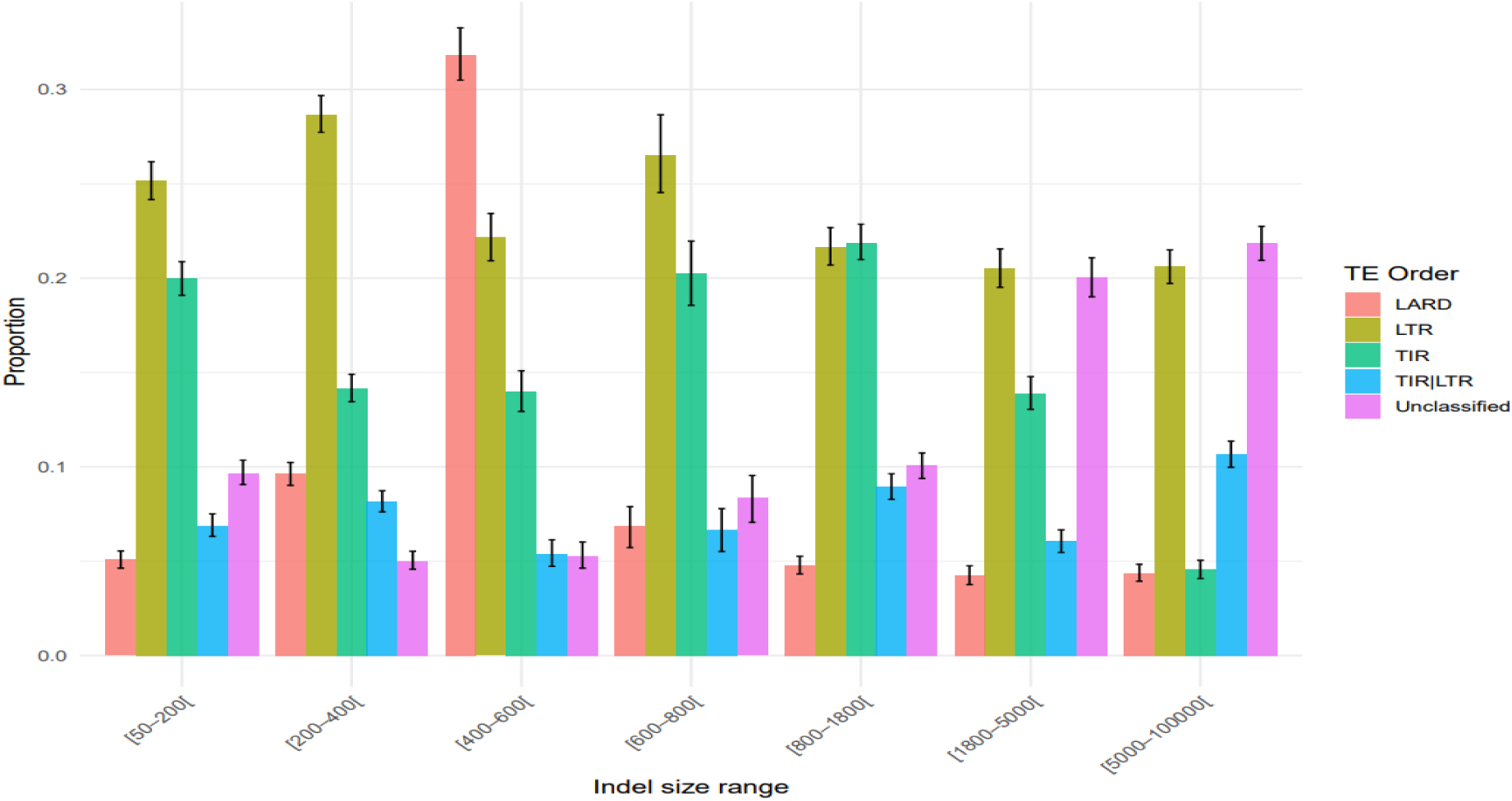
: Proportional representation of the five most frequent Transposable Element (TE) orders across indel length bins. Each bar represents the mean proportion of a specific TE order within a defined indel length interval. These means were derived from 1,000 bootstrap replicates and the 95% bootstrap confidence intervals are indicated by error bars. Note that the sum of proportions for a given bin may not reach 100%, as the figures exclusively showcase the five TE orders with the highest overall frequencies.

We also found that TE order composition varied across chromosomes and size bins; for instance, LTR element proportions differed between Chromosome 1 and Chromosome 3, and between specific indel size bins (Suppl.File_S1, Table 9). Figure 5 provides a detailed view of how different TE orders are non-randomly distributed across the spectrum of indel sizes. An enrichment analysis further highlighted these disparities, revealing a depletion of TIR elements in indels larger than 5,000 bp. (Suppl. Table S1, section 8, Table_T9).

**Figure 5:**
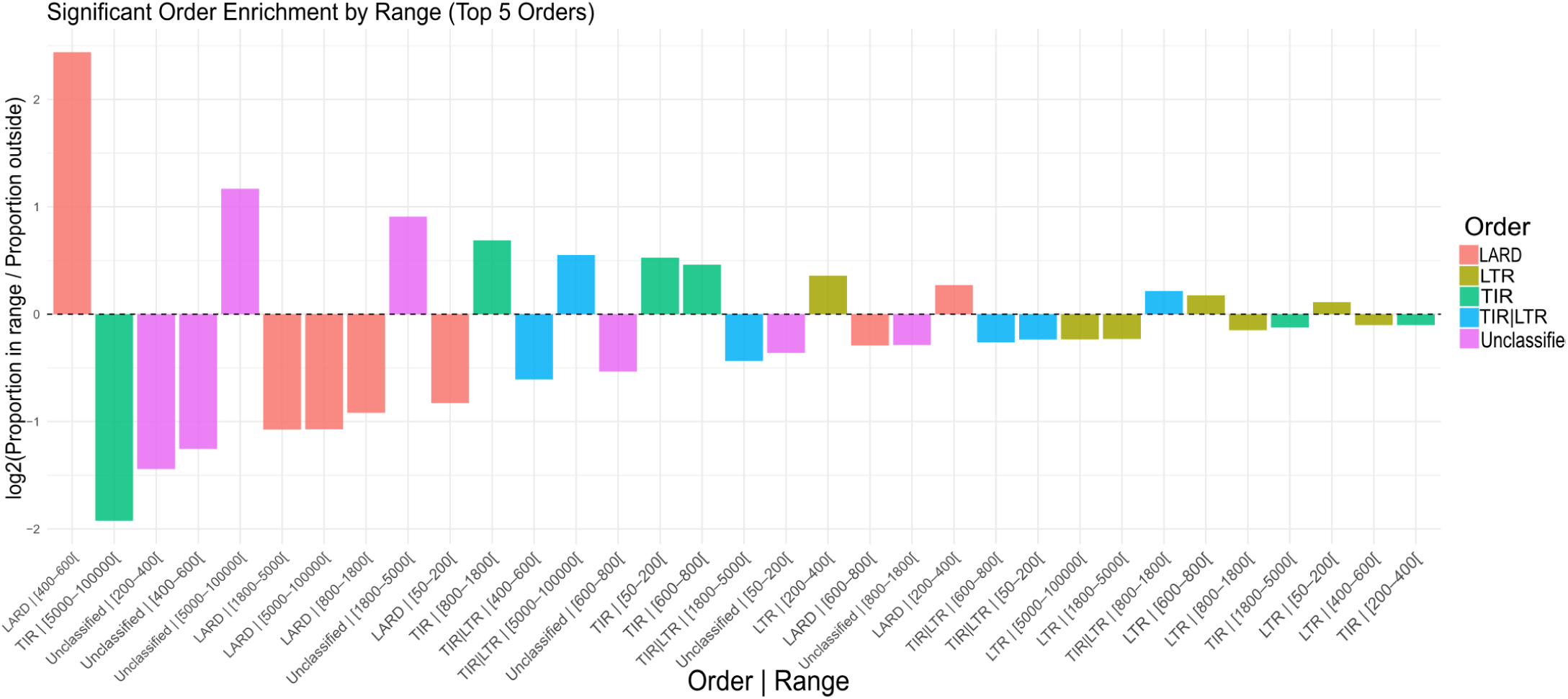
Enrichment and depletion patterns of transposon across indel lengths. The x-axis denotes the association between each TE order and its corresponding indel length bin. Positive values indicate an enrichment of a particular TE order within that indel bin, meaning its proportion is higher than its overall representation. Conversely, negative values indicate a depletion, indicating a lower proportion. The associations are ordered from left to right based on the absolute magnitude of their enrichment or depletion, highlighting the most divergent relationships first.

#### Enhanced mapping with graph-based approaches

Read mapping performance varied considerably depending on the specific tools employed and the methodologies used for projecting to genomes (e.g. how BAM files were generated). Notably, differences in post-mapping filtering capabilities were software dependent (a detailed discussion on this is available in Suppl. File S1, section 9). Specifically, we found that “proper mapping” or “paired segments” flags associated with output BAM files are not directly comparable between current Variation Graph (VG) tools and Minimap2. VG tools appeared to employ a more stringent definition of proper mapping compared to Minimap2. Concurrently, analysis of MAPQ score extraction from all mapped samples confirmed that the corresponding scoring scheme is software-specific invalidating direct comparisons based on MAPQ values. Furthermore, MAPQ distributions generated by Minimap2 and VG Giraffe exhibited substantial differences (Suppl. File S1, section 10). Despite these limitations and to provide an approximate overview of mapping quality variations, we categorized corresponding reads into MAPQ extremes (Table 4).

**Table 4.** : Read counts and proportion of reads associated with MAPQ score extremes. Following the mapping of the 325 short-read samples, reads were classified based on their associated MAPQ. These scores in [0;60] and scores <5 and >55 are associated with a low-quality mapping or a high-quality mapping, respectively (1st column). Line “Filtered-out” corresponds to reads that did not pass the samtools filters preceding mapping, and that are not associated with a MAPQ value. For each pair of tool and MAPQ bin, the left value reports the absolute number of reads (in milliards of reads) and the right value (in parentheses) reports the corresponding proportion among the total of reads.

| MAPQ bin | Minimap2<br>- Rojo_HCUR ref - |  | VG Giraffe<br>- pangenome graph - |  |
| --- | --- | --- | --- | --- |
|  | Count (% of total) |  | Count (% of total) |  |
| ≤ 5 (lowest) | 3.71 | (31.7) | **5.07** | **(43.3)** |
| Other values | 1.14 | ( 9.7) | **1.33** | **(11.3)** |
| ≥ 55 (highest) | **5.59** | **(47.8)** | 4.34 | (37.1) |
| Filtered-out | **1.27** | **(10.8)** | 0.97 | ( 8.3) |

This analysis revealed that Minimap2 yielded a higher proportion of reads with high MAPQ scores, while graph-based mappings generally had lower MAPQ scores but mapped more reads overall (Table 4).

To refine this picture, Figure 6 reports mapping gains when using the graph, e.g. the proportion of reads that could be mapped on the graph, but not when using the single reference assembly Rojo_HCUR. Crucially, graph-based approaches demonstrated enhanced mapping. All 325 short-read samples showed improved mapping to the pangenome graph, ranging from 0.5% to an outlier of 9.2% increased mapped reads compared to a single reference. A weak inverse correlation was observed: lower sequencing depths yielded higher gains, while samples with over 60 million reads converged to a stable 2.4% gain.

**Figure 6:**
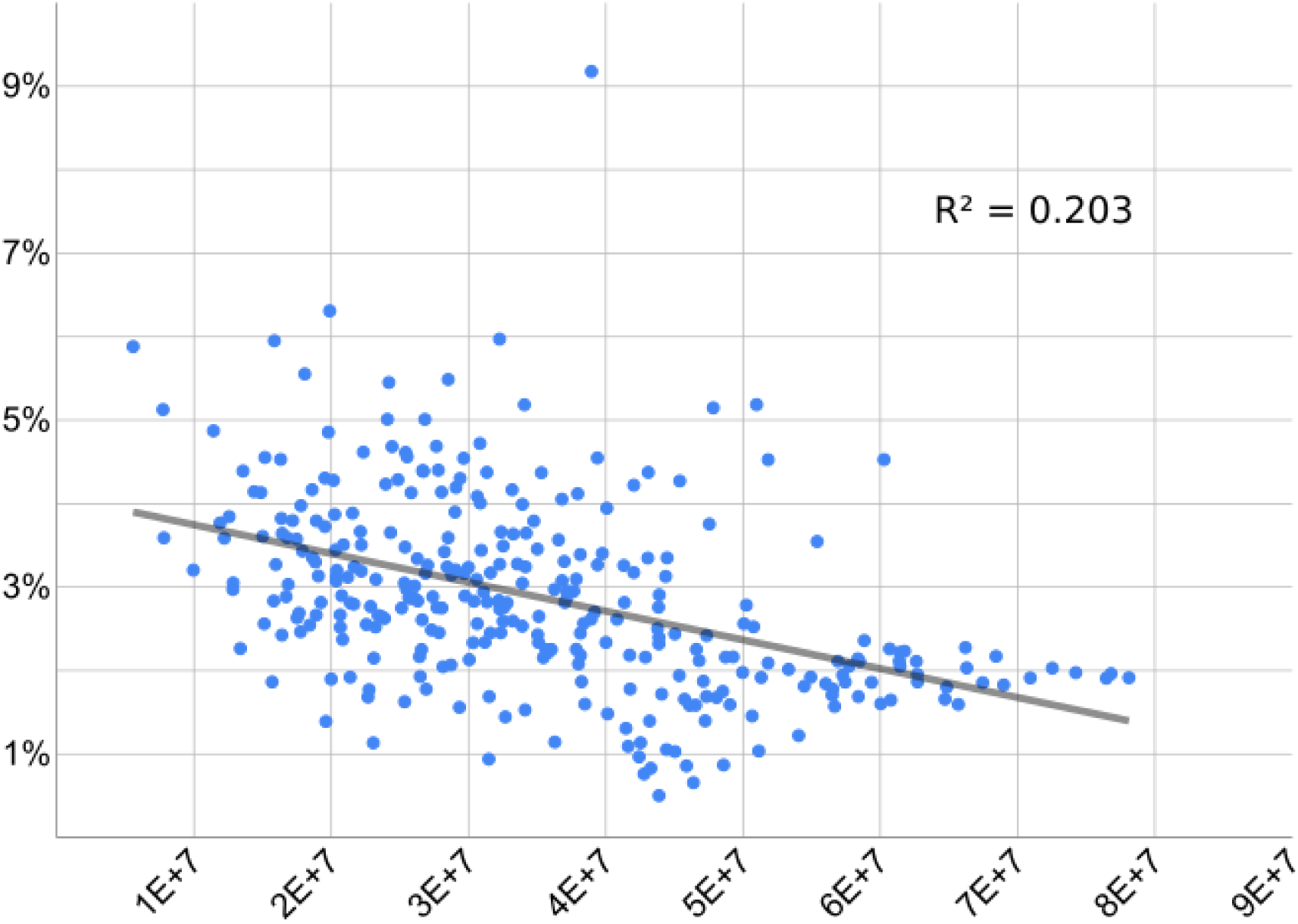
**Pangenome Graph Enhances Read Mapping for Low-Depth Accessions**. This analysis quantifies the **mapping gain** observed for 325 low-depth short-read accessions when mapped to a pangenome graph, as opposed to a traditional single-reference mapping approach using *Rojo_HCUR*. The X-axis represents the total number of reads generated for each accession, while the Y-axis displays the percentage of supplementary reads successfully mapped to the pangenome graph that were not mapped to the *Rojo_HCUR* reference. The grey line illustrates the linear regression between these two variables, revealing a weak negative correlation between sequencing depth and mapping gains.

#### Characterisation of structural variation in the DAM gene locus within the pangenome

As a showcase of the pangenome utility, we focused on the Dormancy-Associated MADS-box (DAM) gene locus to exemplify SVs within the pangenome graph, given their conserved synteny and phenotypic impact (Balogh *et al*., 2019; Quesada-Traver *et al*., 2022). Figure 7A displays the genomic organisation of this tandem-repeat locus. Despite general conservation, four prominent graph loops, indicative of large indels, were identified in the DAM gene locus (Figure 7B and Suppl. File S1, section 12), for assembly details and corresponding graph coordinates).

**Figure 7.**
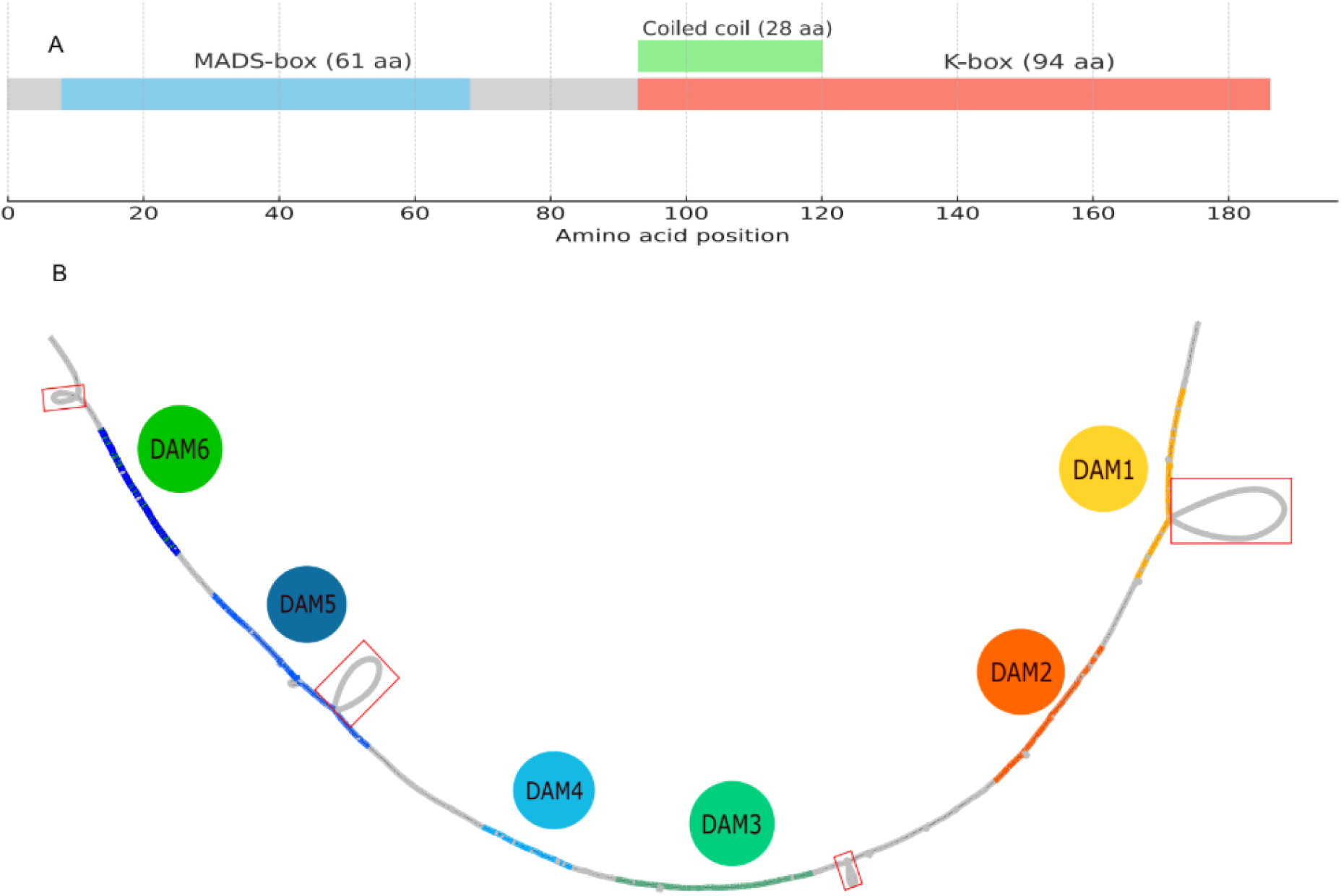
: Schematic representation of the domains of a DAM gene, here DAM5 in *P. armeniaca* and graphical representation of the DAM gene locus extracted from the pangenome graph. A) These genes show a conserved structure that includes at least a MADS-box (in blue) and a K-box (in red), for a final protein product of around 200aa. B) The DAM region was extracted from the graph, with node selection bounded by coordinates −3000 bp upstream of DAM1 and +7000 bp downstream of DAM6, relative to the Rojo_HCUR path. Coloration of individual DAM genes indicates their homology as determined by BLAST alignment to the Marouch assembly. Red rectangles highlight the four largest SVs observed within the region.

Loop 1, a 12 kb insertion within DAM 1, was specific to two assemblies (CH_240 (H2) and RRxCH240_plA (H1)) and posed alignment challenges for Minigraph-cactus. Loop 2, a 2.1 kb variation in the intergenic region between DAM 2 and DAM 3, was shared across 14 assemblies. Loop 3, a 7.1 kb insertion within DAM 5, was unique to Rojo_HORA. The fourth loop, downstream of DAM 6, was shared among seven assemblies (see Suppl. File S1, section 12).

While existing Prunus Genome annotations provided no specific information for these regions (GDR, www.rosaceae.org), our analysis successfully identified transposable elements (TEs) Loop 1 contained a high-confidence LTR element (Gypsy-like transposon, 91.53% identity over 886 bp), and Loop 3 harbored a TIR element (CACTA-like transposon, 94.12% identity over 170 bp). Conversely, no confident TE matches were found in Loops 2 and 4, potentially suggesting different genomic features or highly divergent elements. All corresponding graph and sequence positions are described in Suppl. File S1, section 12.

A notable observation was the complete absence of the DAM gene cluster in the CH_264 assembly within our graph. However, re-examination of PromethION reads (ENA identifier: <u>ERR4656977</u>) for CH_264 confirmed the presence of a DAM gene cluster, suggesting an assembly error, though its precise localization relative to our pangenome remains unconfirmed.

Therefore, the characterization of SVs within the DAM gene locus highlights the utility of pangenome graphs in uncovering complex genomic variations, particularly those associated with transposable element activity, and identifies potential assembly inaccuracies.

## Discussion

In this study, we present a comprehensive pangenome of the Armeniaca section, shedding light on both the genomic contributions of the various genotypes and the structural organization that defines this pangenome.

### Assembly Quality and Curation Challenges

The quality of a pangenome graph is fundamentally linked to the quality and meticulous curation of its constituting assemblies (Michael and VanBuren, 2020). We encountered several challenges during this process, including issues with chromosome labeling and inversions, which necessitated substantial manual intervention. For instance, the GSYX assembly showed signs of chromosome admixture and potential scaffolding issues, as evidenced by its unique coordinate patterns or missing data within the DAM region. Furthermore, the phylogenetic placement of some assemblies, such as Longwongmao, deviated from expectations. We used a distance-based reconstruction that scales well with sets of complete genomes, but remains less accurate than a more advanced phylogenetic study (e.g. careful selection of orthologous loci and the use of a probabilistic model). This choice also explains the relatively long terminal branch lengths of the tree; they mirror the many k-mers that are not shared by genomes and increase dissimilarity. The inclusion of both phased and unphased genomes in this analysis also warrants caution for downstream genotyping applications, given their potential impact on alignment accuracy and variant interpretation. These observations underscore the critical need for rigorous data curation in pangenomics, as inaccuracies can propagate through analyses and complicate the distinction between genuine biological variants and methodological artifacts.

A similar need for extensive manual curation has been observed in other plant pangenomes, particularly in assemblies rich in repeats or with complex SVs. In *Oryza sativa*, for instance, substantial manual efforts were required to validate and refine genome structure across accessions (Zhou *et al*., 2020). Likewise, in *Pisum sativum*, the resolution of centromeric and repeat-dense regions demanded iterative polishing and advanced scaffolding strategies (Kreplak *et al*., 2019; Yang *et al*., 2022). Alternatively, masking these regions may be another strategy considering the lack of tools for properly assembling them. These cases illustrate the widespread nature of such challenges in plant pangenomics, particularly in the context of building graph-based references. Meanwhile, these regions must be effectively masked until molecular tools for their efficient assembly become available Currently, there is a recognized lack of specialized tools beyond basic read mapping and VCF generation that allow comprehensive quality assessment of pangenome graphs and effective discrimination between true biological variation and technical noise. While this study, along with the associated scripts provided on GitHub, offers some practical guidance, more generalized and scalable solutions remain an unmet need in the field.

### Expanding the *Armeniaca* Pangenome

The Armeniaca pangenome continues to expand with the integration of new genotypes, and we anticipate further growth with the inclusion of additional groups and assemblies. As expected, the incremental integration of diverse genotypes systematically reduced the core genome and expanded accessory regions, consistent with previous reports in plant pangenomics (He et al., 2023; Beaulieu et al., 2025). The observed plateau in cumulative growth following the integration of European (EU) genotypes suggests a saturation of genomic novelty within this phylogeographic group, confirming the diminishing returns of extensively sampling closely related individuals (Tettelin et al., 2005). In contrast, the significant genomic gains derived from the incorporation of Wild (W), Chinese (CH), and particularly Manchurian (Man) genotypes underscore the importance of including genetically distant outgroups to capture broader genetic diversity (Li et al., 2023). A similar pattern was observed in Glycine max, where the inclusion of wild relatives (Glycine soja) significantly expanded the pangenome and introduced SVs absent from domesticated accessions (Liu et al., 2020).

This expansion notably occurs at the expense of the core genome, which reached a relatively stable threshold of approximately 27.6 Mb. A similar core genome stabilization has been reported in soybean, where (Liu *et al*., 2020) found that after sampling 27 soybean genomes, the core gene set remained consistent while accessory content continued to increase. The observed stabilization of the core genome suggests the presence of conserved genomic regions that are functionally essential and maintained across all haplotypes of the Armeniaca section. This stabilization trend is consistent with observations in *Oryza sativa*, where analysis of over 3,000 accessions revealed that the core genome plateaued despite the continuous accumulation of accessory variation, indicating a conserved set of essential genes across the species (Wang *et al*., 2018). Importantly, the absence of a definitive logarithmic plateau in cumulative growth across chromosomes further reinforces the observation that continuous pangenomic expansion is likely to happen with the addition of further assemblies, highlighting substantial unexplored genomic diversity.

This expansion was not uniform across chromosomes. When normalized by chromosome length (Suppl. File S1, section 5, table_T2), chromosome 5 despite being the smallest showed the highest density of both conserved and divergent regions, indicating hotspots of structural and sequence diversity. In contrast, chromosome 2 displayed the lowest number of such regions, suggesting a more homogeneous distribution of variation along its length. These differences in chromosomal contribution highlight the importance of considering genome architecture in pangenome analyses and suggest that some chromosomes may play a more pivotal role in shaping intraspecific diversity.

The relatively modest genomic contribution of hybrid haplotypes (RRxCH240A and RRxCH240B) supports the understanding that hybrid genomes primarily represent recombinations of parental genomic variation, rather than significant sources of novel genomic content. This observation aligns with gene-centered pangenome analyses in crops such as sunflower, where hybridization was shown to alter gene presence/absence without dramatically expanding total gene content (Hübner *et al*., 2018). In our case, the limited contribution of hybrid assemblies at the whole-genome level—including intergenic and structural regions—indicates that such genotypes can reasonably be excluded from graph construction, as soon as the parental varieties are included in the graph. Additionally, their low fraction of private sequence argues against major sequencing or graph-related artifacts in their representation.

### Chromosome-Level Pangenome Dynamics and Variation Drivers

The contribution of different assemblies and groups to the pangenome varies significantly across chromosomes. The observation that some chromosomes exhibit a core genome proportion of 50%, while others drop to as low as 11%, warrants further investigation into chromosome-specific evolutionary dynamics in *Prunus*. This variability could be linked to differential selective constraints, recombination rates, chromosomal rearrangements, or historical domestication and other selective events (Groppi *et al*., 2021; Tan *et al*., 2021). Indeed, Groppi *et al*., 2021 showed that half of the 0.5% top genomic regions under selection during domestication map over the chromosome 4. For instance, the CH320 assembly displayed a unique pattern of core/accessory/private proportions across chromosomes, deviating from other members of the Wild group. This suggests a possible ancient hybridization and introgression from *P. sibirica*, aligning with previous hypotheses (Groppi *et al*., 2021).

The considerable influence of the most genetically distant groups on the volume of identified SNPs and Indels is consistent with previously observed correlations between genetic divergence and geographic or taxonomic distances (Jayakodi *et al*., 2024). Regarding Indels, the conserved distribution patterns around 200-400 bp and 800-1800 bp are particularly intriguing. Rather than reflecting direct selective constraints, these recurrent sizes more likely stem from inherent mutational biases linked to the activity of specific transposable element families. Such patterns are consistent with the replication behavior and structural features of TEs, as discussed by(Feschotte, 2008; Oliver and Greene, 2009) who highlighted their pivotal role in genome remodeling through bursts of insertions of specific lengths. Our TE enrichment analysis indeed demonstrated a significant representation of specific TE orders, notably TIRs and LTRs, within these size bins, emphasizing their potential role in shaping genomic diversity via structural rearrangements (Su *et al*., 2019; Morales-Díaz *et al*., 2025). The presence of TE fragments, particularly Gypsy-like and CACTA-like elements, within SVs surrounding the DAM gene cluster suggests a potential mutational contribution of these elements to local genome rearrangements. While such variants could hypothetically influence phenotypic traits such as dormancy (Balogh *et al*., 2019; Quesada-Traver *et al*., 2022), further functional validation is required to assess any causal relationship.. The DAM region also served as a practical illustration of assembly completeness evaluation, as demonstrated by the absence of annotation in the CH_264 assembly in this specific graph region.

### Pangenome Mapping Advantages and Limitations

Pangenome graphs offer a clear advantage over single-reference analyses for SNP and SV detection, and significantly improve read mapping efficiency. Our findings indicate that mapping to the graph increased the number of mapped reads by 0.5% to 9.2%, depending on read depth. However, it’s crucial to acknowledge that conventional metrics for evaluating mapping quality are software-dependent and not directly comparable between graph-based and linear reference-based approaches. This increase in mapped reads can enrich downstream variant calling and associative analyses based on SVs from the graph, as demonstrated in other species like cucumber, barley, and vine (Li *et al*., 2022*b*; Cochetel *et al*., 2023; Jayakodi *et al*., 2024). Overall, these results confirm that graph-based representations are more capable at capturing large insertions or deletions that can be missed by linear-reference pipelines, and confirm similar observations made in tomato (Alonge *et al*., 2020) or in maize (Hufford *et al*., 2021) pangenomes. These gains come nonetheless with substantial supplementary CPU costs (Suppl File S1, section 11), a point on which graph-based methods should improve.

Interestingly, we observed diminishing mapping gains for samples based on higher-depth sequencing – but still gains though. While this appears counterintuitive at first, the general sampling principles behind genome skimming approaches (Straub *et al*., 2012) may be an explanation for this observation. Briefly, genome skimming is used when sequencing capacity per sample is low (few reads) and the information held by genomic elements with high copy numbers are sufficient for the targeted question (taxonomy or populational signals). The repetitiveness of these elements in the genome tends to increase their chance to be randomly sampled in a non-targeted approach, and this sampling bias is consolidated when an assembly strategy is used (Straub *et al*., 2012). As a consequence, low depth sequencing tends to extract a large proportion of repeated elements such as organelles, nuclear rRNA tandem repeats or different families of homologous repeats such as transposable elements. By making the hypothesis that our low coverage samples follows this trend (the large proportions of transposable elements that we identified from our sample tend to support this hypothesis), it is expected that a larger diversity of reads could be mapped to the graph and not the reference genome, as the graph should better represent the diversity of apricot repeats. These observations also raise a warning for posterior association studies : they could be biassed towards certain genomic elements, such as satellites and transposons in the samples of lowest depths (Ou *et al*., 2019).

## Conclusion

Our comprehensive analysis underscores the critical importance of broad genetic sampling and meticulous structural characterization for a complete understanding of genomic diversity and evolution in general and in the *Armeniaca* section in particular. Future studies integrating additional diverse genotypes, particularly through careful selection of accessions aiming at filling identified diversity gaps, and incorporating phenotyping for associative analysis, will greatly benefit from the construction of this super-pangenome. Graph-based SV datasets have demonstrated strong utility for GWAS in crops, including barley (Jayakodi *et al*., 2024), highlighting their potential for unlocking previously unexplored genotype-phenotype associations missed by standard SNP-based analysis. The dataset built in this study will enable us to fully leverage the observed genetic diversity for apricot crop improvement.

## Acknowledgments

IB analysed the data and produced the results, with contributions from QTB, JC and BL. AM contributed to results and graph statistics, by giving early access to the tool Pan1c. JC contributed to transposon-related analyses. QTB, SD, AC and AG contributed to de-novo assemblies. LD, BL & VD managed the project. IB, QTB, JC, LD, VD and BL wrote the manuscript. We also thank Elise Maigne for her help on R code related to some figures. All authors read and validated the manuscript.

IB acknowledges a PhD fellowship from MNSER (ministère de la recherche française) and support from the University of Bordeaux (Ecole Doctorale 154). SD was supported by an apprenticeship grant from INRAE. This work is funded by the Horizon Europe FRUITDIV project (#101133964). This work was also supported by state funding managed by the French National Research Agency under the France 2030 program [grant number ANR-22-PEAE-0005]. De novo genome assemblies of wild Armeniaca were supported by and the University of Bordeaux, Département des Sciences de l’Environnement (WOODYSV project, 2020-2022) and ANR JCJC PLEASURE ANR-21-CE20-0005. We are grateful to the genotoul bioinformatics platform Toulouse Occitanie (Bioinfo Genotoul, https://doi.org/10.15454/1.5572369328961167E12), to the Centre de Bioinformatique de Bordeaux (CBiB) bioinformatic platform for providing computing and storage resources and to the CNRGV (Centre National de Ressources Génomiques Végétales, INRAE Occitanie-Toulouse) for the optical maps. We acknowledge valuable inputs from the Pangenome network of the PEPR Agroécologie et Numérique, AGRODIV (ANR-22-PEAE-0005 - AgroDiv) flagship, as well as the BReIF e-infrastructure (ANR-22-PEAE-00014-BREiF) for useful tools to retrieve and submit meta-data associated to our dataset. We thank the INRAE CATI BARIC and Sébastien Carrère (LIPME), for the supply of the functional annotation pipeline: nextflow fonctionnalAnnotatio, also available on Zenodo (https://zenodo.org/records/7603192).

---

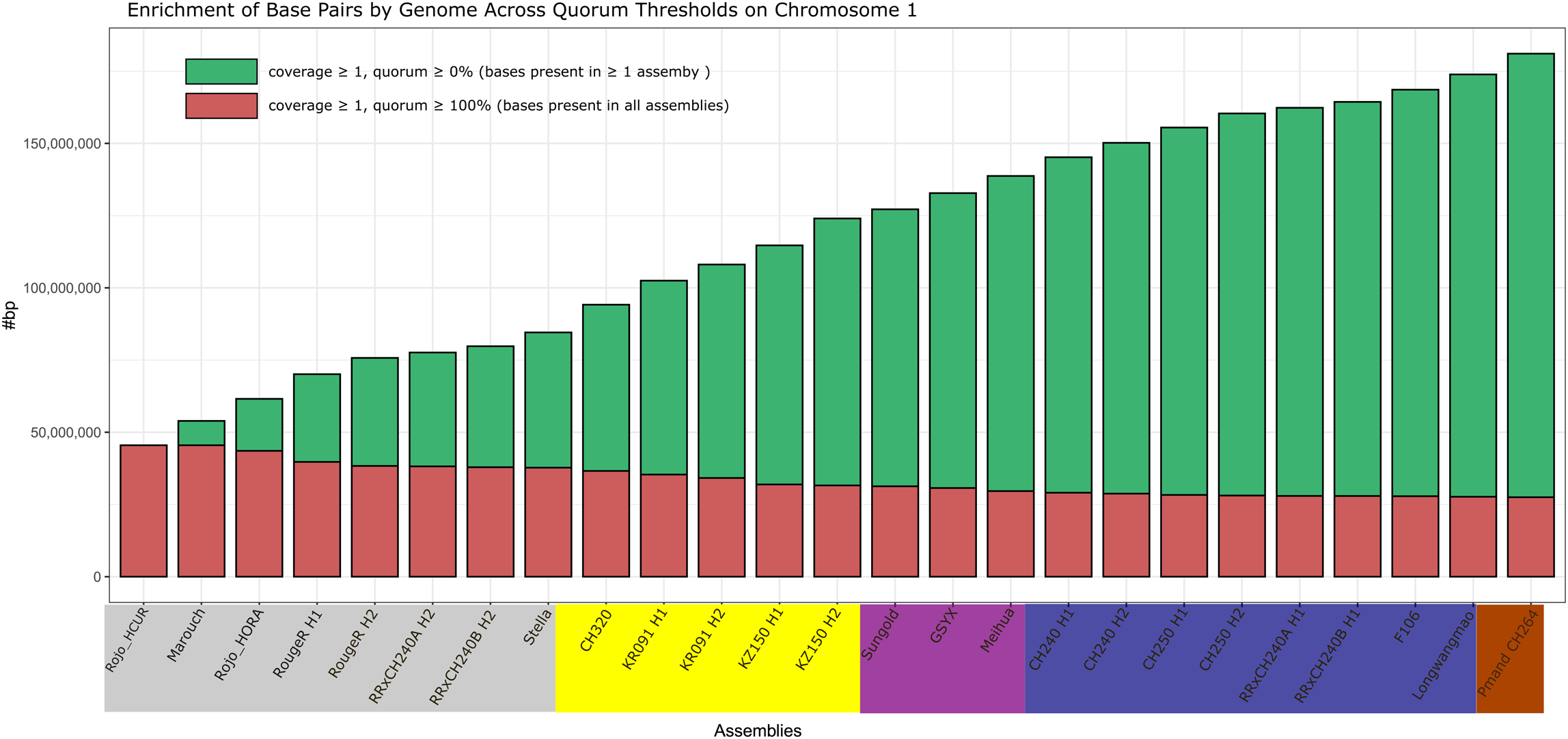

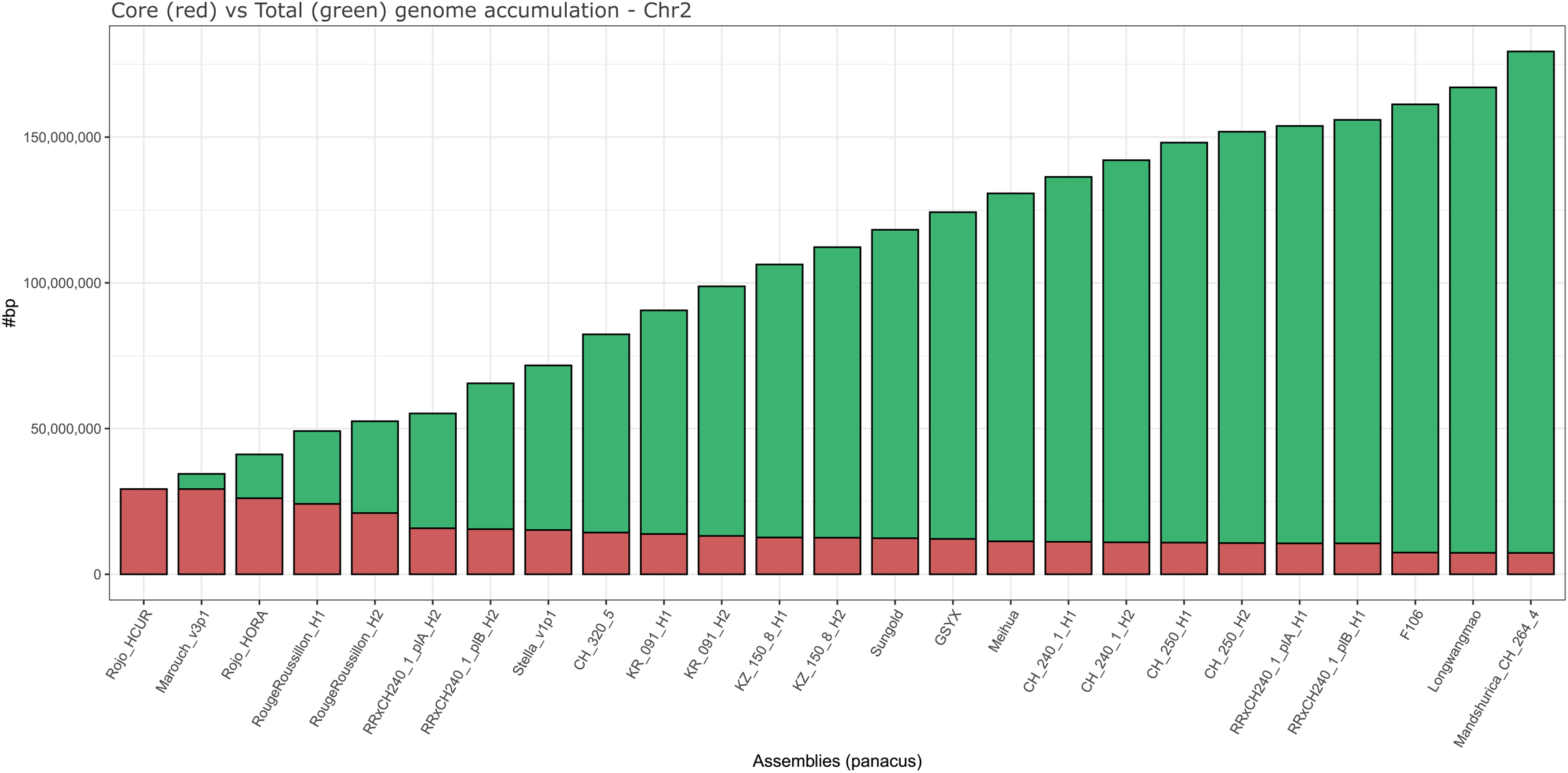

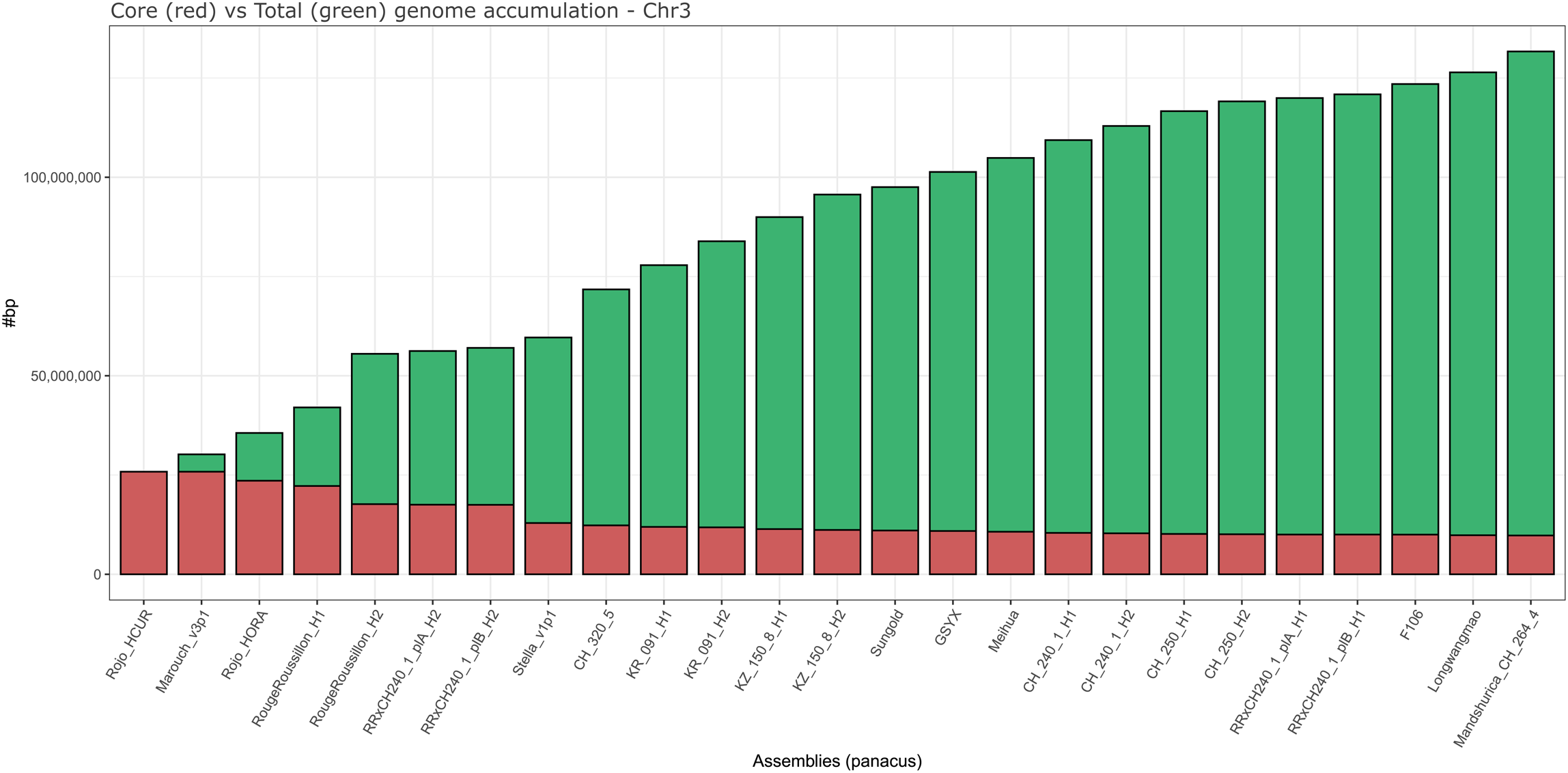

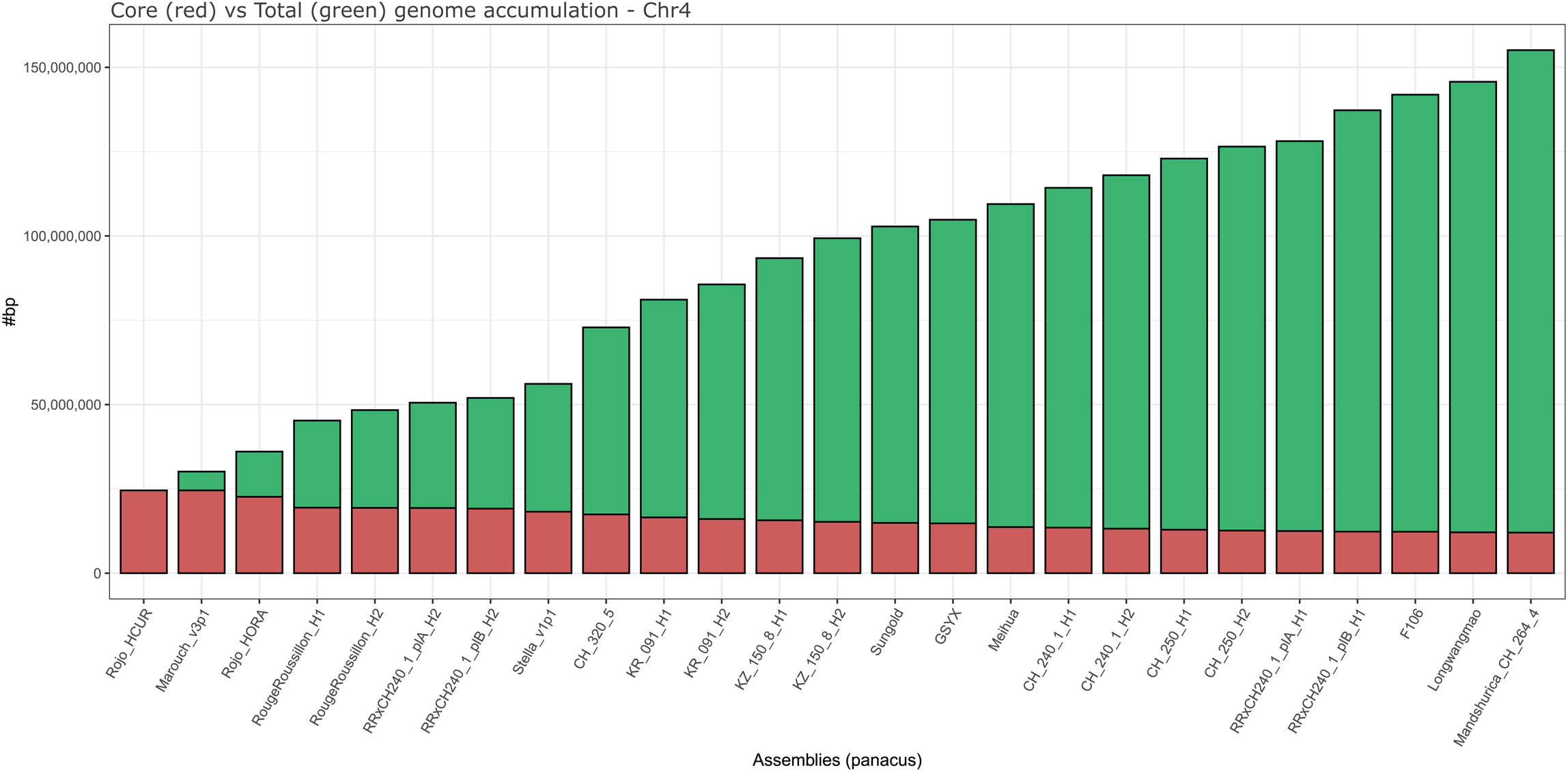

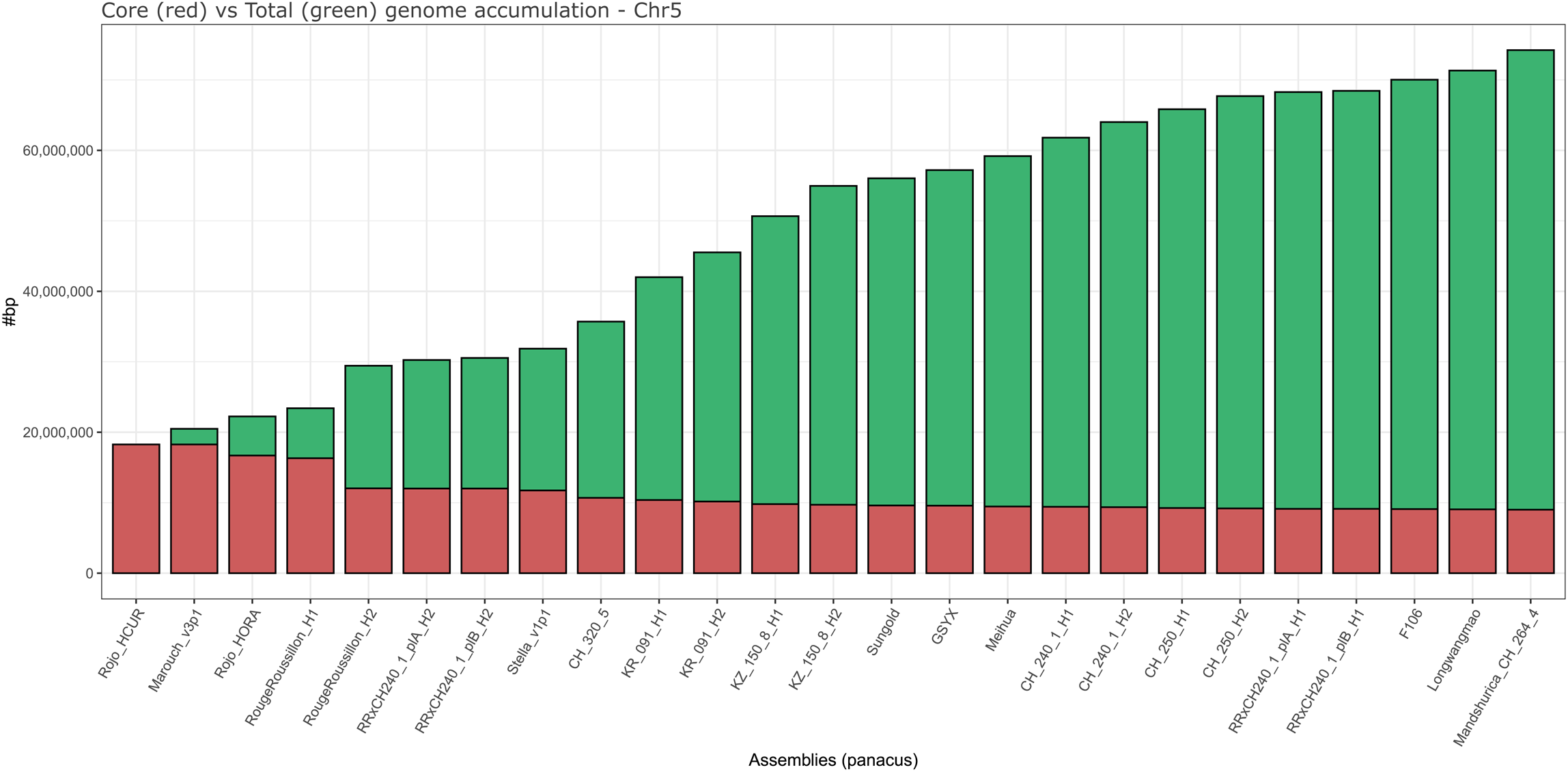

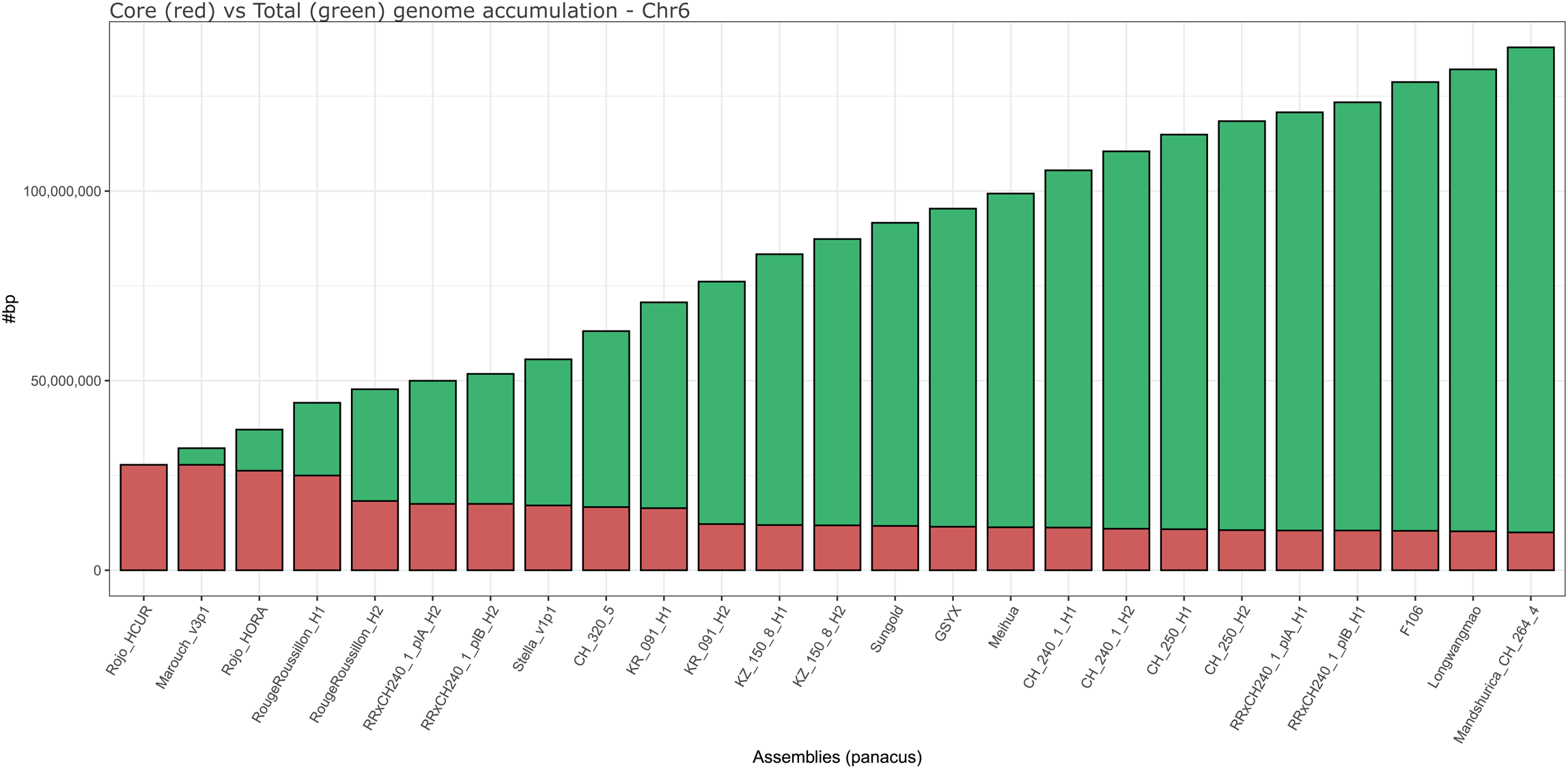

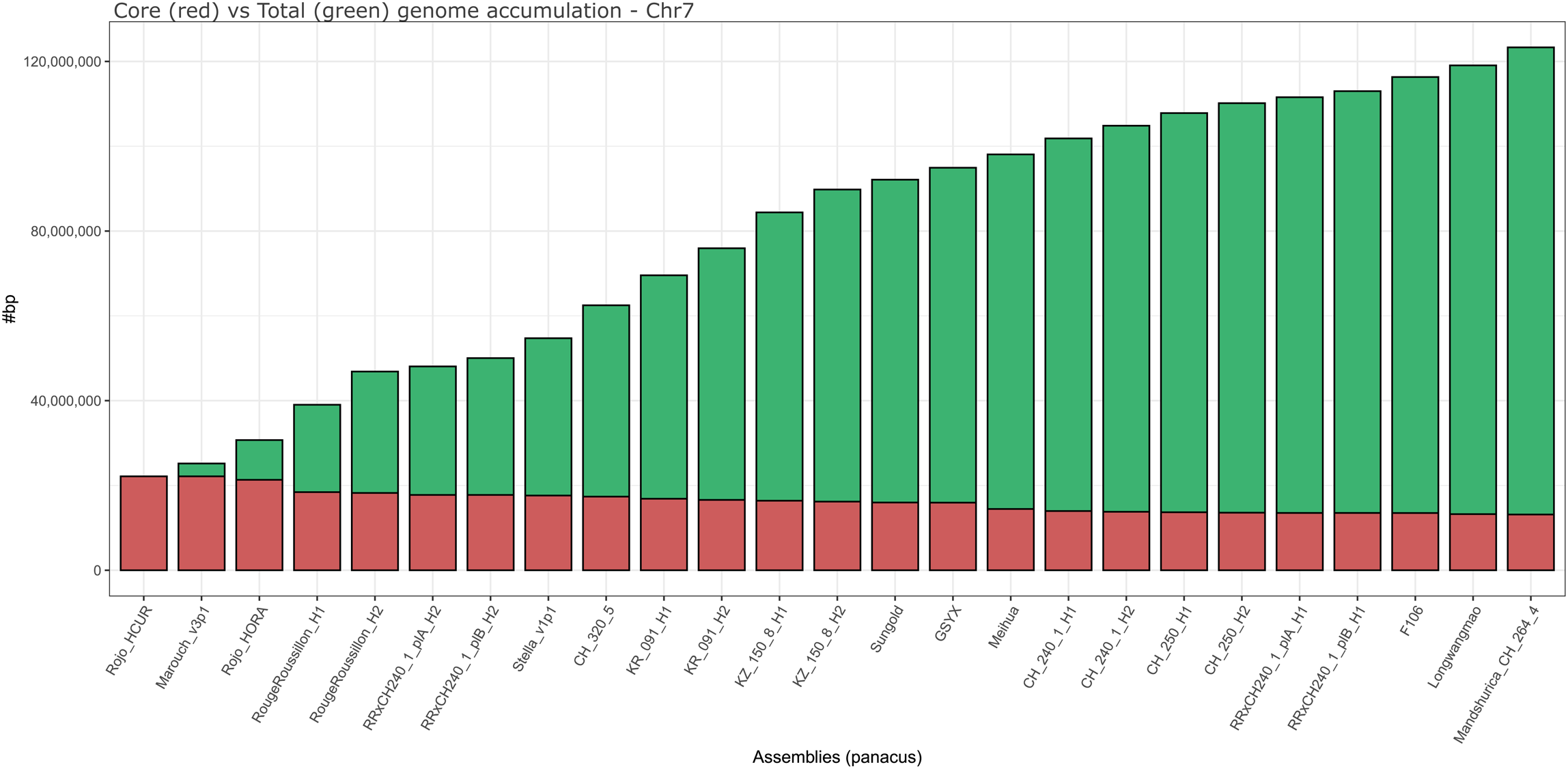

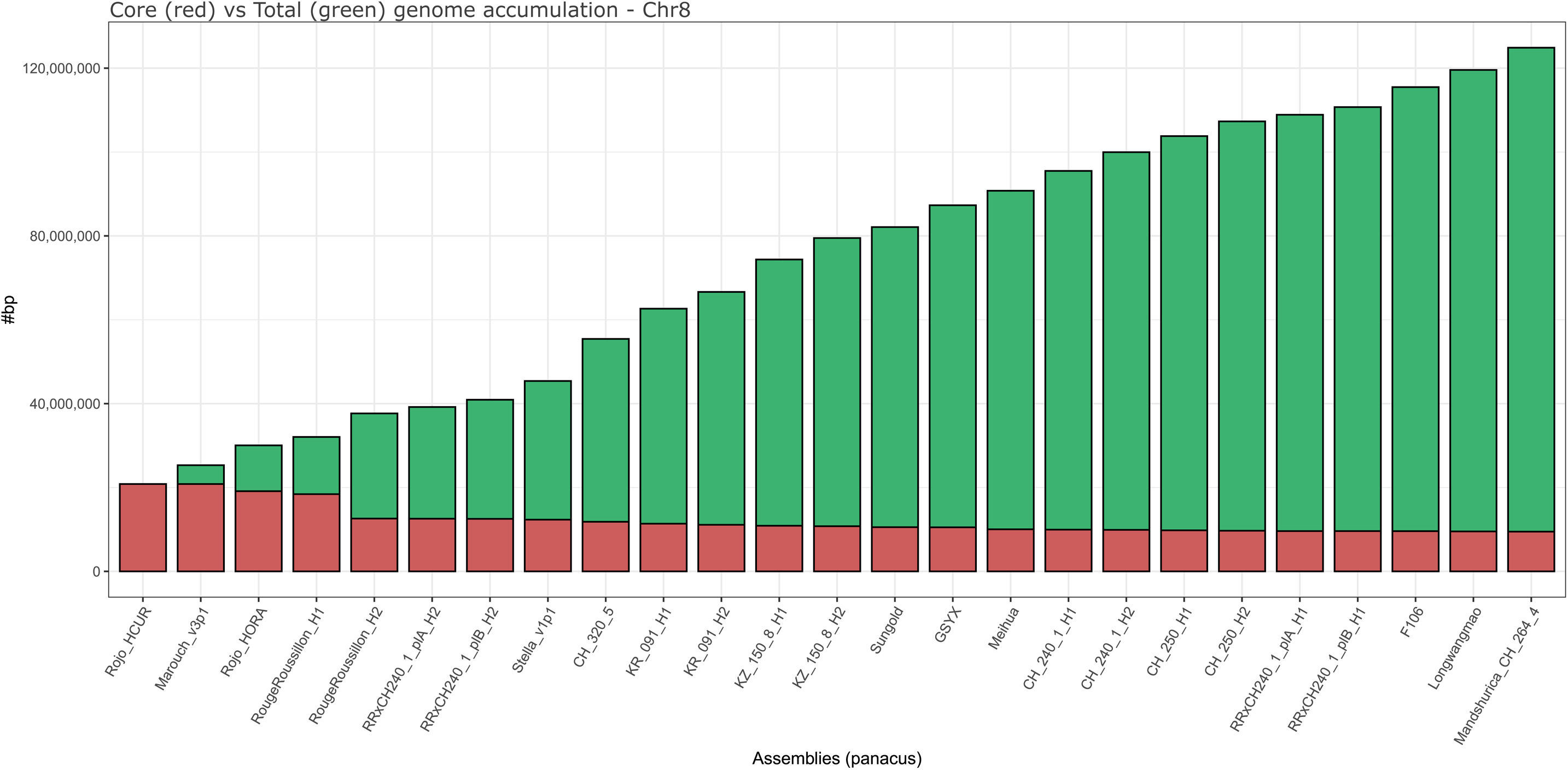

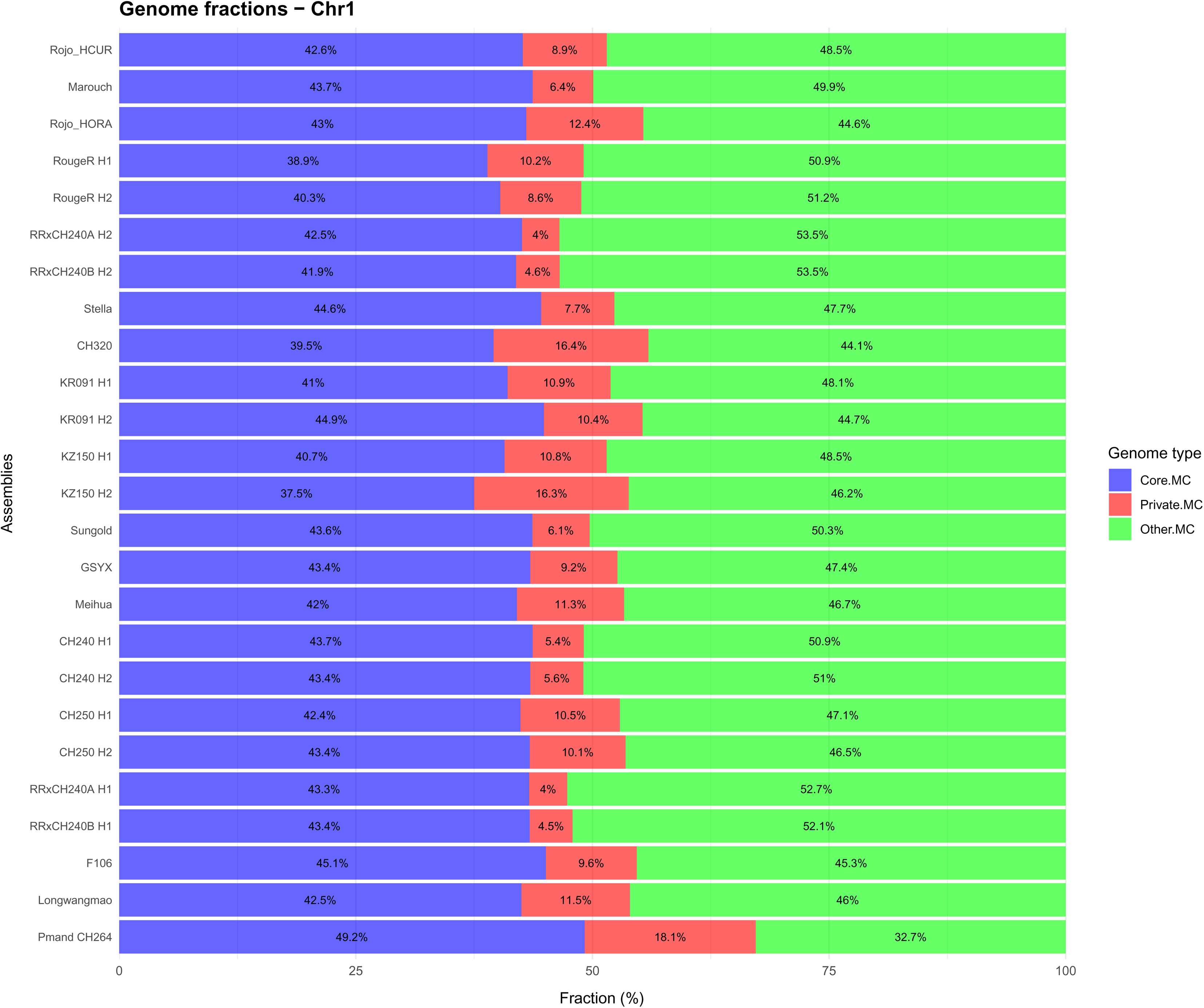

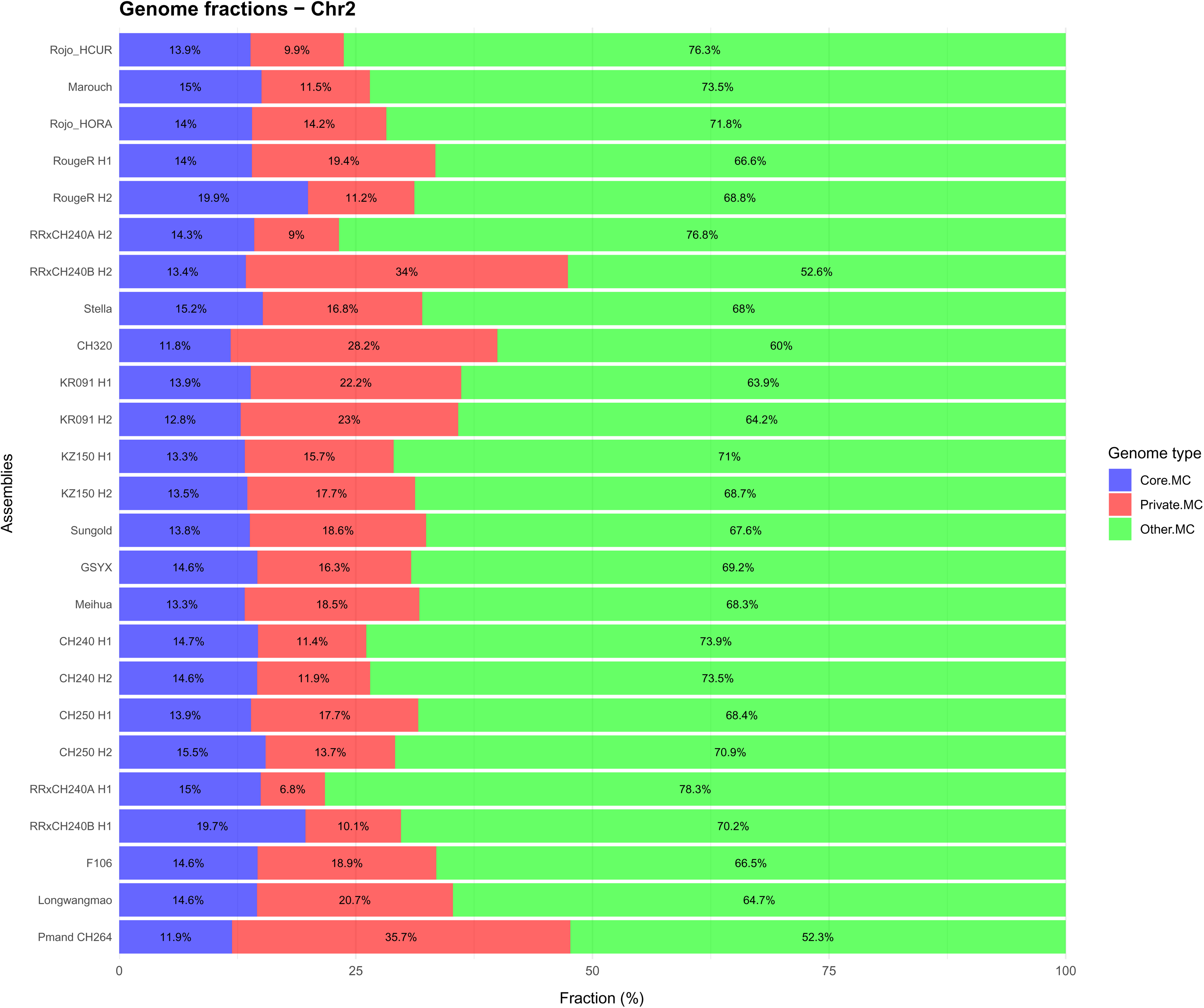

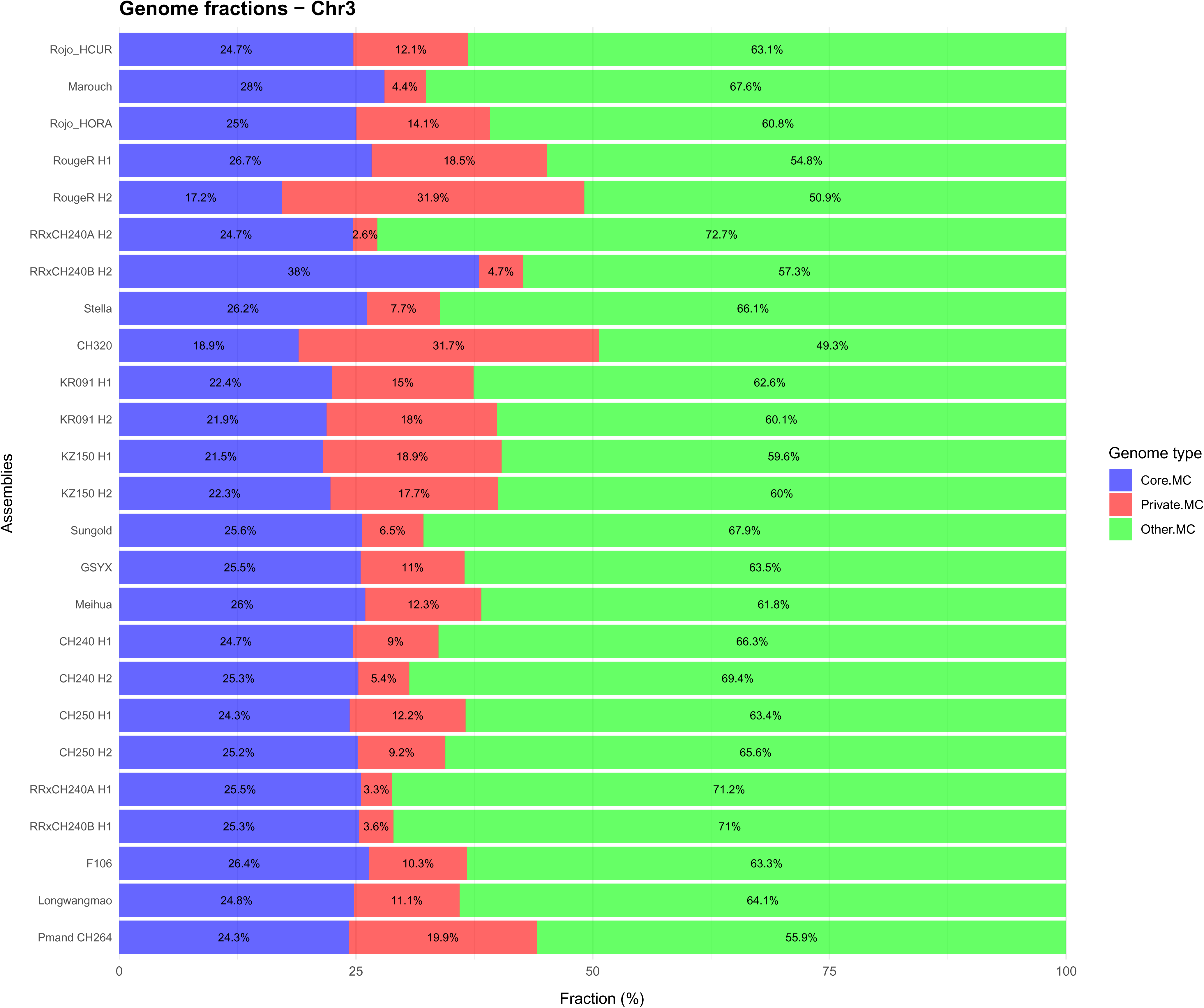

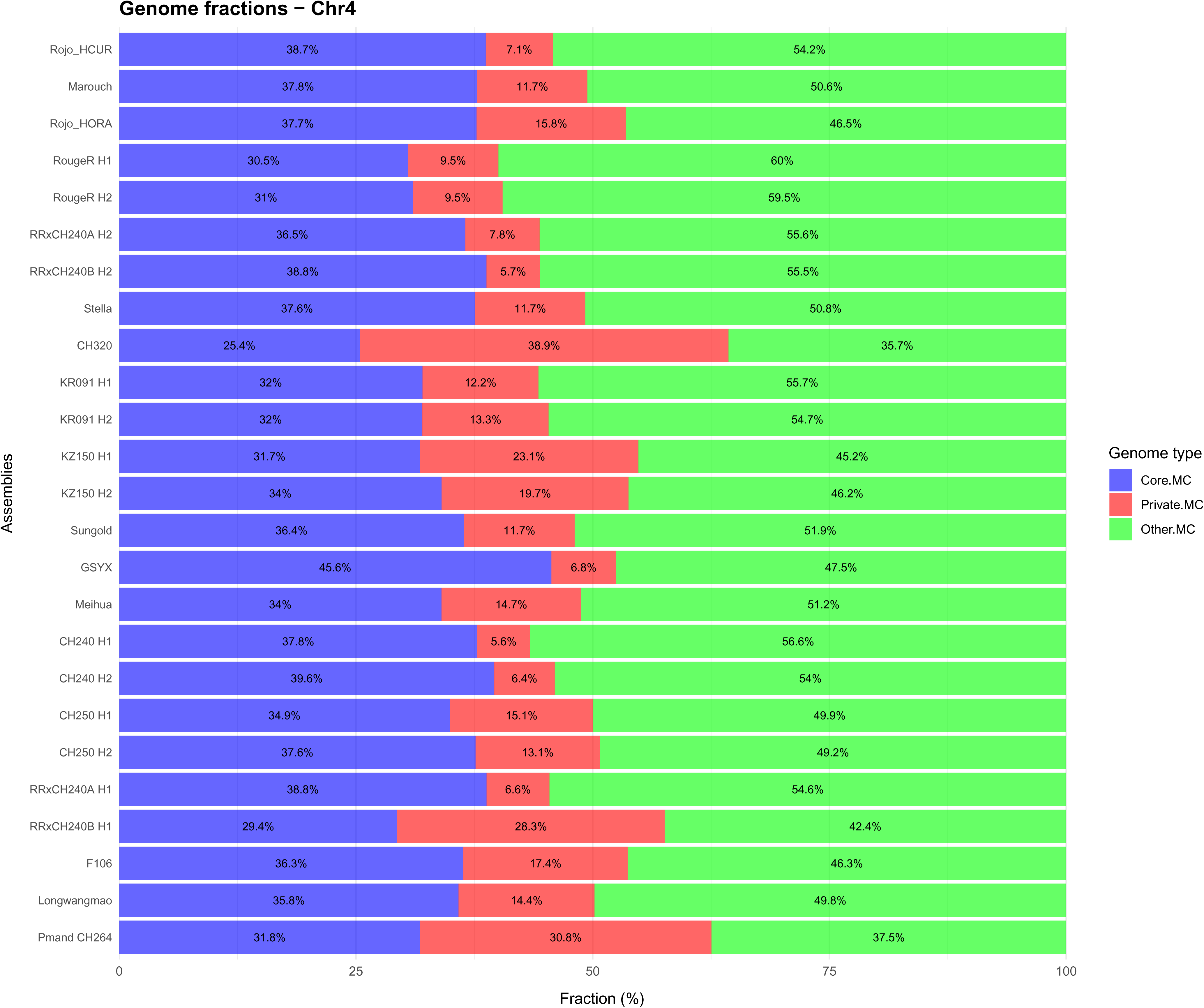

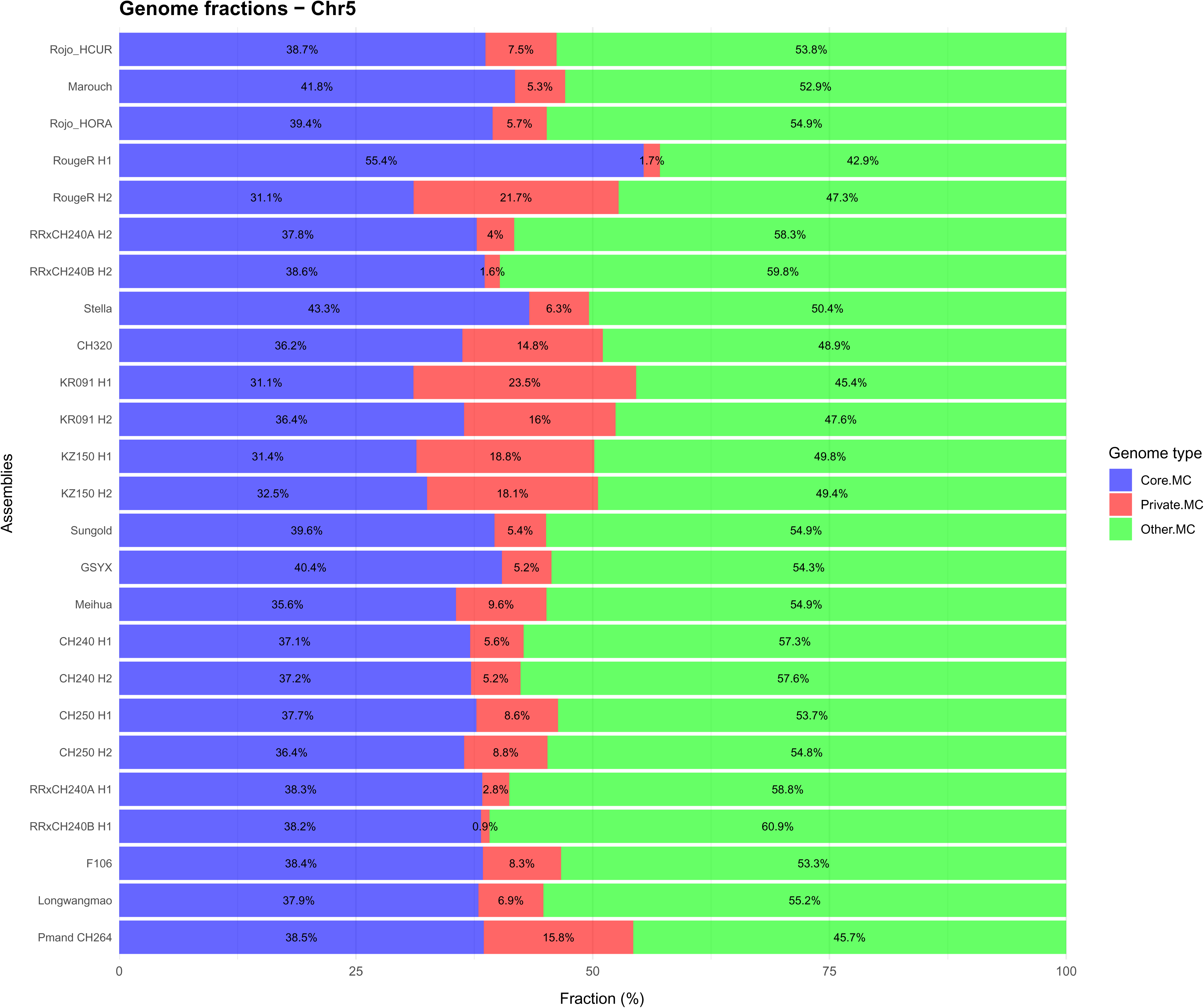

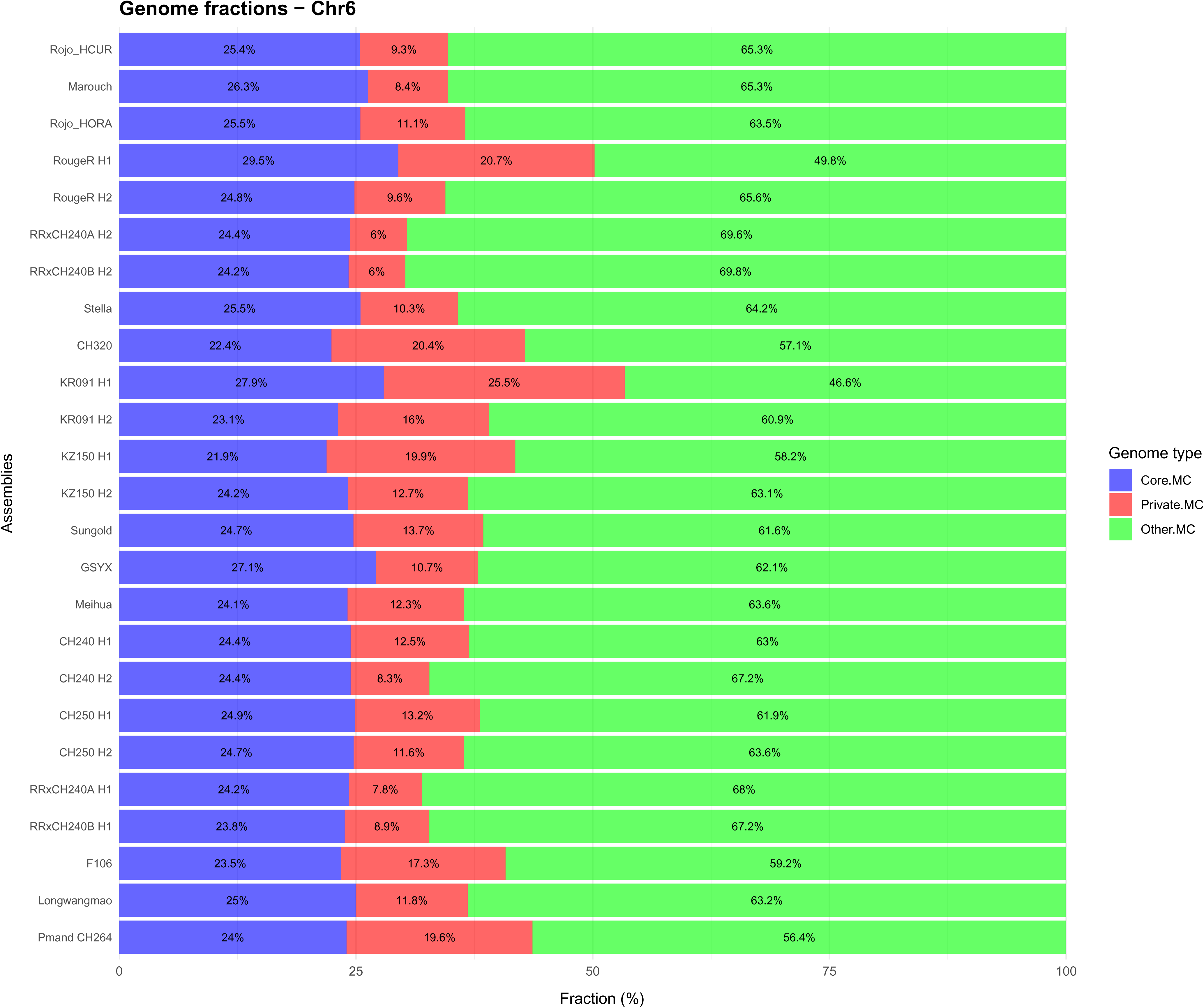

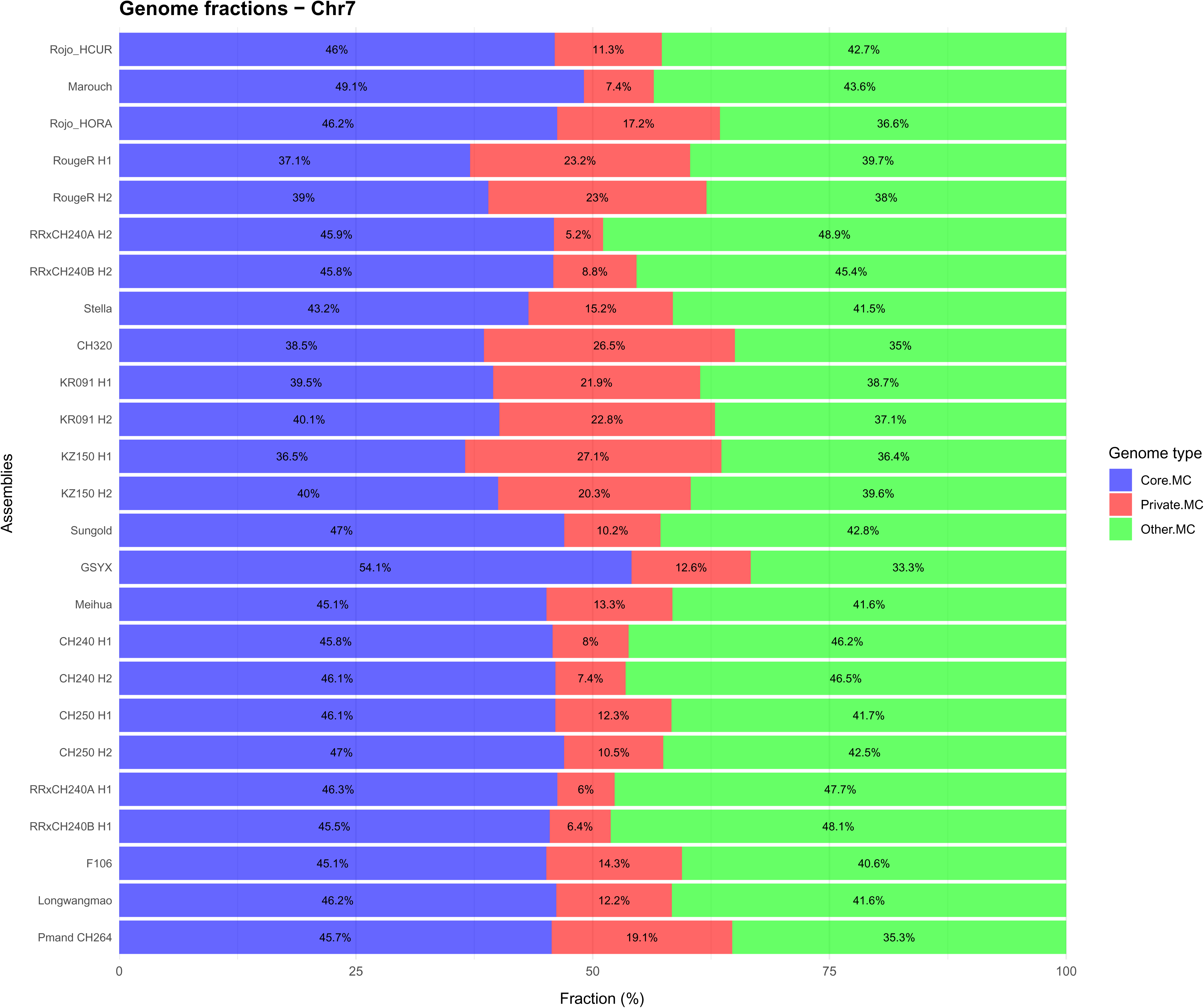

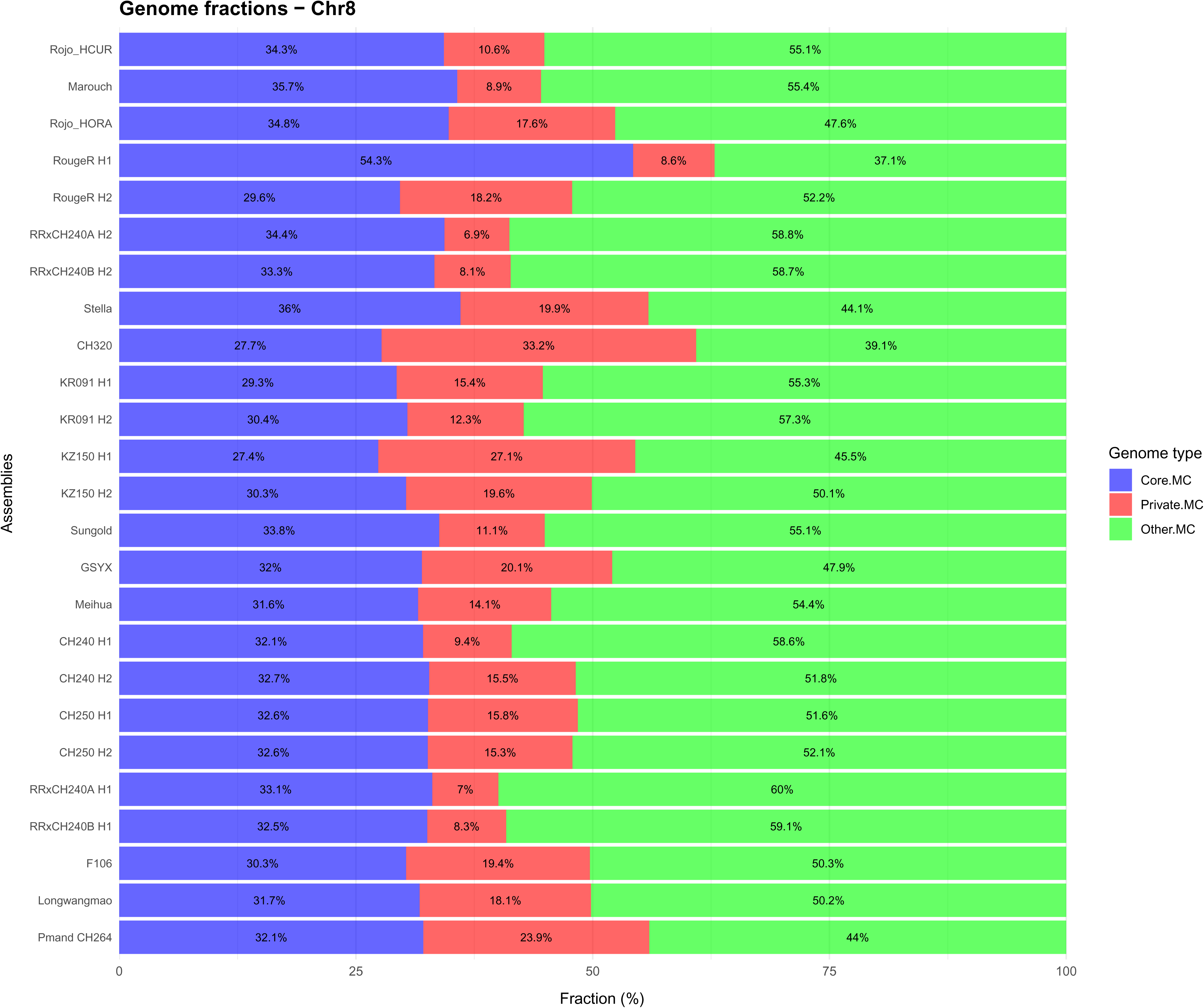

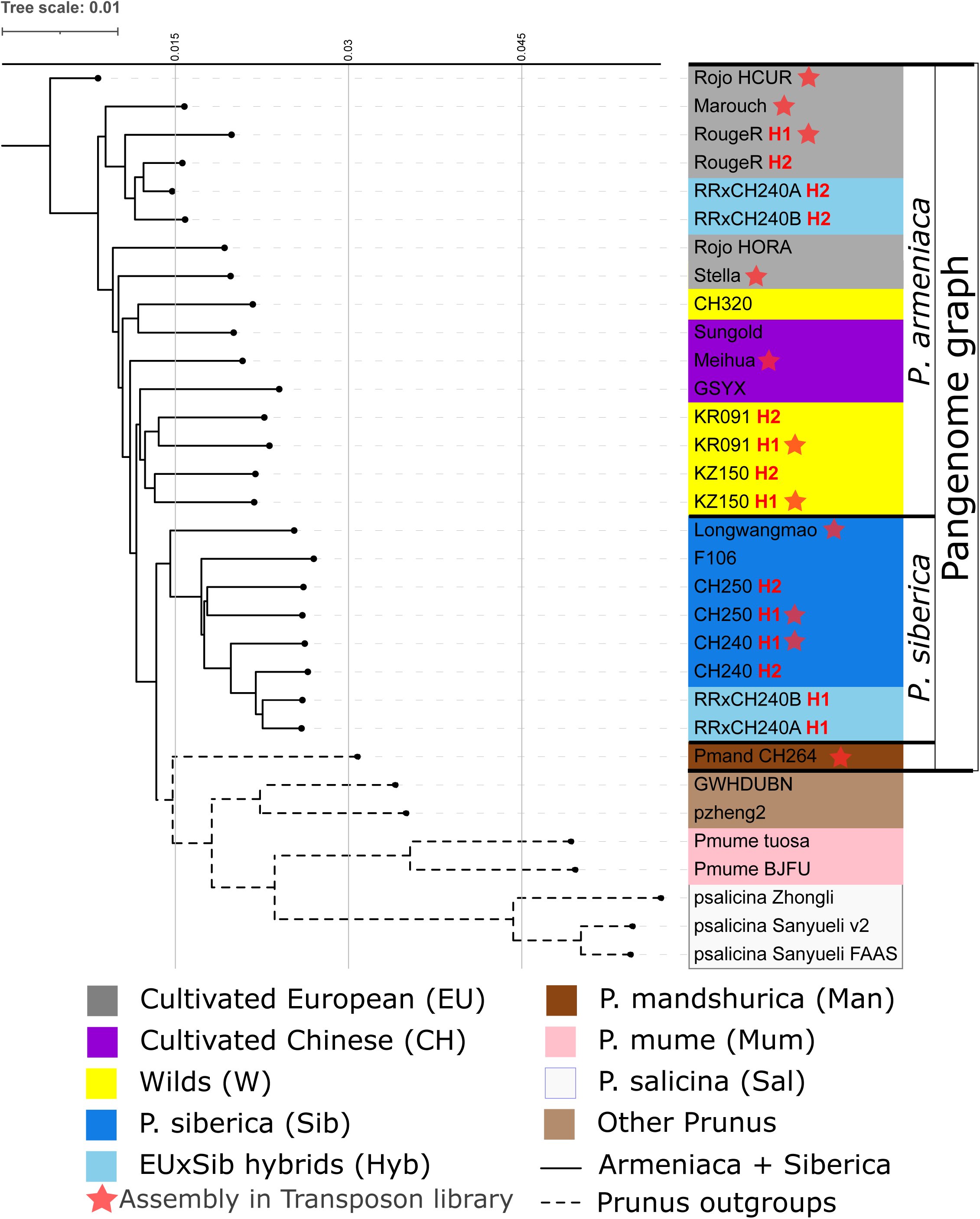

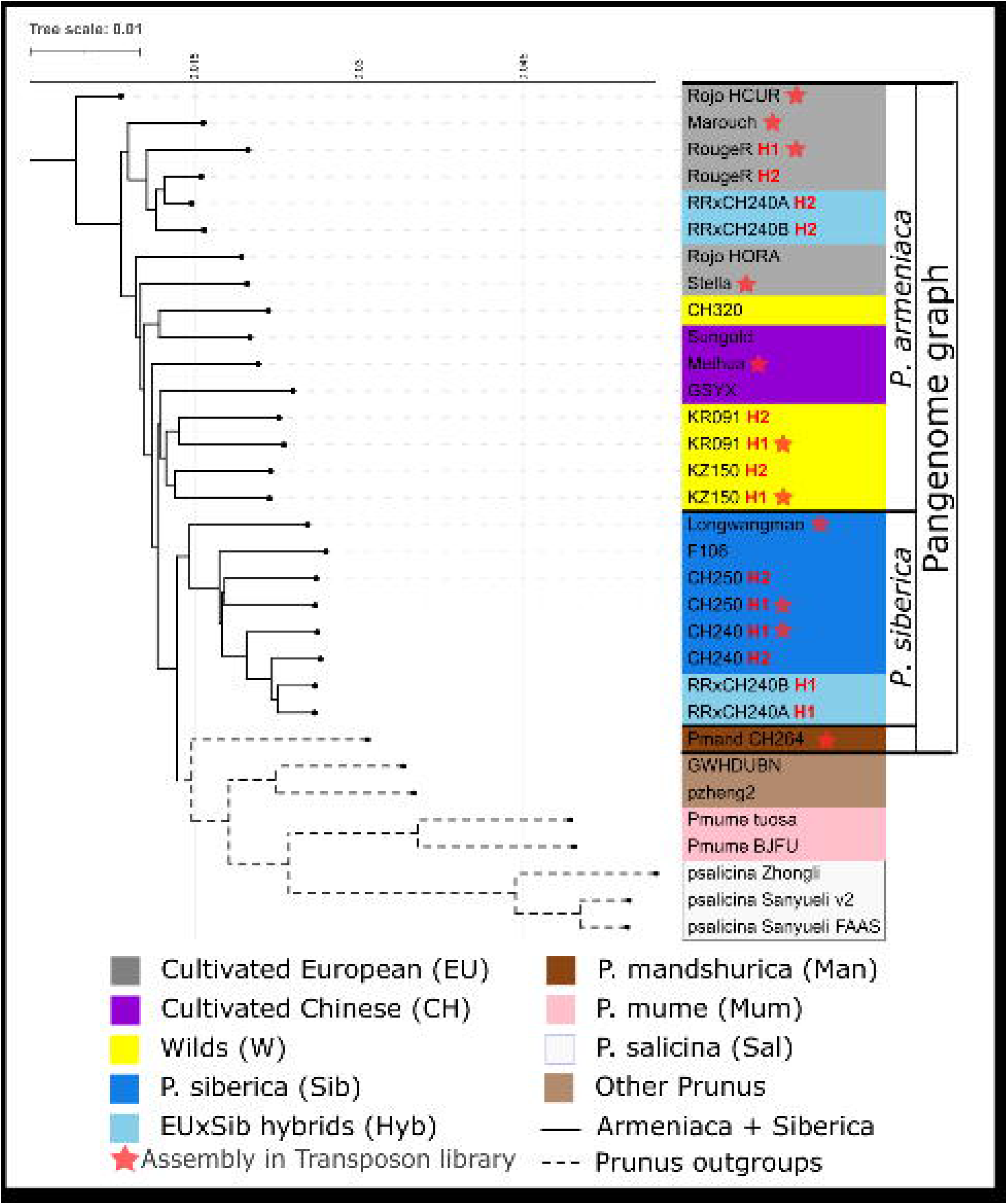

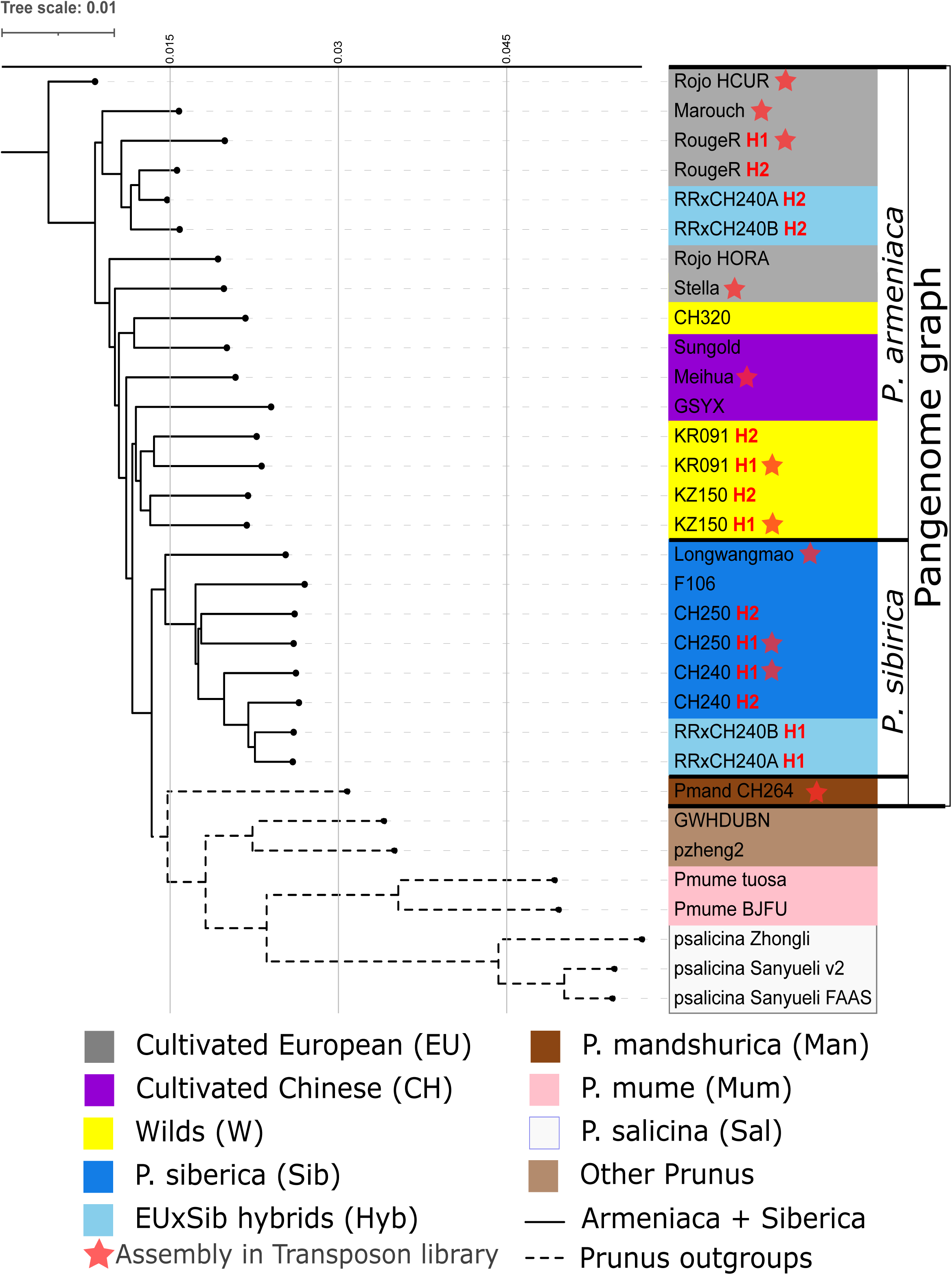

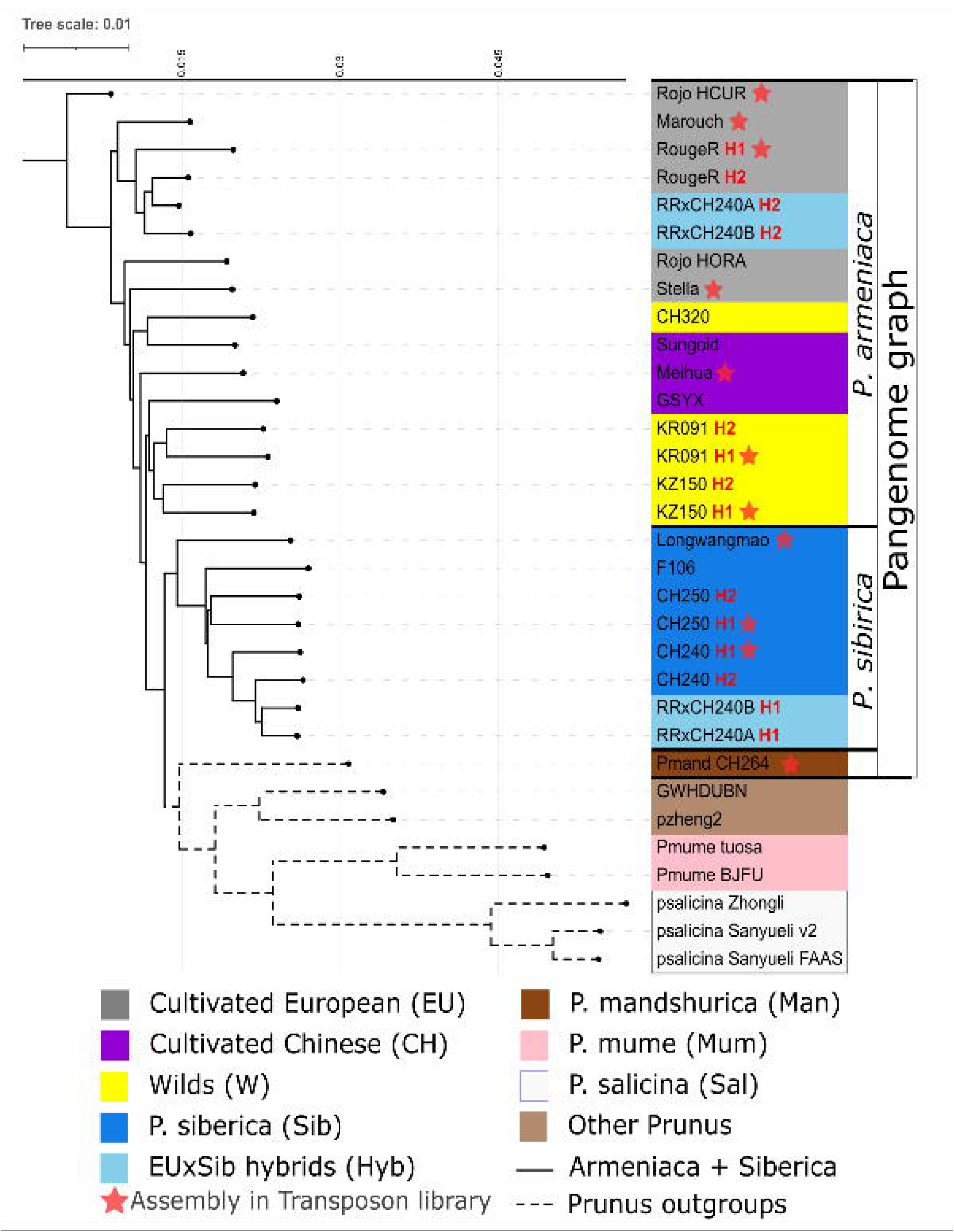

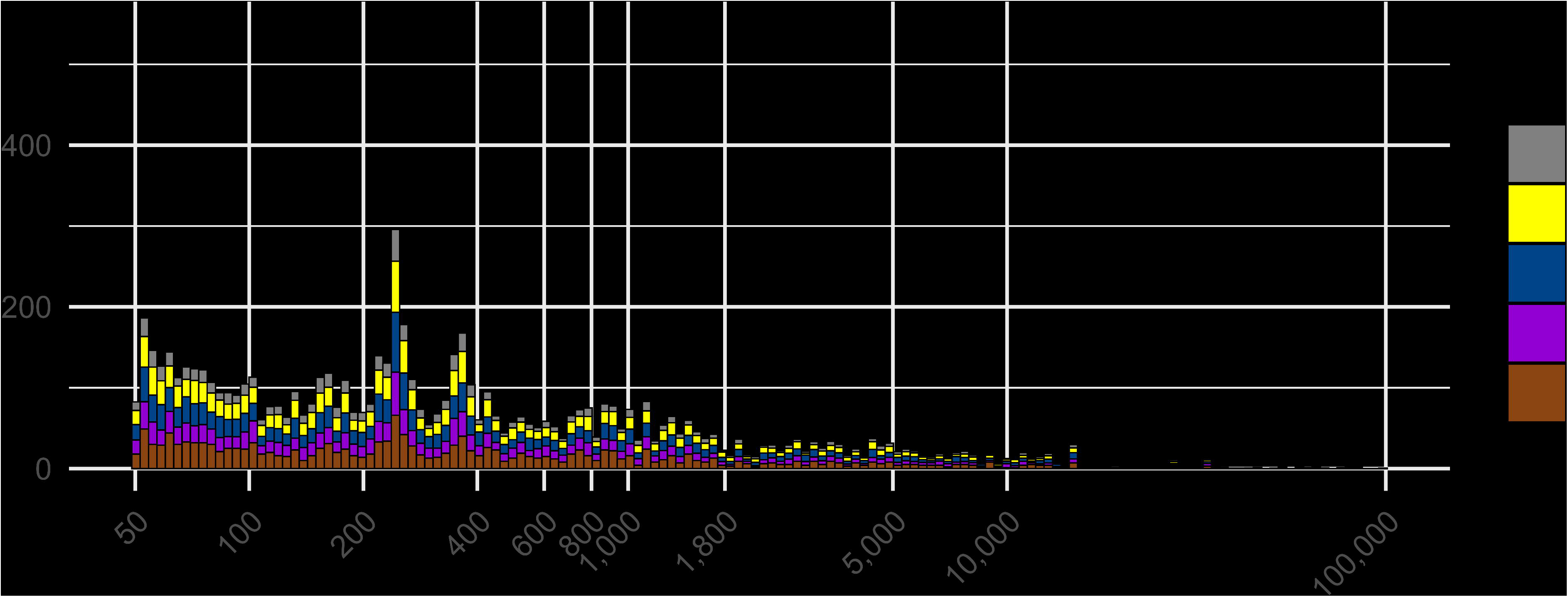

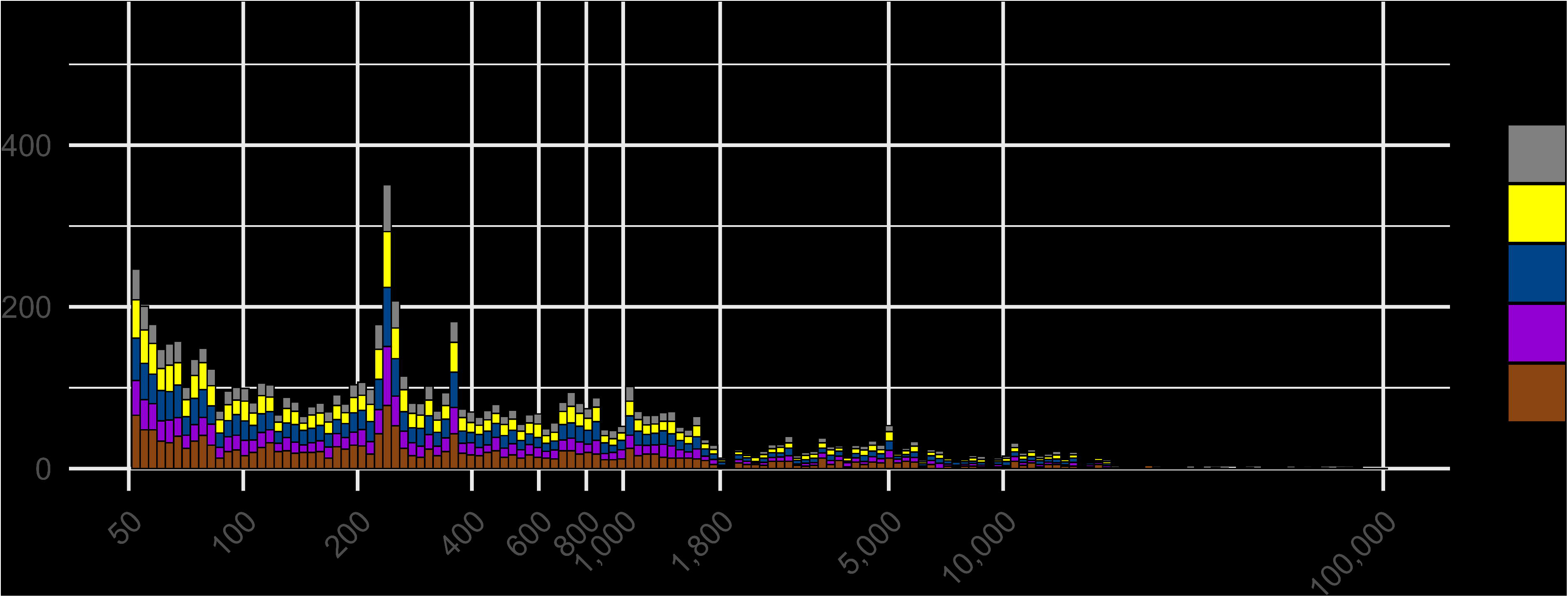

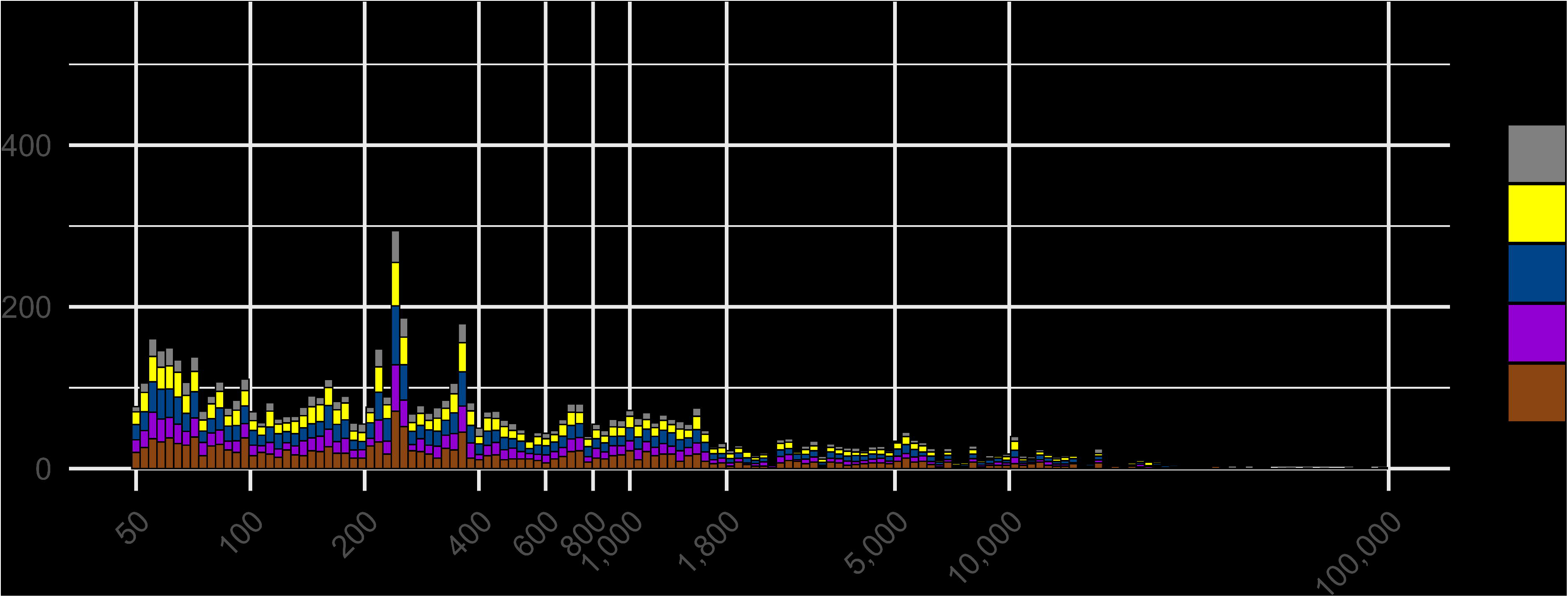

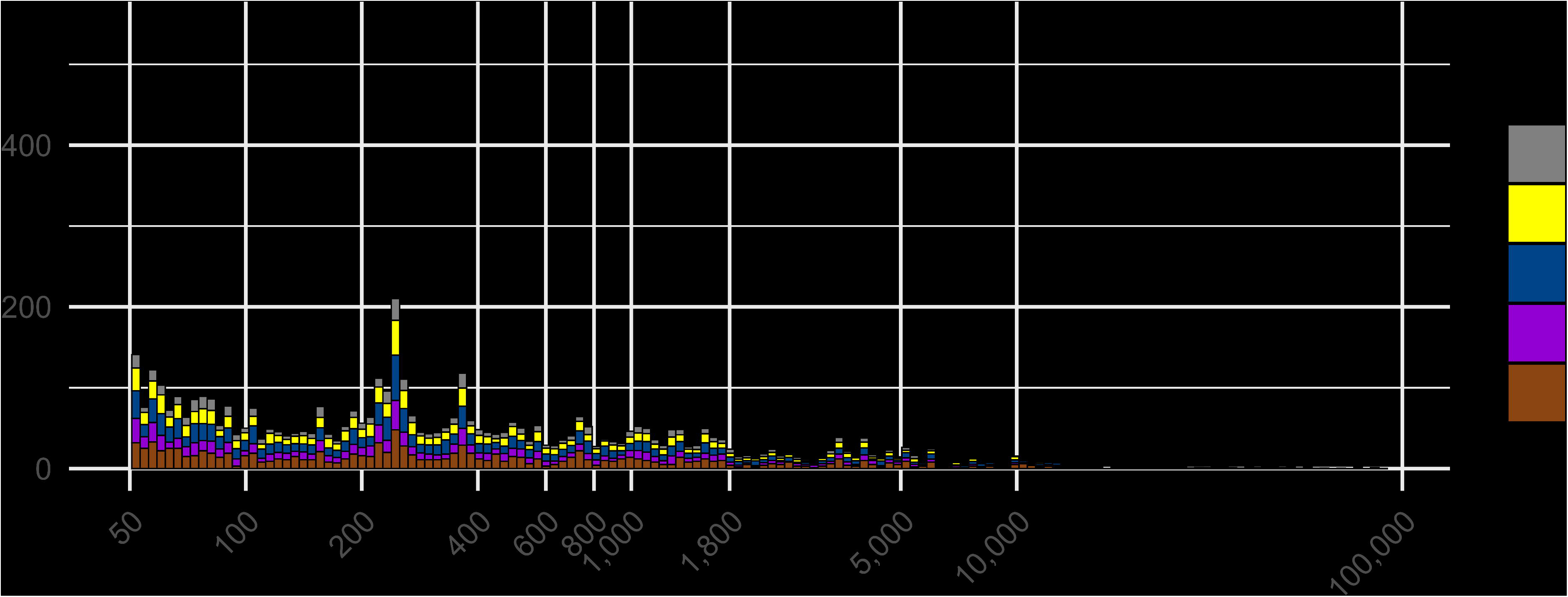

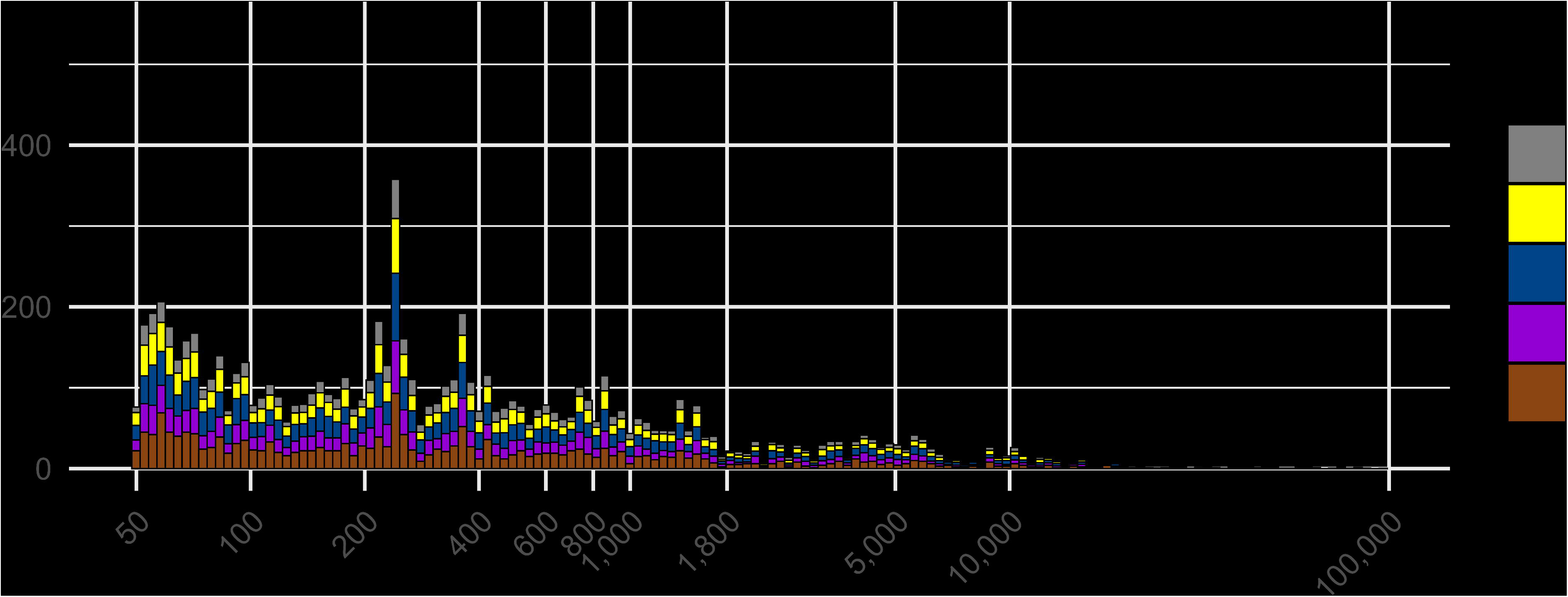

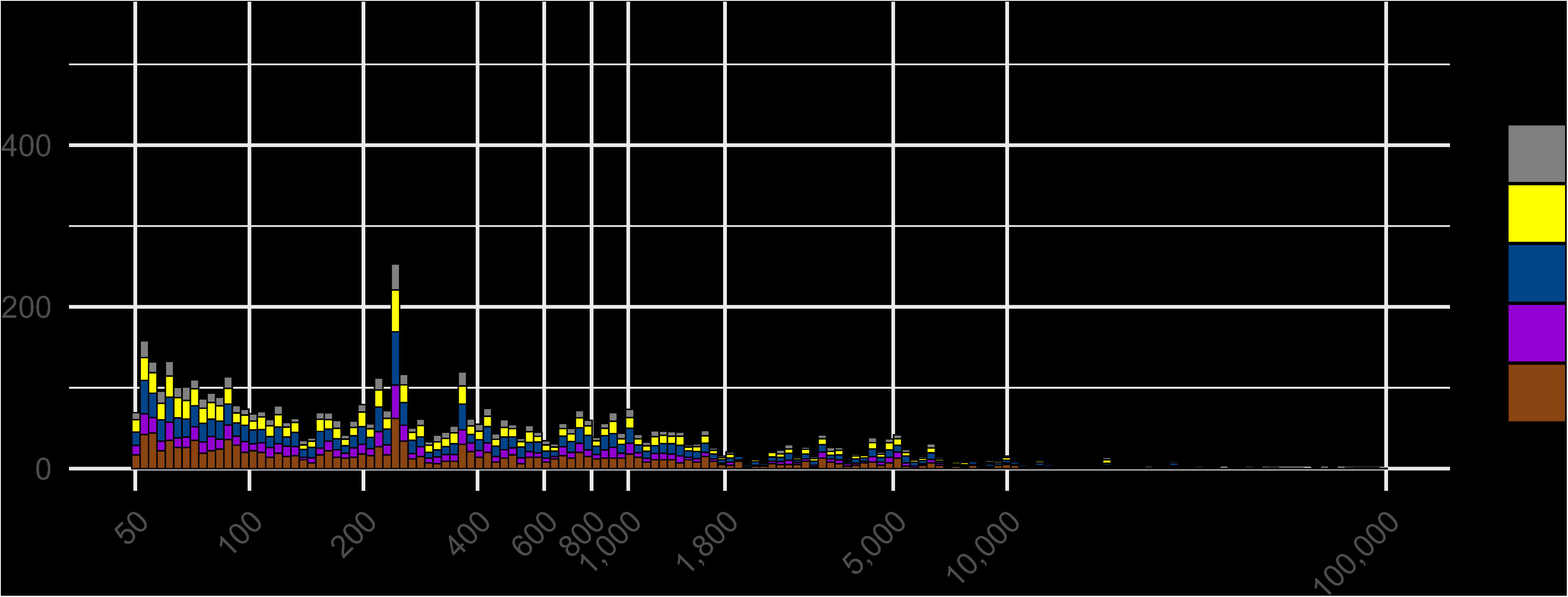

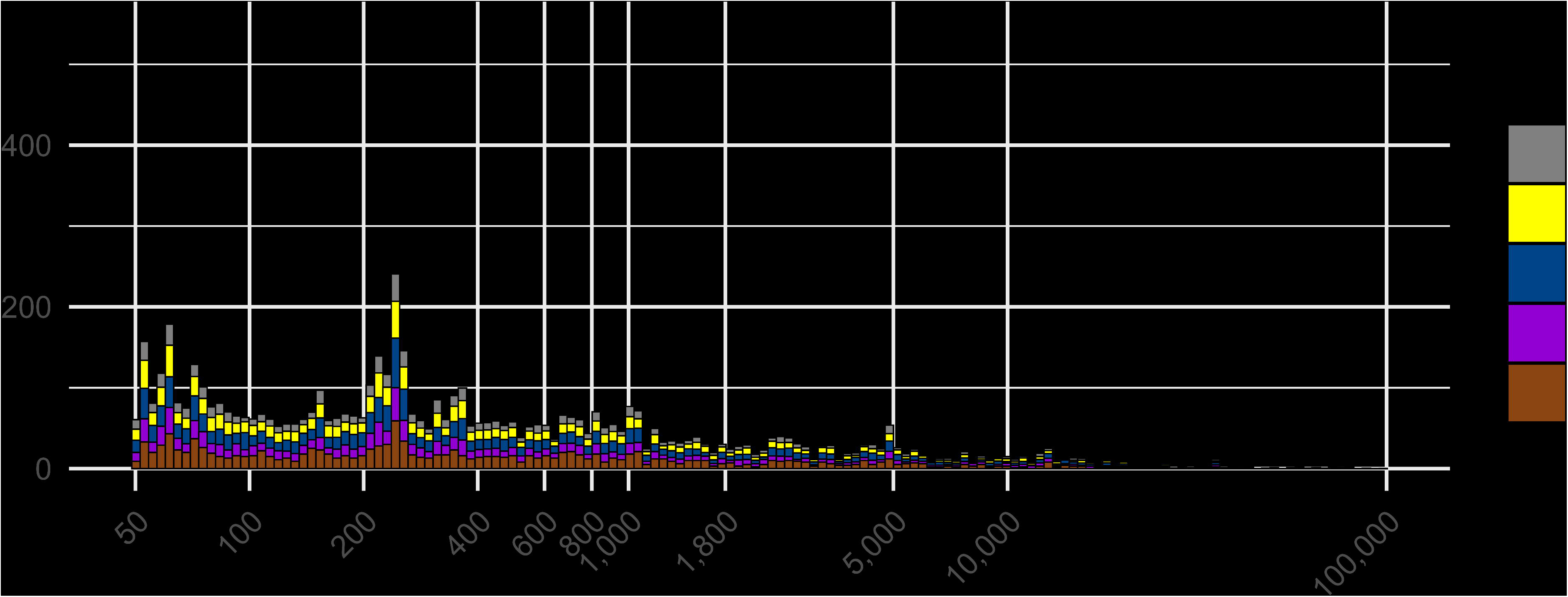

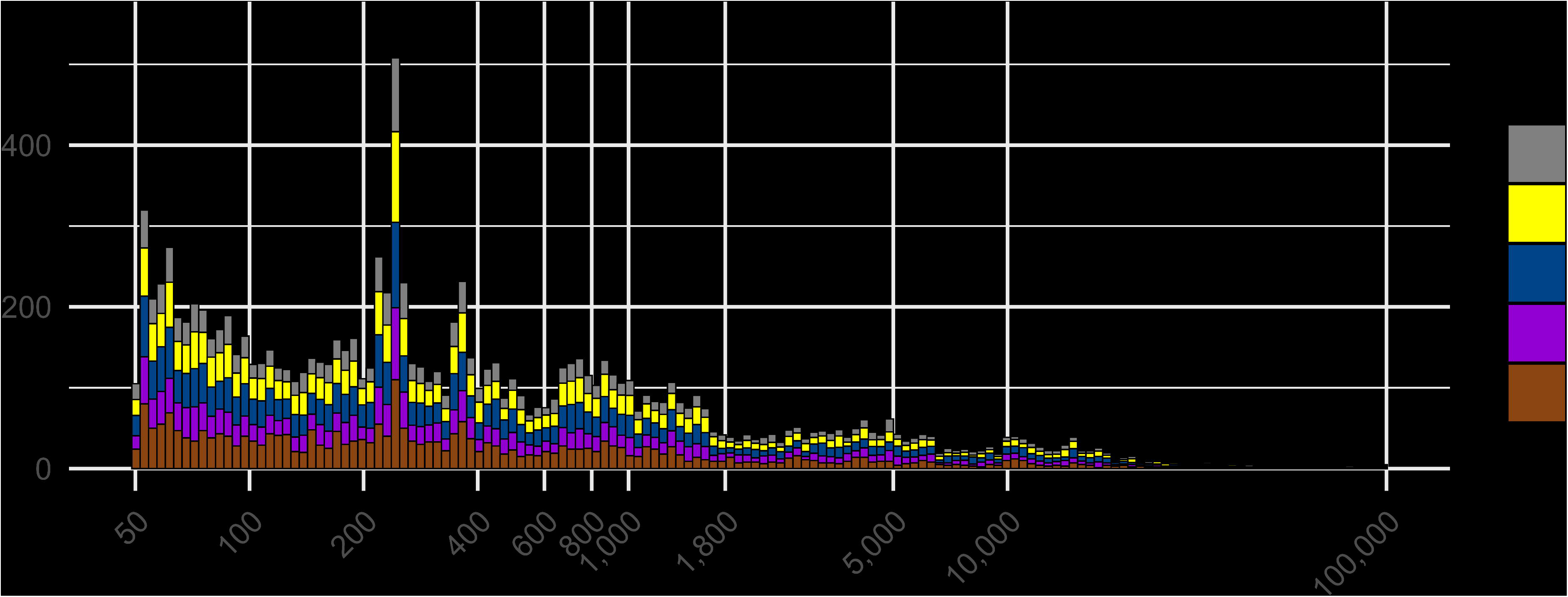

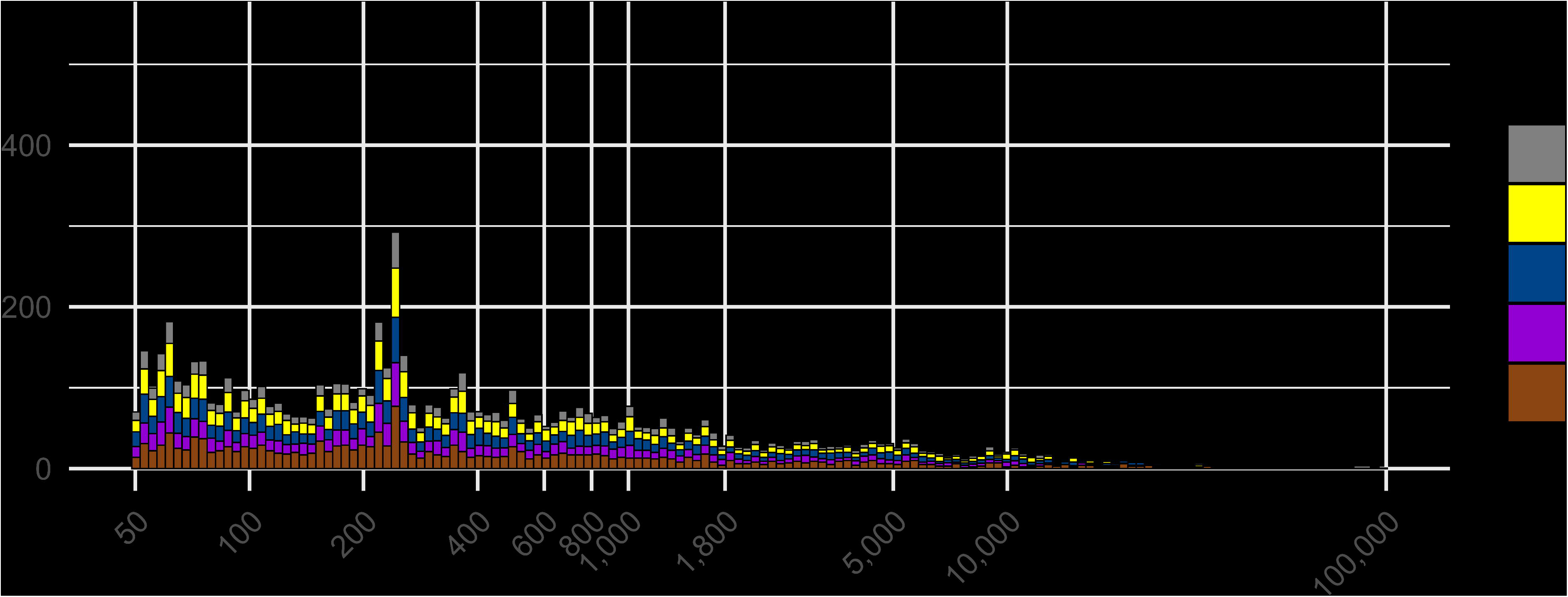

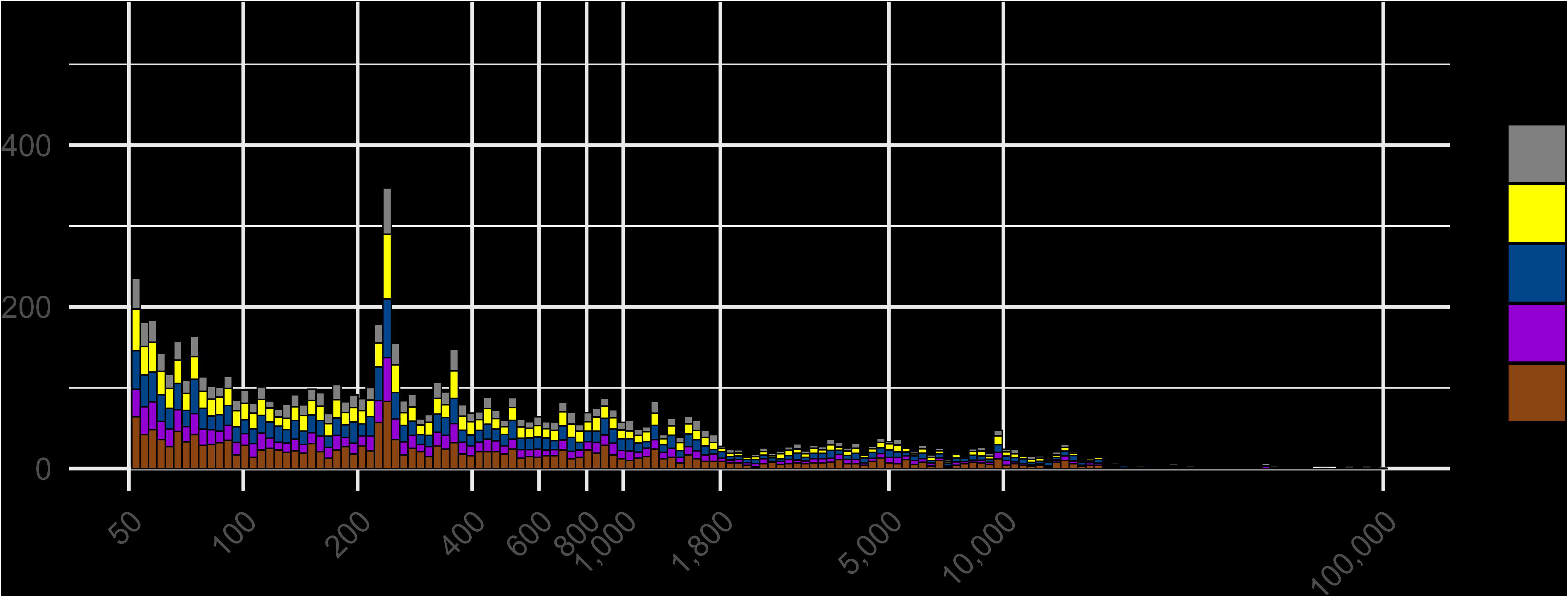

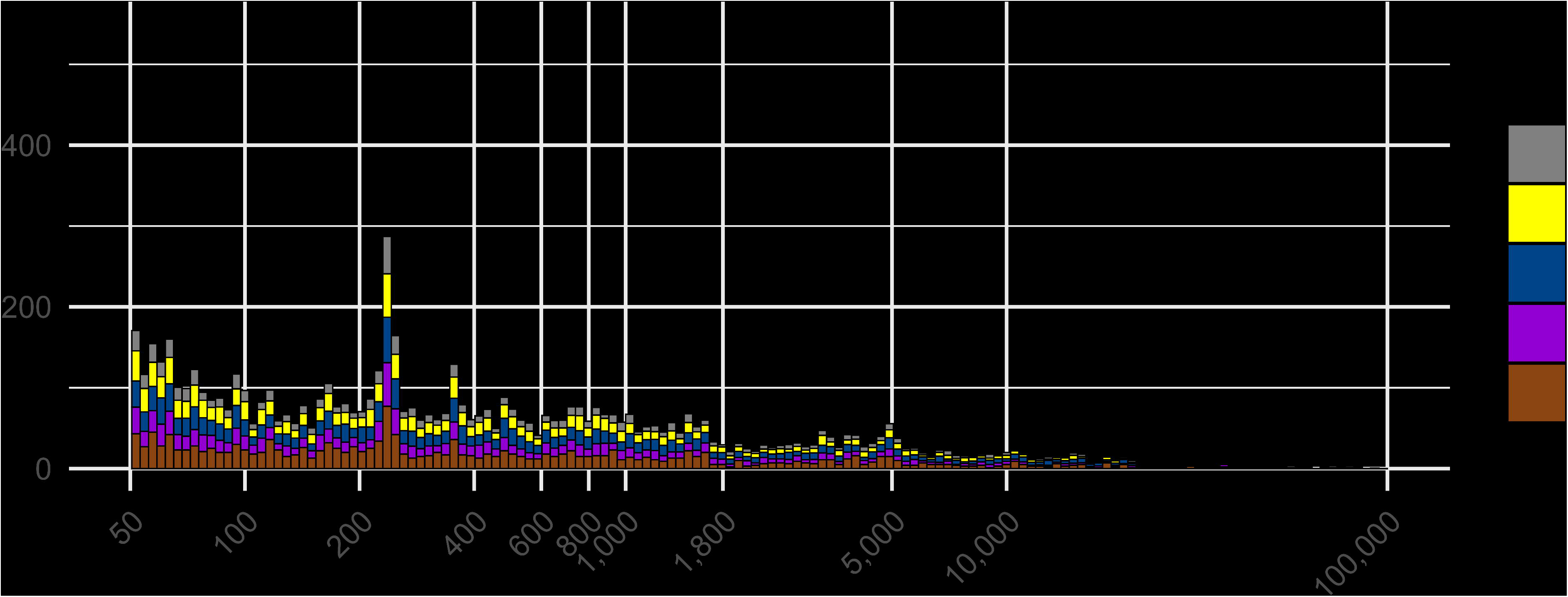

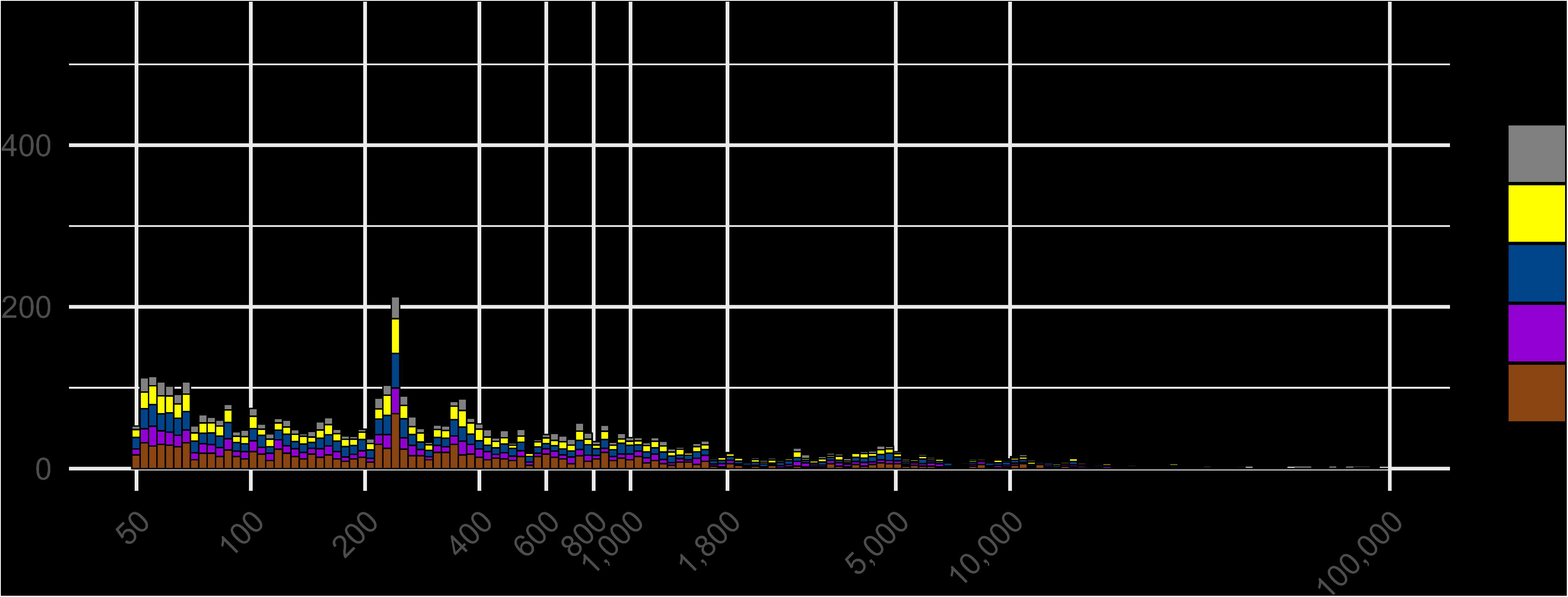

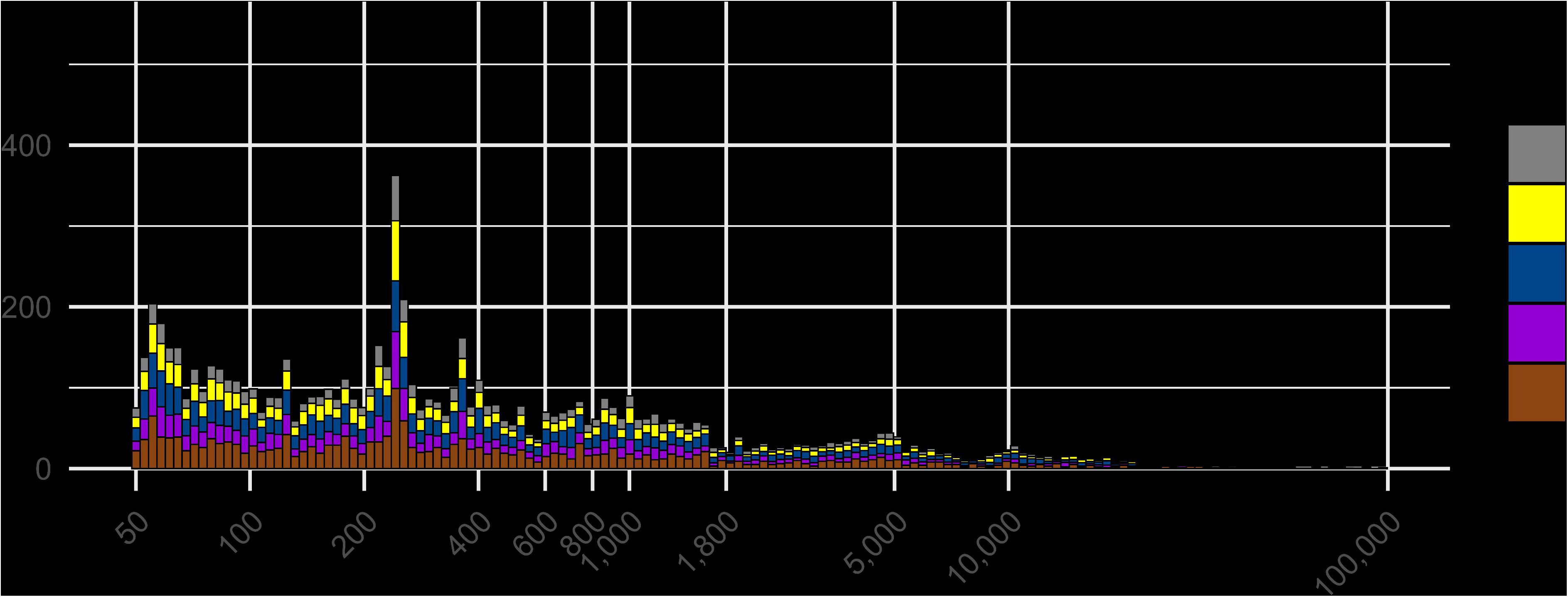

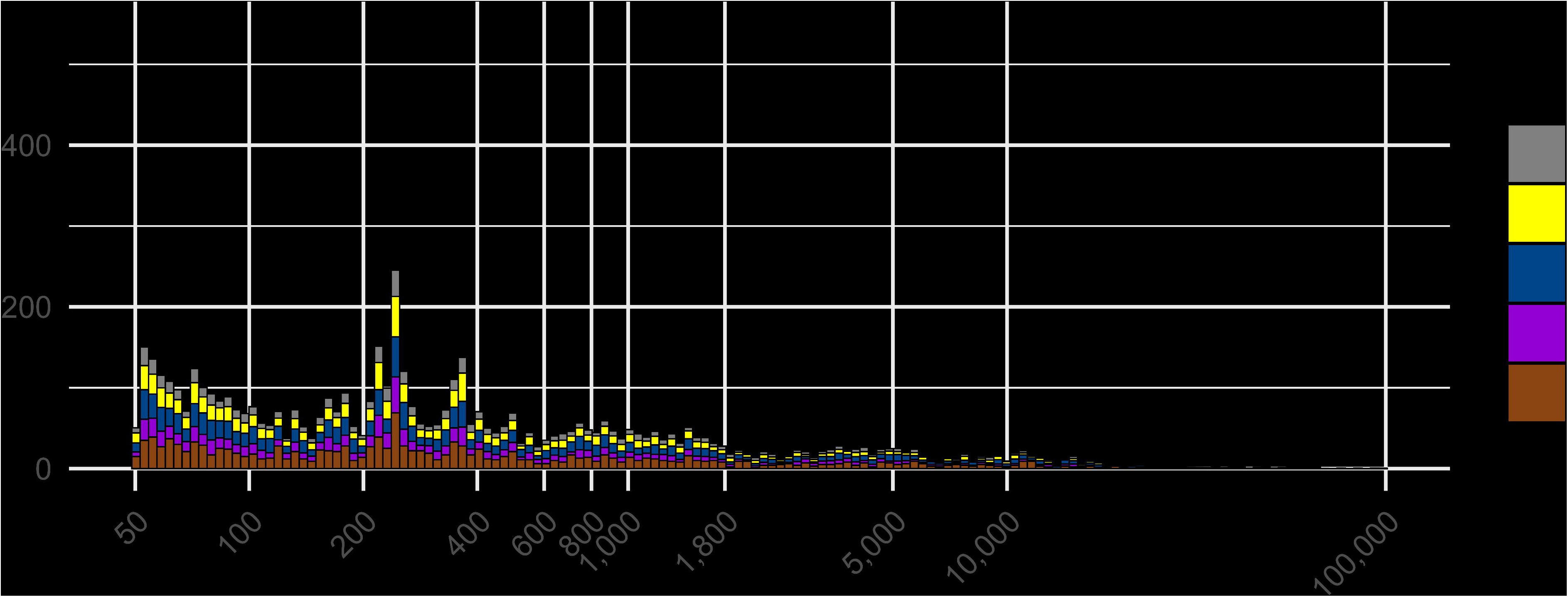

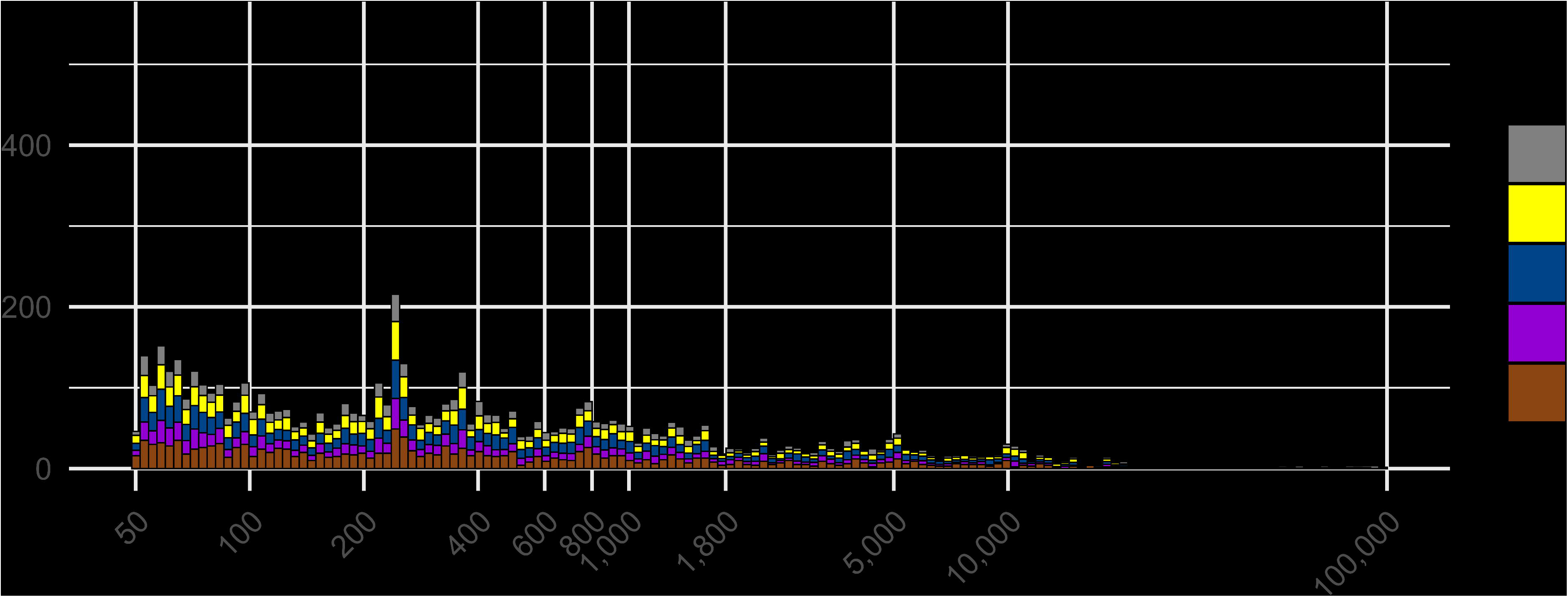

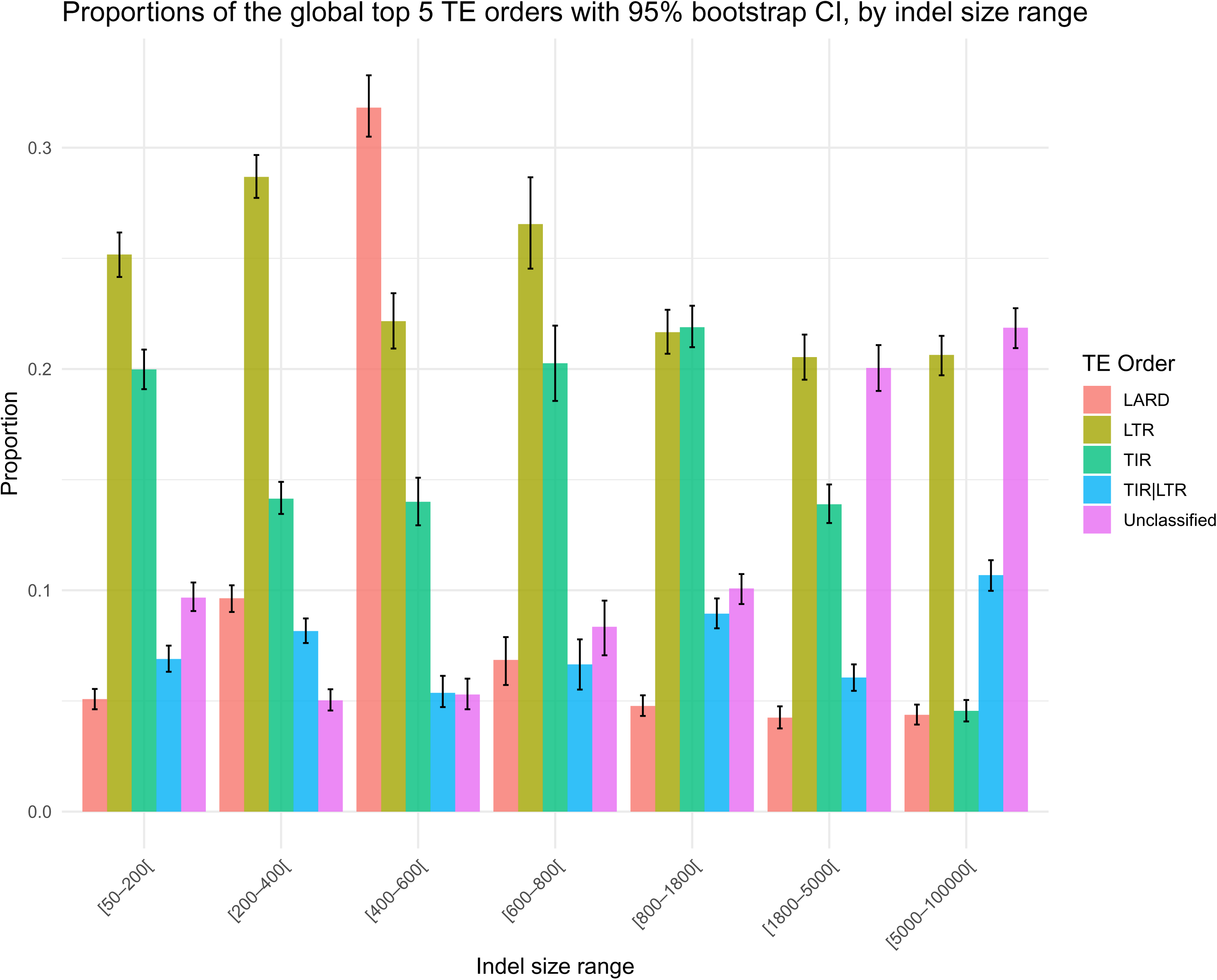

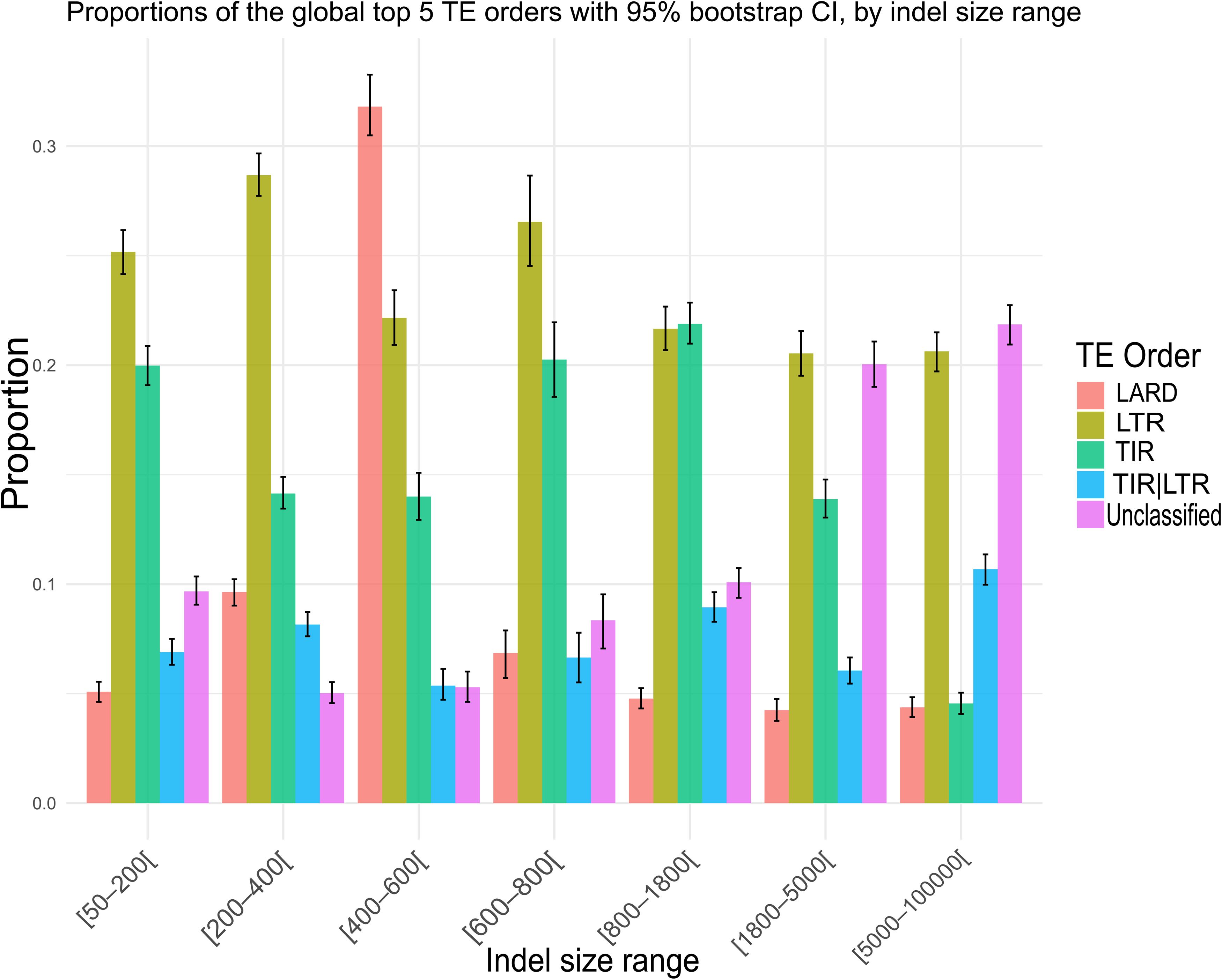

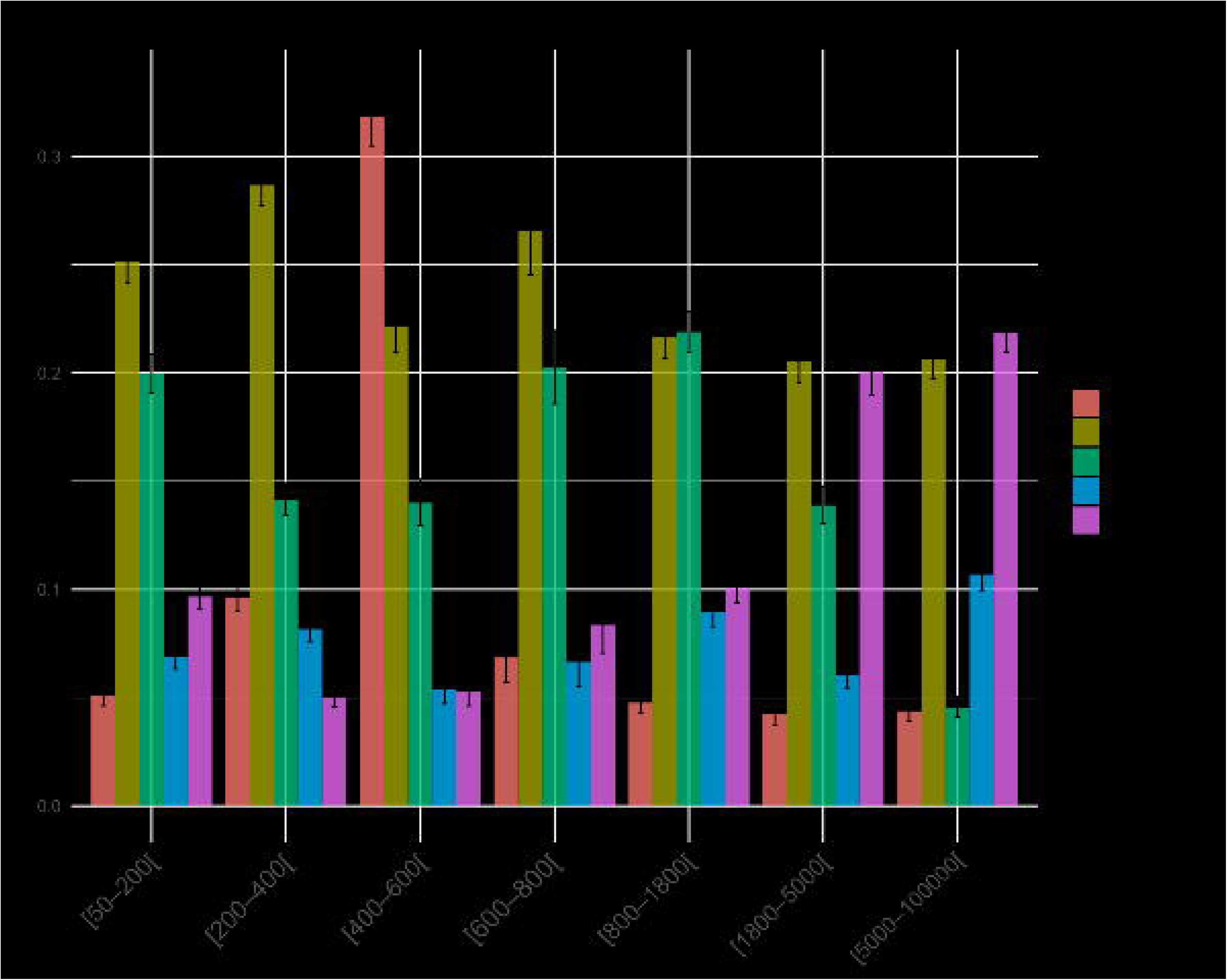

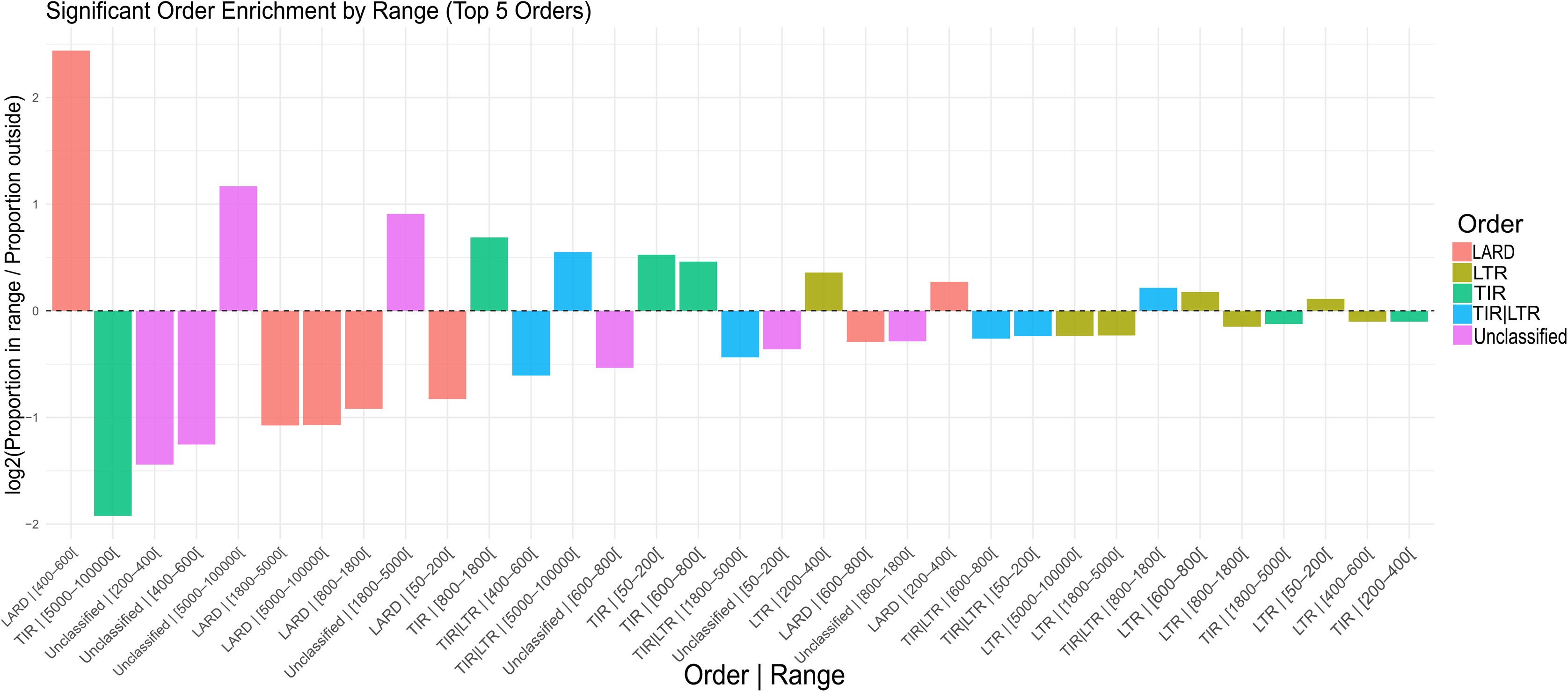

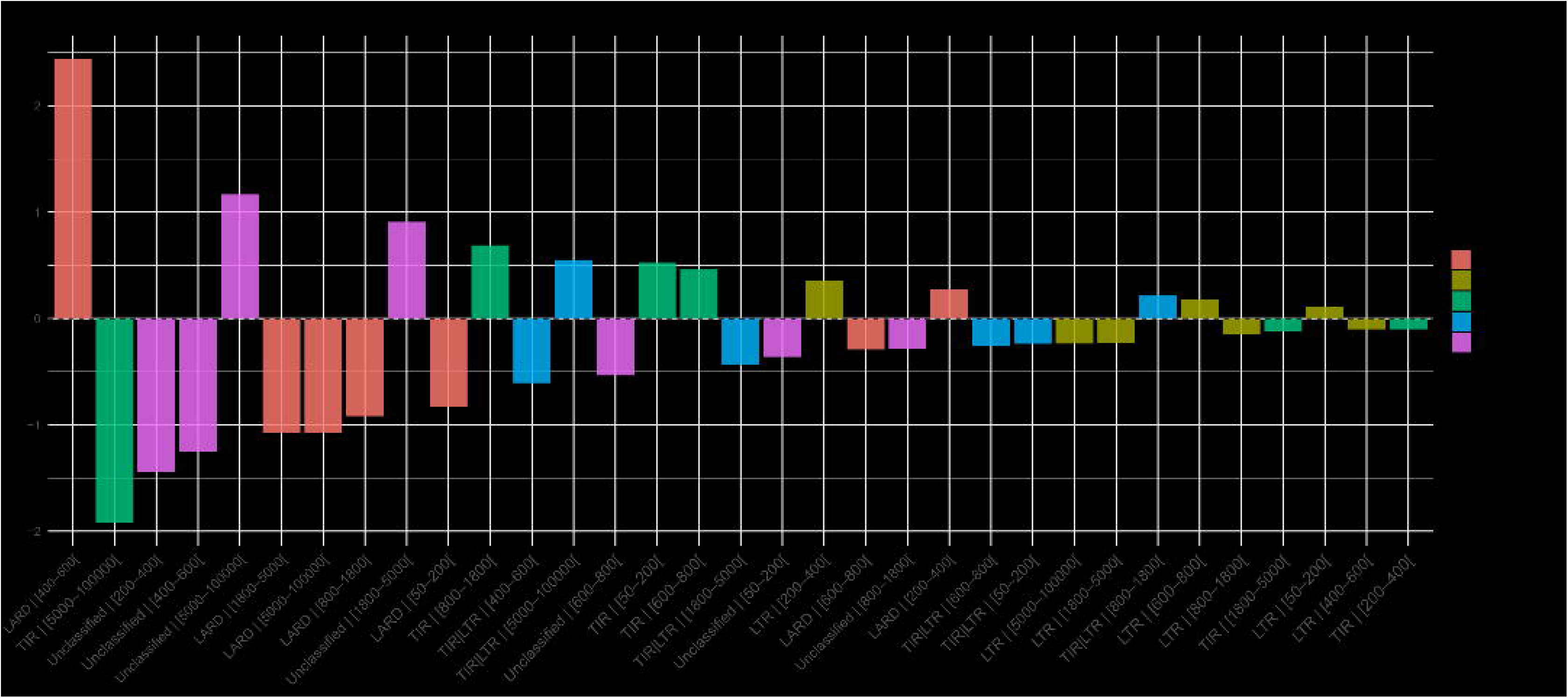

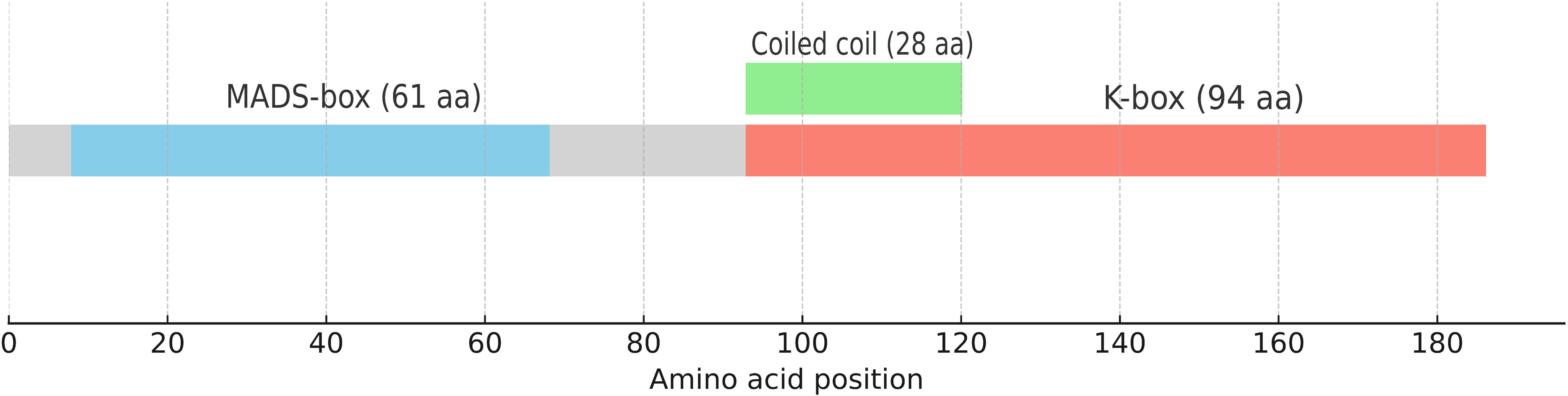

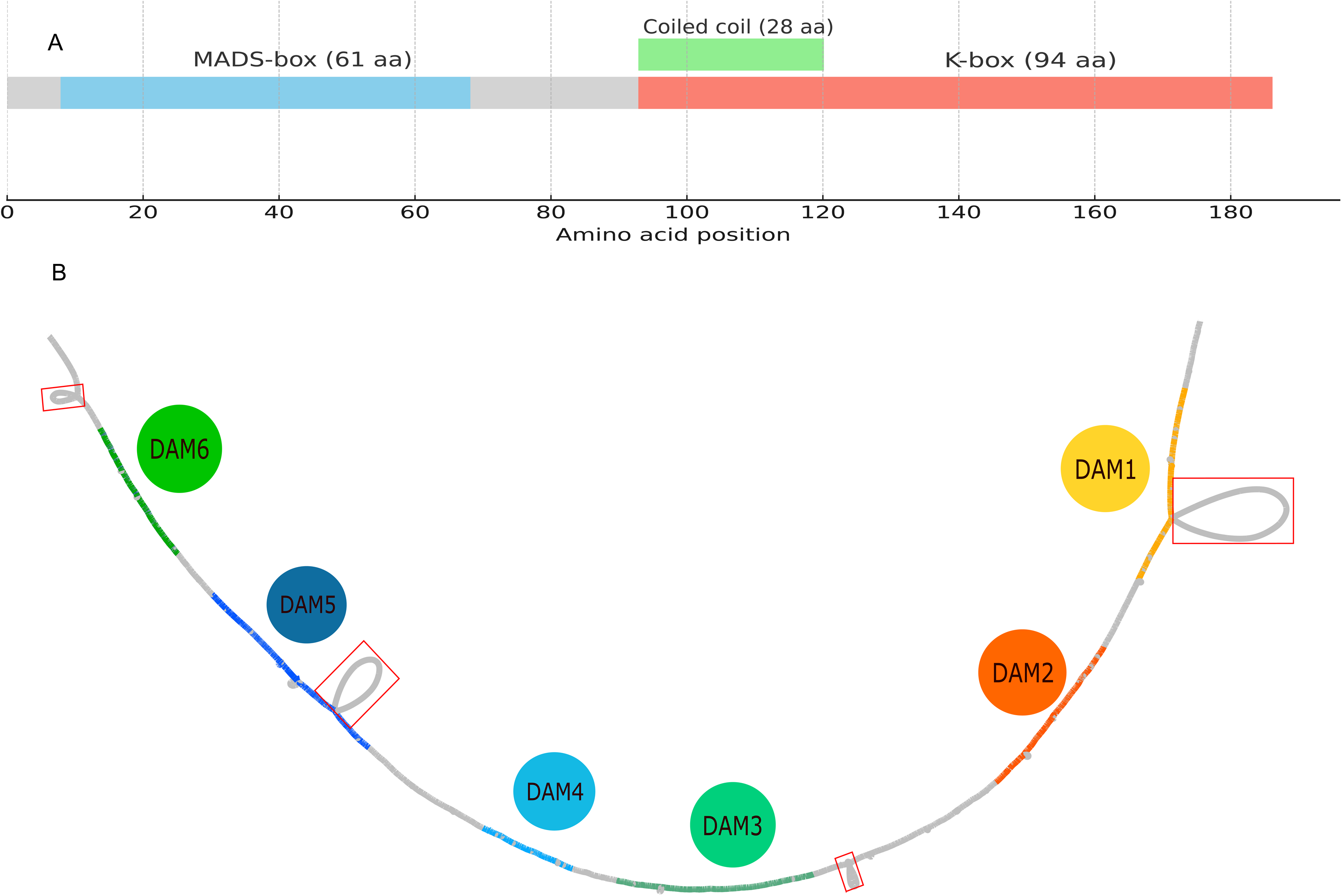

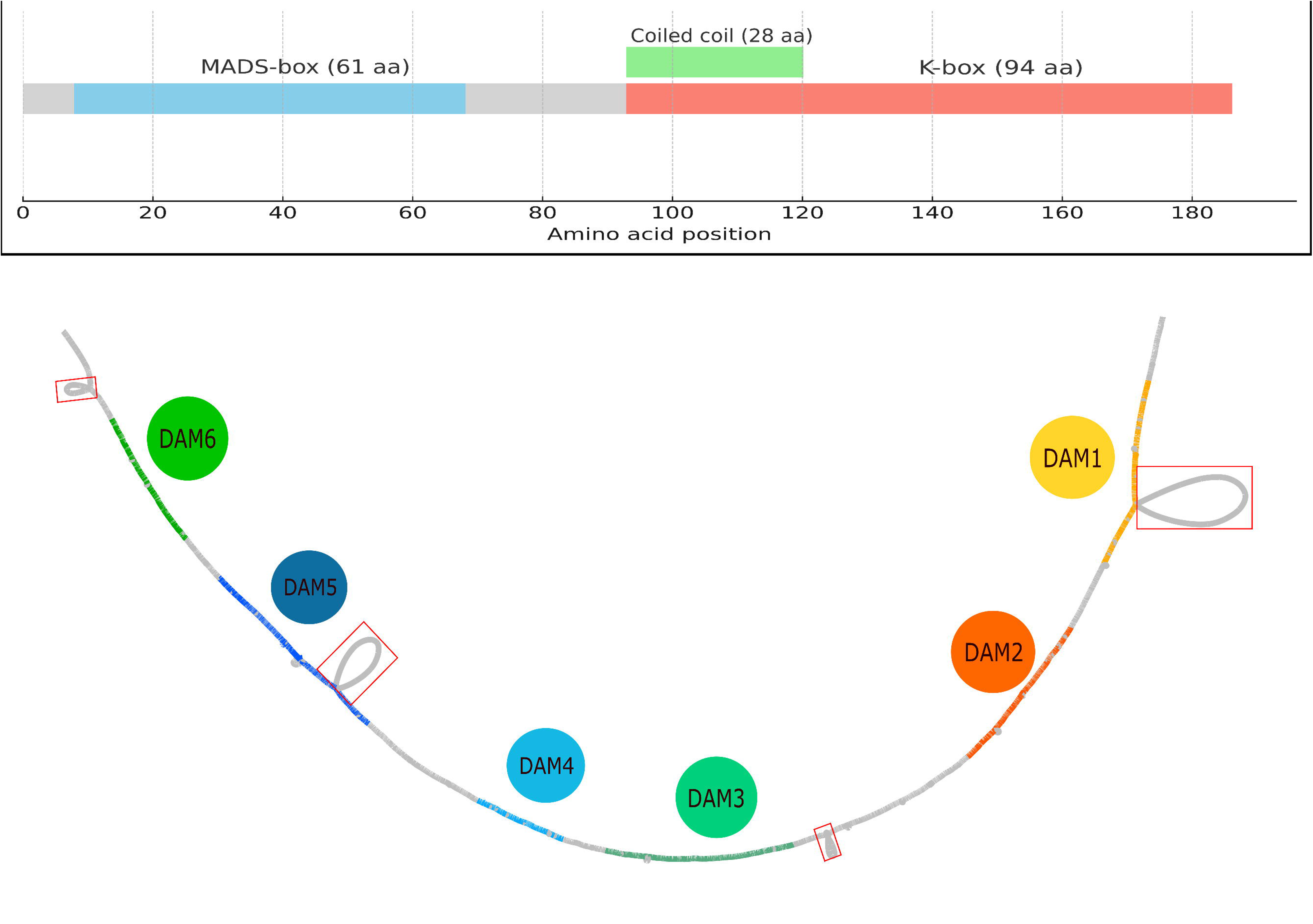

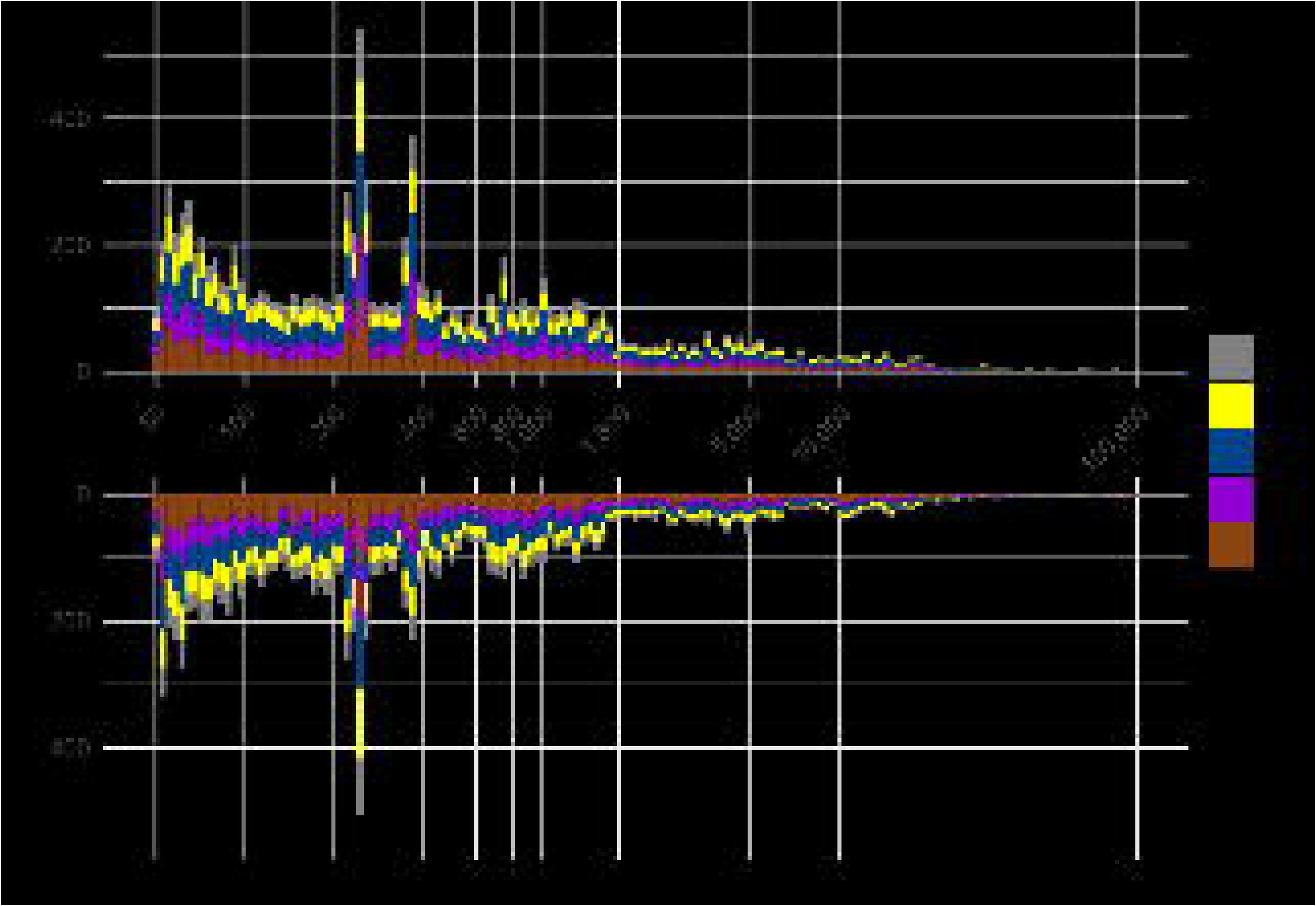

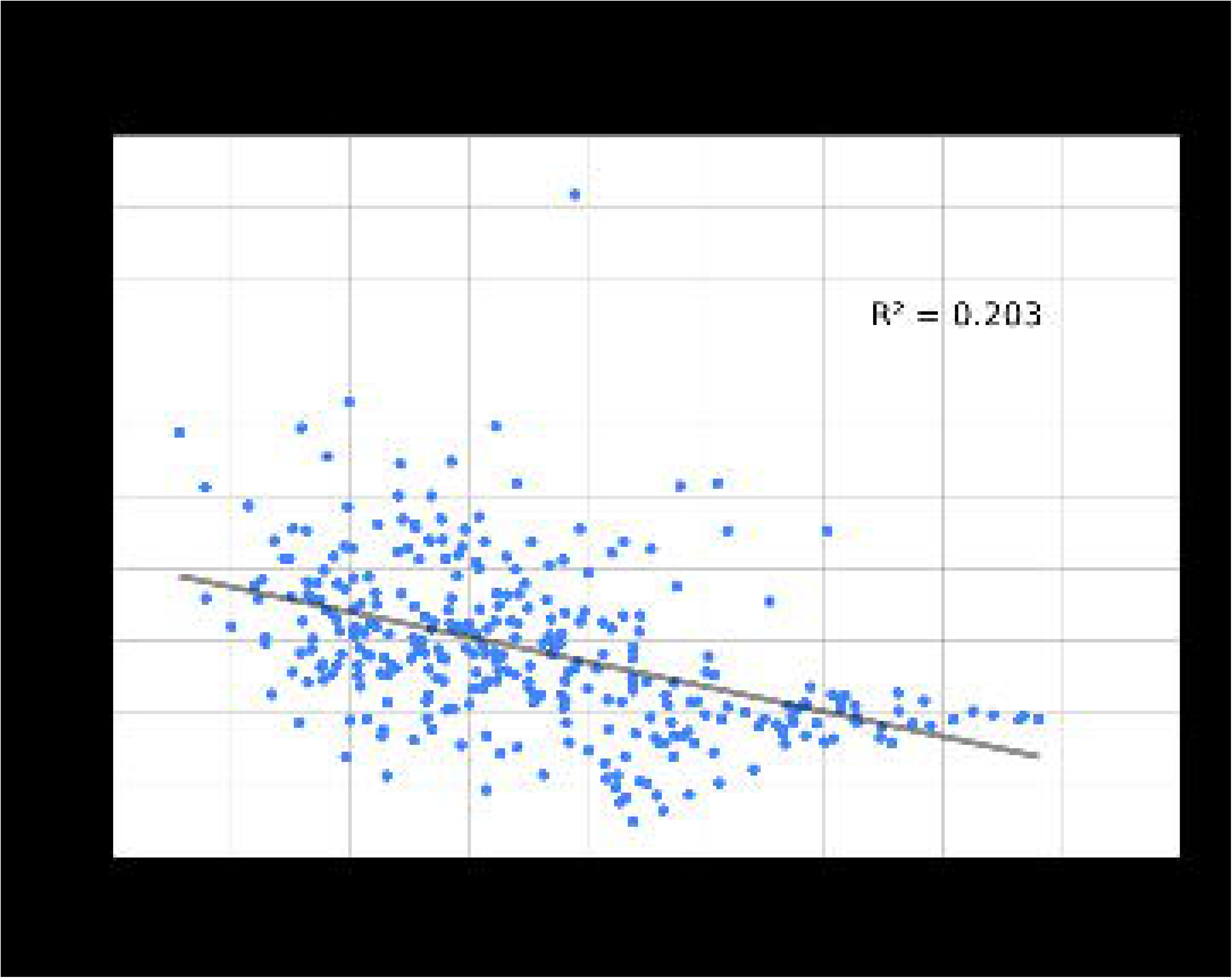

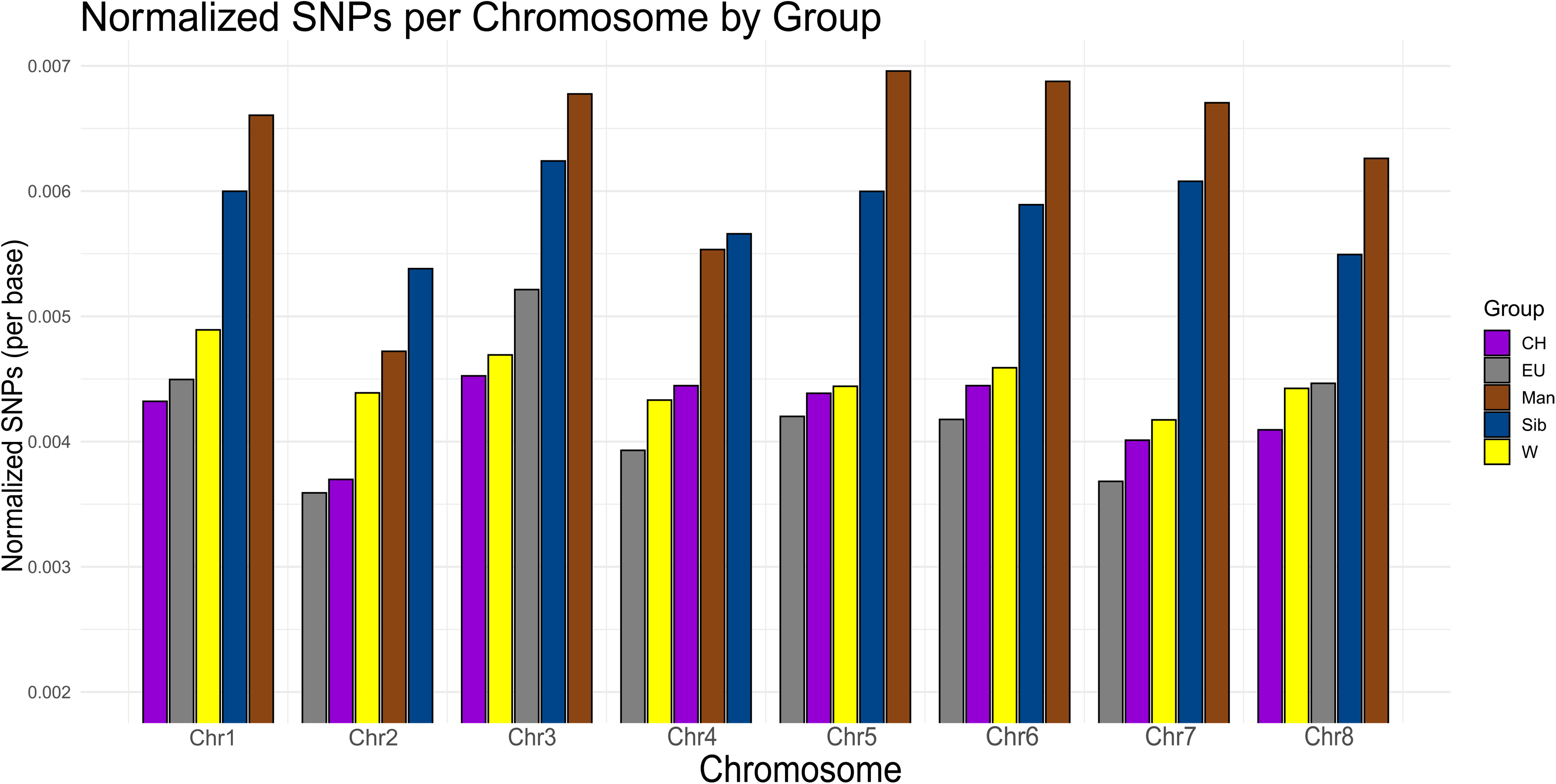

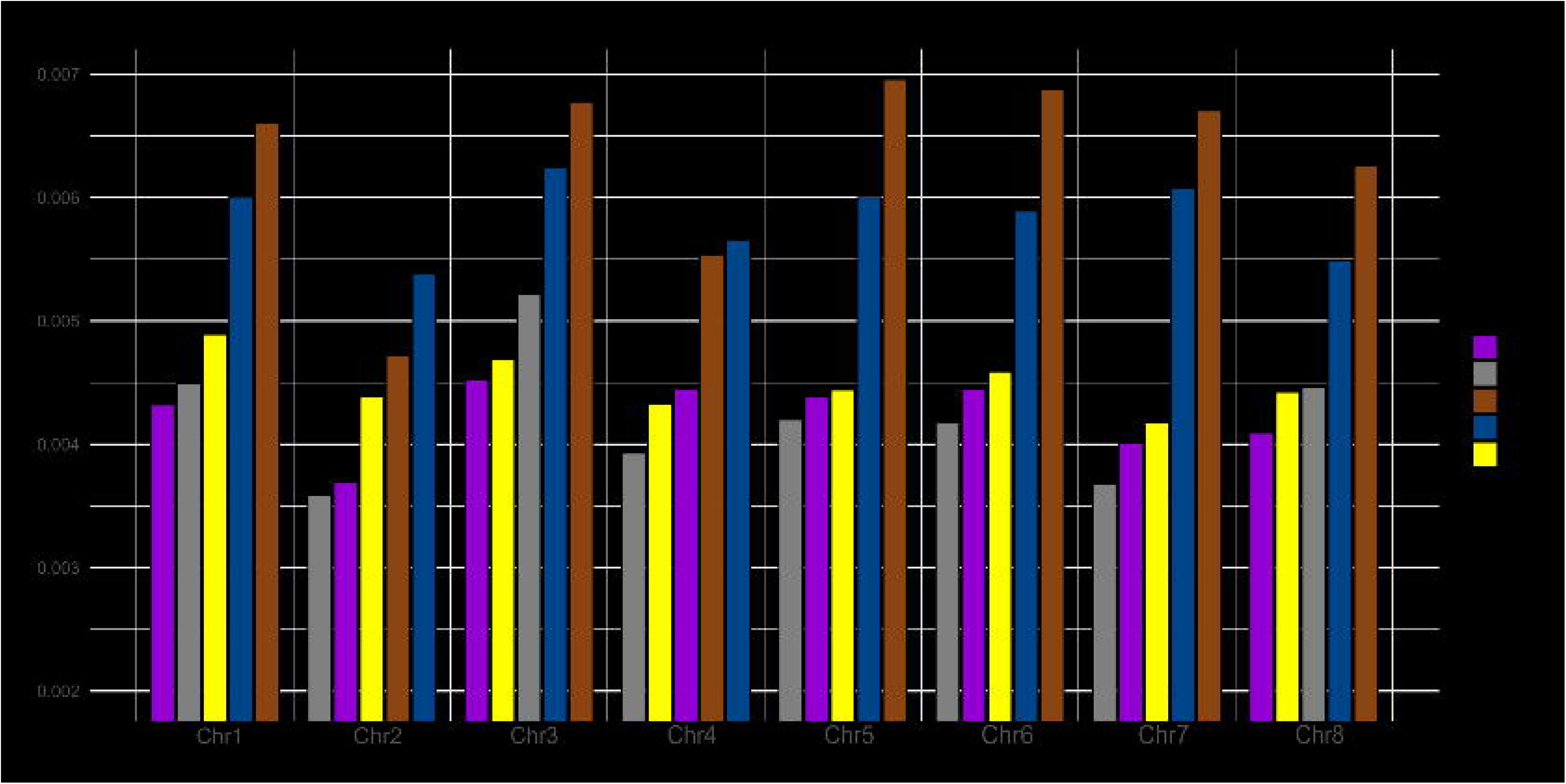

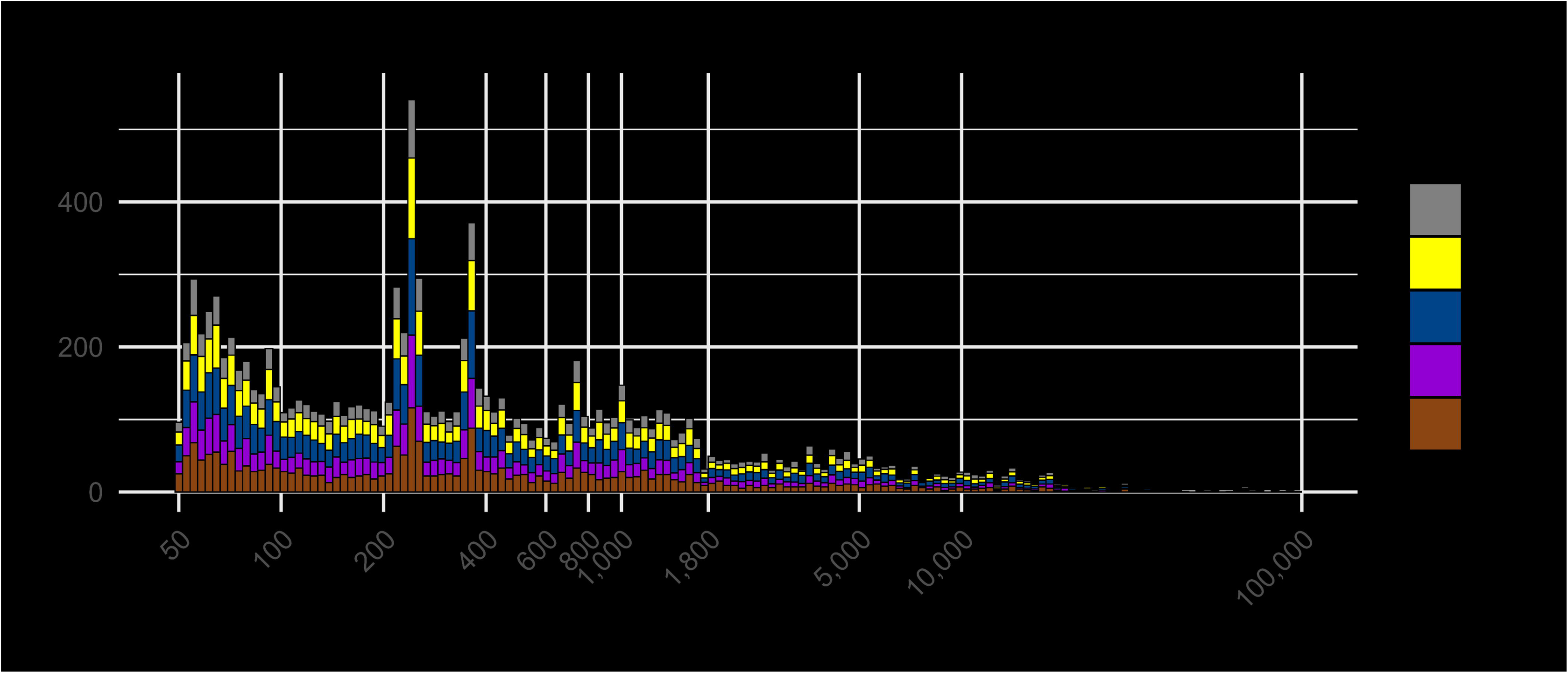

