## Supplementary.file 1 for "Building pangenomes for domesticated and wild tree species: genomic complexity and strategies"

1. INRAE, UMR 1332 BFP, 71 Avenue Edouard Bourlaux, 33882, Villenave d'Ornon, France.
2. INRAE, MIAT, UR 875, Centre Occitanie-Toulouse, 24 Chemin de Borde Rouge, 31320 Auzeville-Tolosane France.
3. INRAE, Univ. Bordeaux, BIOGECO, F-33610 Cestas, France.
4. Université de Rouen Normandie, UFR Sciences et Techniques, 3 Av. Pasteur, 76000 Rouen, France.
5. Université Paris Saclay, INRAE, CNRS, AgroParisTech, GQE - IDEEV, 91190 Gif-sur-Yvette, France / New York University Abu Dhabi, Abu Dhabi, United Arab Emirates
6. Univ. Bordeaux, Centre de Bioinformatique de Bordeaux (CBiB), Bordeaux, 33076, France.
7. URGI - Université Paris-Saclay, INRAE, BioinfOmics, URGI, 78026, Versailles, France

#### Table of contents

**Section 1 : De novo genome assemblies** 2
**Section 2 : Assemblies corrections** 4
**Section 3: Genetic distances relative to reference assembly Rojo_HCUR** 9
**Section 4: Phylogenetic tree of selected assemblies** 10
**Section 5 : Assembly Set and Summary of Graph Metrics** 11
**Section 6: Complements on P. armeniaca Transposon library generation** 13
**Section 7 : Supplements on variants statistics** 14
**Section 8 : Supplements on transposons analyses** 15
**Section 9 : Filtering of reads after mapping** 17
**Section 10 : MAPQ scores distribution following mapping** 20
**Section 11 : Computational cost of mapping** 21
**Section 12 : Data relative to DAM Genomic Regions** 22

### Section 1 : De novo genome assemblies

High-fidelity (HiFi) long-read sequencing data generated using PacBio technology were employed for the *de novo* assembly of six *Prunus armeniaca* (apricot) genomes ( RougeR, KR091, KZ150, CH240, CH250, RRxCH240). For four of these genomes (RougeR, KR091, RRxCH240A and RRxCH240B), assemblies were generated using the Asm4pg v1.1.0 pipeline (https://forge.inrae.fr/asm4pg/GenomAsm4pg), which incorporates integrated steps for quality control, assembly, and polishing optimized for plant genomes. Genome assembly was performed using **hifiasm** (v0.16) (Cheng *et al.*, 2021), followed by scaffolding with **RagTag** (v2.0.1) (Alonge *et al.*, 2022), using the reference genome 'Marouch#14' as a guide. Scaffolds were further anchored and ordered using **ALLMAPS**, based on high-density genetic maps from (Groppi *et al.*, 2021). This approach enabled the placement of over 90% of assembled sequences onto chromosomes.

For the remaining two genomes (KZ150 and CH250), additional data types were incorporated to enhance assembly quality. Long-range scaffolding was performed using optical maps (790X and 3200X, respectively), while **Illumina** short reads were used for polishing and error correction (Groppi *et al.*, 2021). This strategy enabled the anchoring of all sequences onto the sixteen apricot chromosomes of each diploid genome.

In summary, for the KZ150 and CH250 genome assemblies, high-fidelity long-read sequencing data (HiFi reads) were assembled using *hifiasm* v0.15.2 (https://github.com/chhylp123/hifiasm) with default parameters to generate two haplotype-resolved assemblies (hap1 and hap2). The resulting GFA-format graphs were converted into FASTA sequences using *gfatools* v0.4. At this stage, the assemblies consisted of 745 contigs for hap1 and 333 for hap2, with total assembly sizes of 254.2 Mb and 243.8 Mb, respectively. The N50 values were 11.6 Mb (hap1) and 10.3 Mb (hap2). Detailed scaffold statistics were computed directly from the *hifiasm* output. Hybrid scaffolding, based on the optical maps, was, in a second stage, performed at CNRGV (Centre National de Ressources Génomiques Végétales, INRAE Centre de Occitanie-Toulouse), integrating genetic maps with the initial assemblies. Paired-end Illumina reads were quality-filtered using *fastp* and mapped to the scaffolded assemblies using *BWA-MEM*. Average insert size (269 bp) and read length (151 bp) were estimated from *Qualimap* reports. Gap closing was conducted using *GapCloser* with a configuration file based on empirical insert size and read parameters. This process significantly reduced the number of ambiguous 'N' bases: from ~7.9 million to ~37,000 in HS1, and from ~1.6 million to ~32,000 in HS2. Minor reductions in total assembly size were observed post-gap closure. In silico PCR was then performed using the UCSC *isPCR* tool against both haplotype assemblies (HS1 and HS2) using a marker set comprising SNPs and SSRs. Results were filtered and used to anchor genetic markers to the scaffolds. In a third stage of the assemblies, the ALLMAPS pipeline (https://github.com/tanghaibao/jcvi/wiki/ALLMAPS) was used to order and orient scaffolds into chromosome-scale pseudomolecules based on the integrated genetic maps. PCR results were parsed to generate BED input files. The final assemblies consisted of eight ordered scaffolds per haplotype (HS1 and HS2), representing the eight chromosomes, with total sizes of 218.5 Mb and 214.1 Mb, respectively. Unscaffolded sequences totaling ~2.8 Mb were retained separately. Alignments against a reference genome (Marouch #14) using D-GENIES revealed an inversion on chromosome 7 in both haplotypes. The corresponding chromosome sequences were extracted, reverse-complemented using *EMBOSS revseq*, and reintegrated into the respective assemblies. Finally, to correct for potential phasing inconsistencies, chromosome 1 was swapped between the HS1 and HS2 assemblies. Individual chromosome sequences were extracted and merged using custom *awk*-based scripts and cat commands to regenerate the final genome assemblies for each haplotype.

For all genomes, assembly completeness was evaluated using **BUSCO** (v5.3.1) ((Manni *et al.*, 2021, Preprint)) with the *eudicots_odb10* lineage dataset, with all assemblies achieving completeness scores exceeding 98%. The final assemblies exhibited high contiguity, with N50 values ranging from 21 to 29 Mb and L50 values between 3 and 4. Whole-genome alignments were visualized using **D-GENIES** ((Cabanettes and Klopp, 2018)) to assess structural accuracy and, where necessary, correct chromosome orientations.

### Section 2 : Assemblies corrections

Before being used as an input to pangenome graph construction using minigraph-cactus, several assemblies were manually corrected.  These corrections belongs to two main categories:

1. Inconsistent chromosome number labeling
2. Inconsistent chromosome sequence orientation

To assess the genomic relationships across assemblies, Mash distances were calculated for every chromosome against all selected scaffolds within each assembly. A Mash distance of 0 indicates identical k-mer content between two sequences, while increasing values signify greater divergence.
As depicted in the figure below, which illustrates comparisons for chromosome 1 across a subset of assemblies (represented on the T-axis), this analysis successfully identified erroneous chromosome labeling [*P. mume* tortuosa and BJFU and *P. salicina* Sanyueli_FAAS].

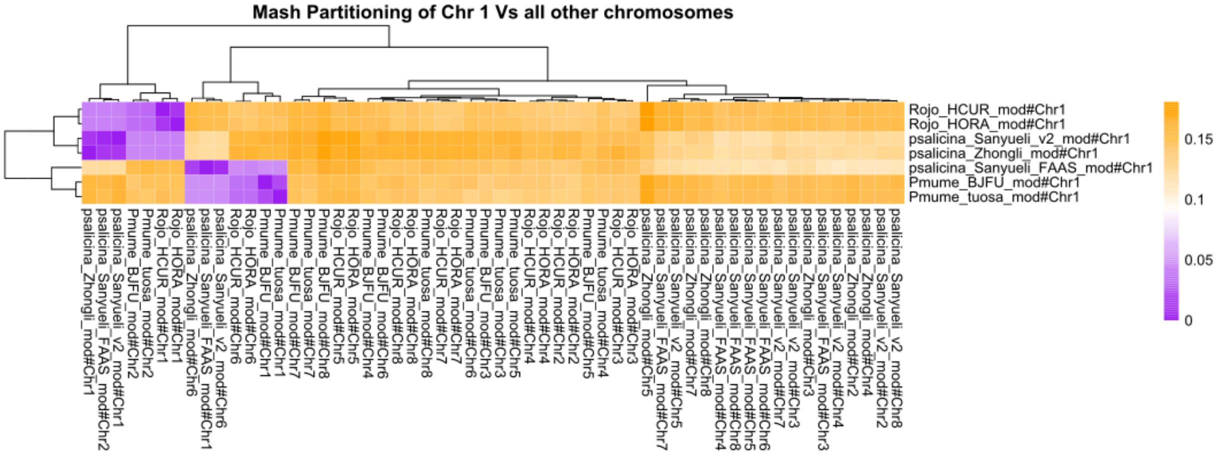


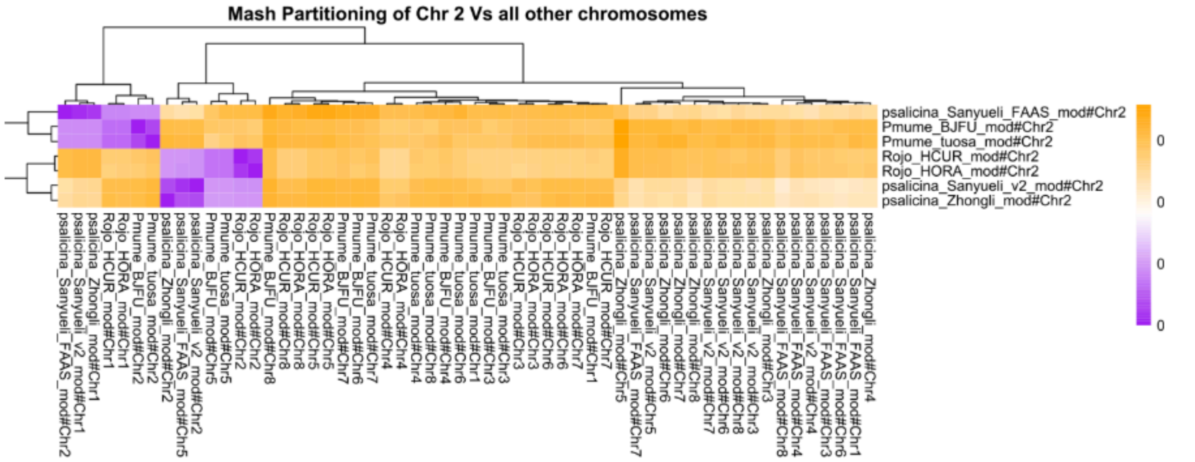

**Mash Partitioning of Chr 3 Vs all other chromosomes**

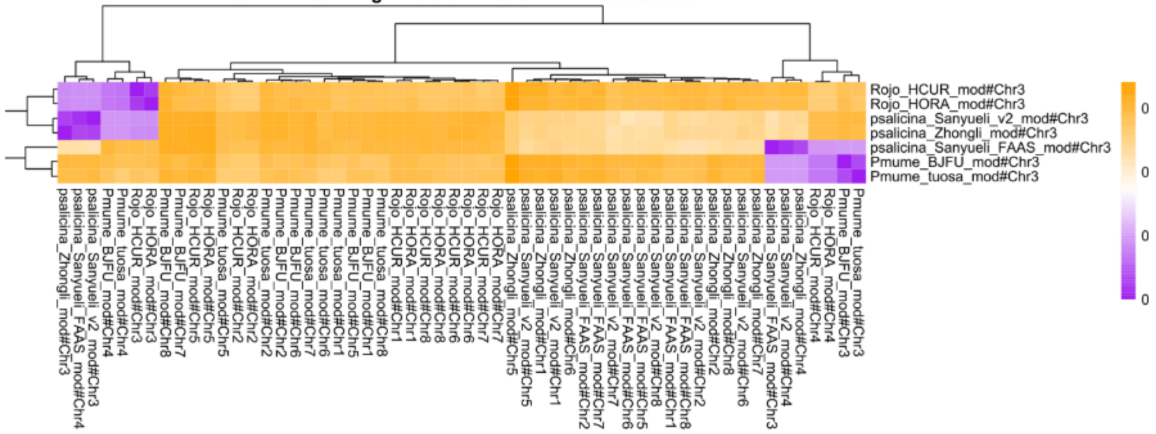


**Mash Partitioning of Chr 4 Vs all other chromosomes**

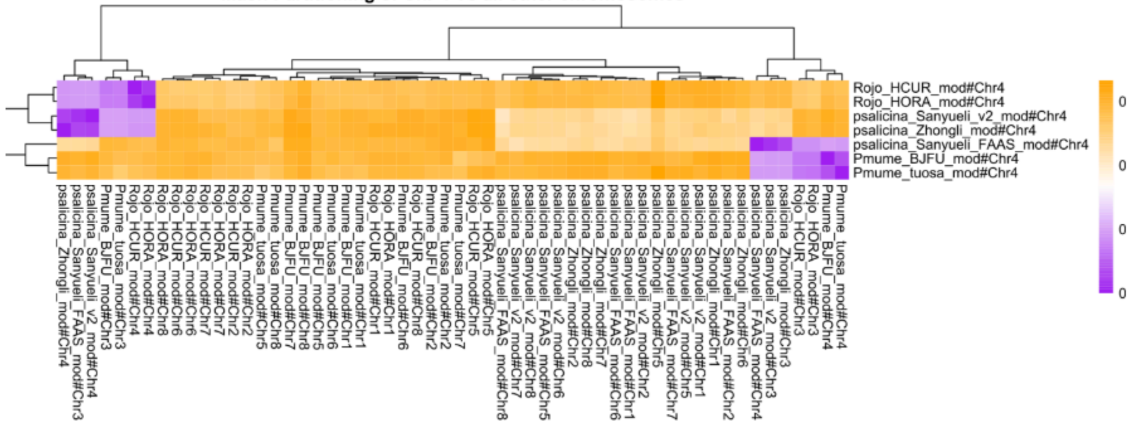


**Mash Partitioning of Chr 5 Vs all other chromosomes**

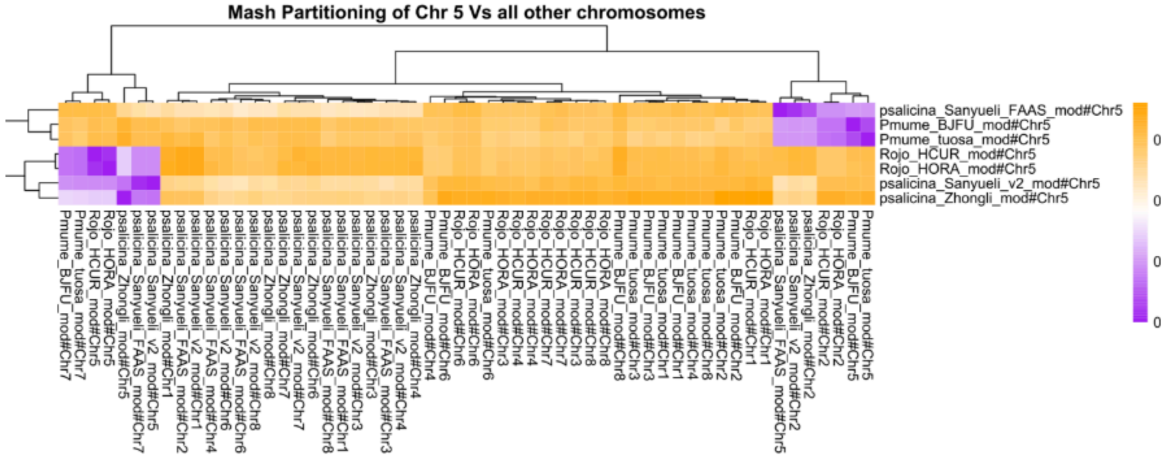

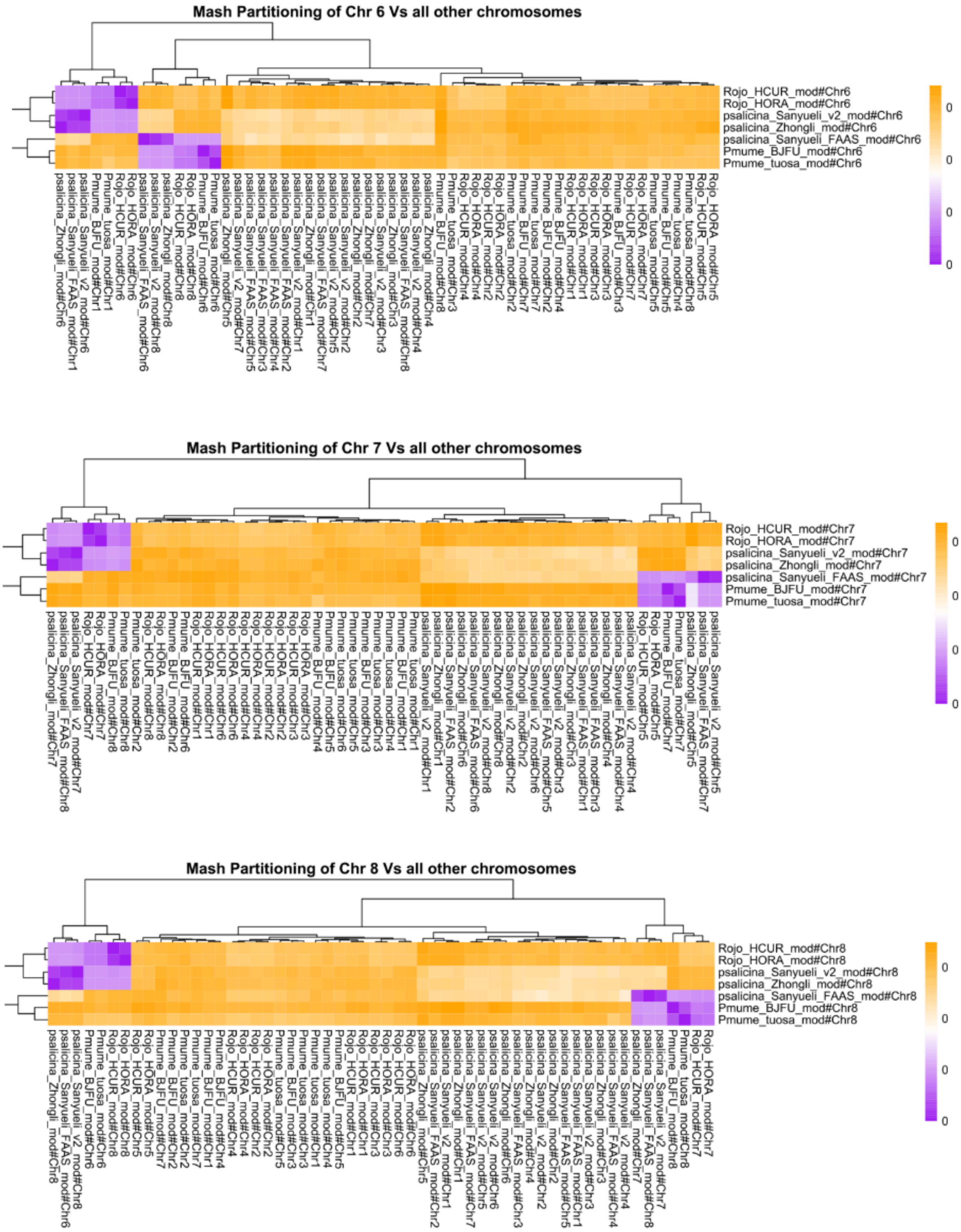


*Figure F1-A to F1-H: Inconsistencies in Chromosome 1 to 8 Labeling Across Prunus Assemblies. Analysis of Mash distances for chromosome 1 to 8 across all retrieved Prunus assemblies revealed notable discrepancies in chromosomal labeling. A Mash distance approaching 0 indicates a high degree of similarity in k-mer content, signifying close genetic proximity.*

Intriguingly, chromosome 1 of our reference assembly (Rojo_HCUR) exhibited an unexpected high proximity (low Mash distance) with chromosome 2 in several assemblies, specifically those of *P. salicina*, *P. mume* BJFU, *P. mume* tortuosa. This finding strongly suggests inconsistencies in chromosome labeling within these assemblies when compared to standard Prunus genomic nomenclature. Equivalent figures can be found for other chromosomes in Suppl Figures F1-A to F1-H.

After applying this analysis to the 8 chromosomes, we relabeled scaffold sequences in every assemblies, as described in the following text file : All_assemblies_uniformed_chromosome_label.txt:

For the GZYX assemblies, manual re-orientation of several chromosomes was necessary to align them with the established orientations depicted in the Genome Database for Rosaceae (GDR, www.rosaceae.org). Given the observed colinearity across *Prunus* genomes, we utilized the eight-chromosome Peach genetic architecture as our reference. The initial state of these misorientations is illustrated in the first figure, with the subsequent figure demonstrating the successful manual correction. While some smaller regions (e.g., within chromosome 2) appear to retain a reversed orientation, the predominant correct orientation of their respective chromosome sequences led us to interpret these as putative genuine inversions. Consequently, these were preserved without modification for the construction of the final graph.

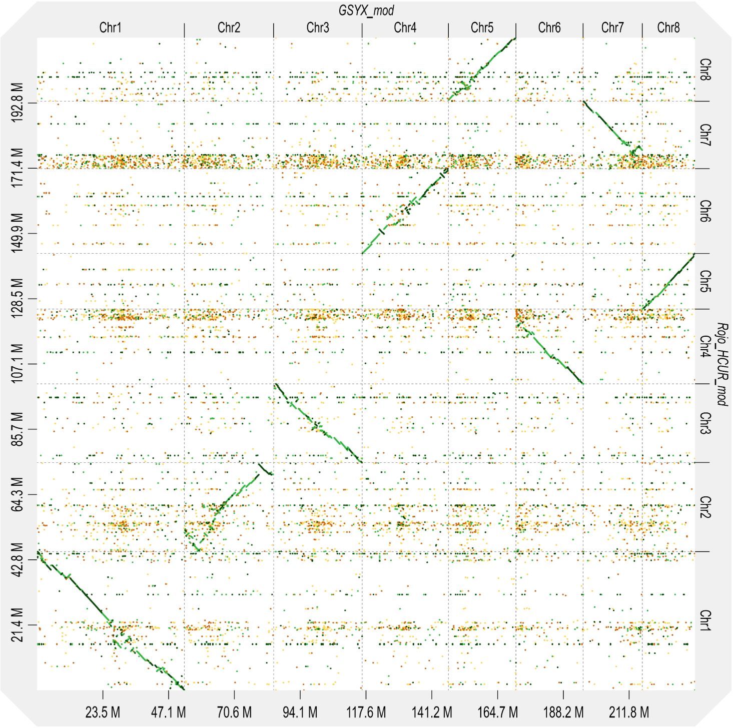

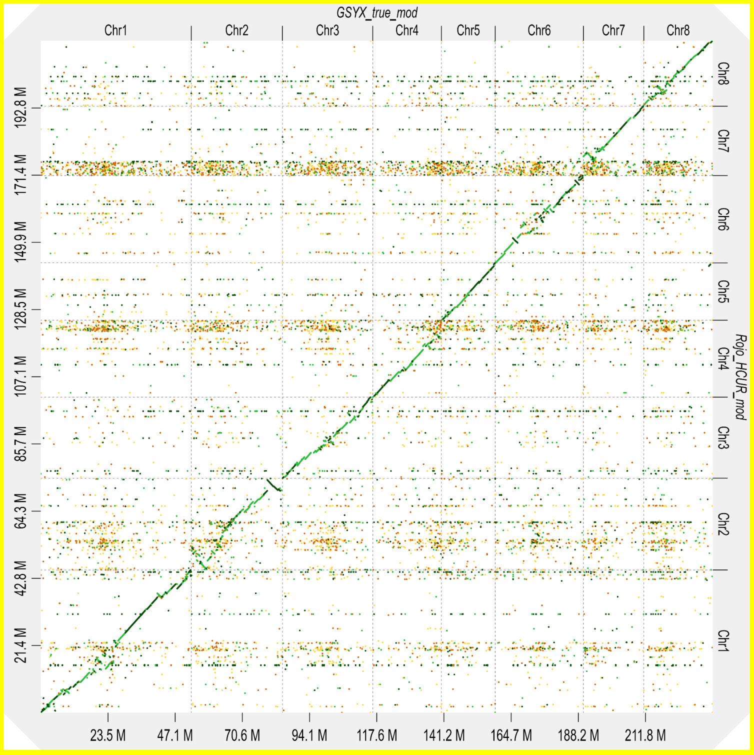


**Figure 1** : **Dotplot comparison of the GSYX haplotype before and after correction.** *Dotplots generated with D-Genies show the alignment of the GSYX assembly against the reference (Rojo_HCUR) before (left) and after (right) correction. The correction process improves structural consistency and continuity, reducing noise and misalignments, particularly in complex or repetitive regions.*

### Section 3: Genetic distances relative to reference assembly Rojo_HCUR

Using pantools and k-mer distances between assemblies, a genetic distance relative to reference assembly Rojo_HCUR has been estimated (Figure 2). The distance matrix can be found in Suppl Table T2.

Based on these results, we established a distance threshold of 0.03. All assemblies falling below this value were retained, as illustrated in the figure below. Despite its proximity to this threshold, the *P. mandshurica* CH_264 assembly was also included to ensure the representation of at least one outgroup within the pangenome graph.

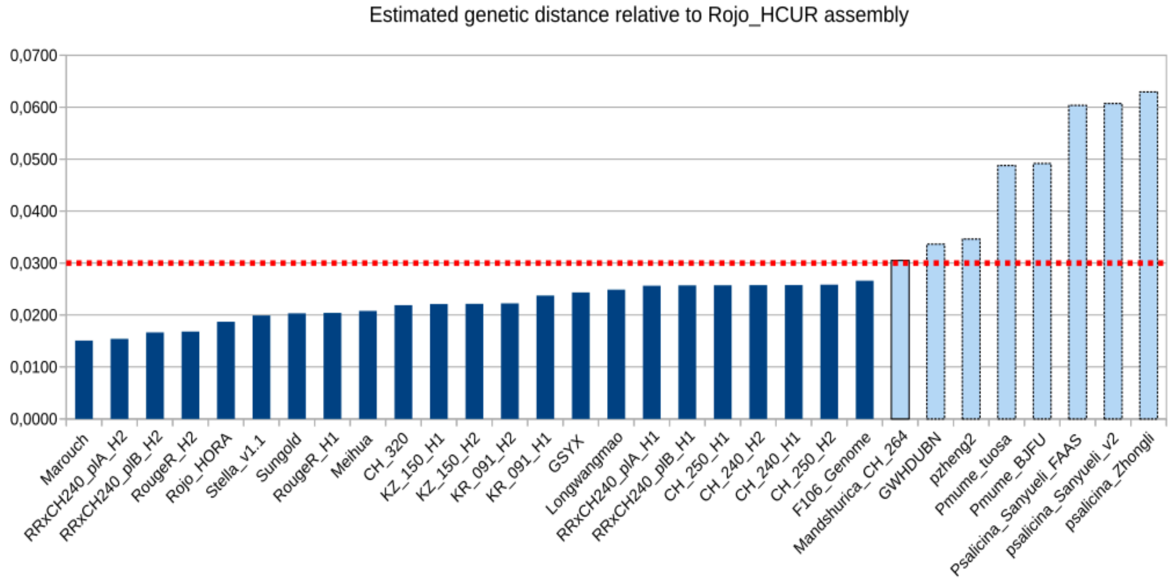


**Figure 2 : Estimated genetic distance relative to Rojo_HCUR assembly**
*Assemblies used for the k-mer-based phylogenetic analysis are displayed along the X-axis, ordered by their genetic distance to the reference Rojo_HCUR. The Y-axis represents the genetic distance, with a threshold of 0.03 indicating the maximum value considered in the graph.*

### Section 4: Phylogenetic tree of selected assemblies

The phylogenetic tree representing the 25 assemblies chosen for graph construction is provided below in both Newick and graphical formats. This tree exhibits a Robinson-Foulds (RF) distance of 2 when compared to the tree encompassing all assemblies, indicating a high degree of topological concordance with only two differing bipartitions.

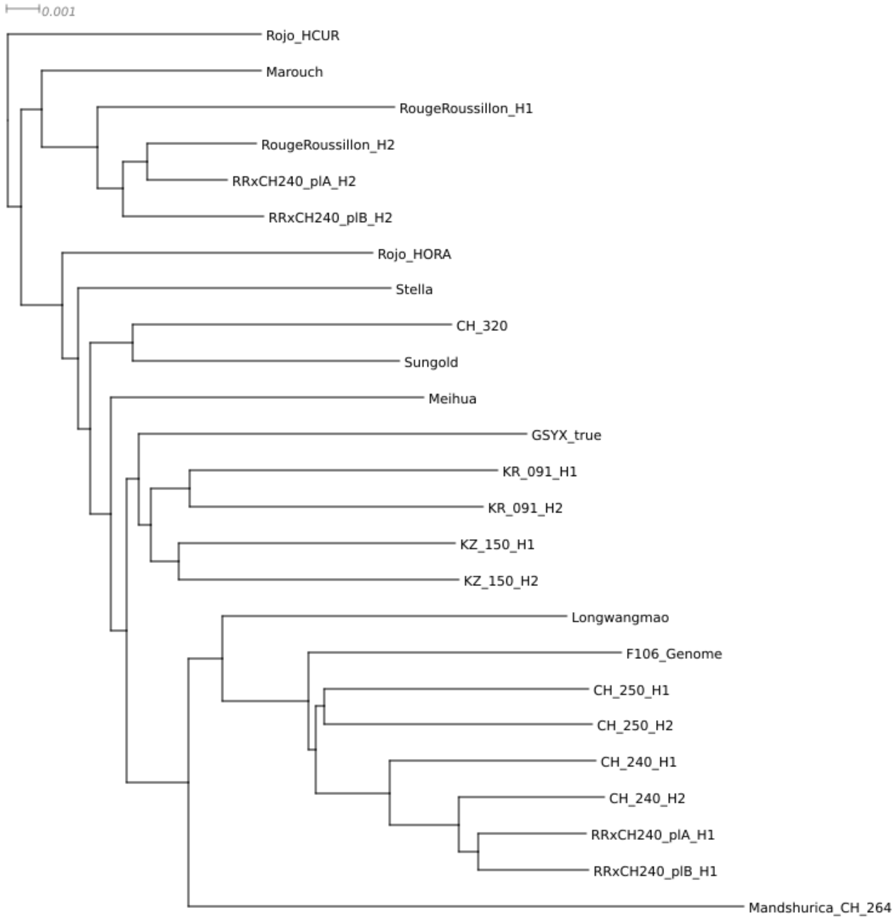


Corresponding newick tree, given as input to Minigraph-cactus:

```
(Rojo_HCUR:0.00788576,((Marouch_v3.1:0.00681225,(RougeRoussillon_H1:0.00924739,(RRxCH240_1_plB
_H2:0.00434348,(RRxCH240_1_plA_H2:0.00247215,RougeRoussillon_H2:0.00337104):0.00074349):0.0008
2004):0.00170963):0.00065576,(Rojo_HORA:0.00966848,(Stella_v1.1:0.00971425,((Sungold:0.0082678
6,CH_320_5:0.00991524):0.00134332,(Meihua:0.00973899,((GSYX_true:0.0120919,((KZ_150_8_H1:0.008
60665,KZ_150_8_H2:0.0087122):0.00085192,(KR_091_H1:0.00959655,KR_091_H2:0.0091358):0.00118202)
:0.00039442):0.00036758,(Mandshurica_CH_264_4:0.01728973,(Longwangmao:0.01074548,(F106_Genome:
0.0097333,((CH_250_H1:0.00823693,CH_250_H2:0.00832687):0.00026228,(CH_240_1_H1:0.00639755,(CH_
240_1_H2:0.00452189,(RRxCH240_1_plA_H1:0.00336376,RRxCH240_1_plB_H1:0.00344785):0.00057376):0.
00216333):0.00230586):0.0002406):0.00267844):0.00105252):0.00191063):0.00048916):0.0006502):0.
00036991):0.00048354):0.00128998):0.00041504);
```

### Section 5 : Assembly Set and Summary of Graph Metrics

A series of supplementary tables accompanies this study to provide detailed insights into the structure, composition, and diversity of the apricot pangenome:

| Group | Assembly label | Haplo[1] | Species | Population[2] | Technologies[3] |
|---|---|---|---|---|---|
| EU | **Marouch** | | *P. armeniaca* | C1 | PB, SR |
| | **Rojo_HORA** | | | C1 | PB, HiC, 10x, SR |
| | **Rojo_HCUR** | | | C1 | PB, HiC, 10x, SR |
| | **Stella** | | | C1 | PB, SR |
| | **RougeR** | *H1* | | C1 | PB-hi |
| | **RougeR** | *H2* | | C1 | PB-hi |
| CH | **Sungold** | | *P. armeniaca* | NA | PB, HiC,BN |
| | **GSYX** | | | NA | PB-hi |
| | **Meihua** | | | NA | PB,HiC, SR |
| W | **KZ150** | *H1* | *P. armeniaca* | W2 | PB-hi, SR, BN |
| | **KZ150** | *H2* | | W2 | PB-hi, SR, BN |
| | **KR091** | *H1* | | W1 | PB-hi |
| | **KR091** | *H2* | | W1 | PB-hi |
| | **CH320** | | | NA | ONT |
| Sib | **Longwangmao** | | *P. armeniaca* | NA | PB, HiC,BN |
| | **CH240** | *H1* | *P. sibirica* | W4 | PB-hi |
| | **CH240** | *H2* | | W4 | PB-hi |
| | **CH250** | *H1* | | W4 | PB-hi, SR, BN |
| | **CH250** | *H2* | | W4 | PB-hi, SR, BN |
| | **F106** | | | NA | PB, HiC, BN |
| Hyb | **RRxCH240A_H1** | *H1* | *P.sibirica x P. armeniaca* | NA | PB-hi |
| | **RRxCH240A_H2** | *H2* | | | PB-hi |
| | **RRxCH240B_H1** | *H1* | | | PB-hi |
| | **RRxCH240B_H2** | *H2* | | | PB-hi |
| Mand | **Pmand CH264** | | *P. mandshurica* | *P. mandshurica* | PB, ONT, SR |
| Mum | **Pmume Tortuosa** | | *P. mume* | *P. mume* | ONT, HiC, SR |
| | **Pmume BJFU** | | | NA | SR |
| Other | **GWHDUBN** | | *P. armeniaca var. armeniaca syn. Prunus hongpingensis* | NA | PB-hi |
| | **Zheng 2** | | *Prunus zhengheensis* | | PB-hi |
| Sal | **Sanyueli_v2** | | *P.salicina* | NA | PB, SR |
| | **Zhongli** | | | | ONT |
| | **Sanyueli_FAAS** | | | | PB, SR |

1. Phased asssemblies were split as H1/H2 as they can be represented as different paths in a pangenome graph
2. Genetic clustering based on Groppi et al
3. PB=PacBio ; PB-hi=PacBio Hifi ; HiC = Hi-C-based scafolding ; ONT= Oxford Nanopore ; BN= BioNano ; 10x=10x scCNV ; SR=Short reads

Suppl. Table_T1 : List of assemblies recovered for this study and their classification. This table lists the 32 genome assemblies included in the analysis, along with their classification into phylogeographic groups. Only 25 assemblies [Eu to Mand] are used in the pangenome. It serves as a foundational reference for understanding the genomic diversity incorporated into the pangenome.

| | Assemblies | | General statistics | | | | | Pansel statistics | | | |
|---|---|---|---|---|---|---|---|---|---|---|---|
| | Average bp | Total bp | Total bp | # Nodes | # Edges | Mean node length (bp) | Compress-ion factor | #Conserved (p<0.05) | #Divergent (p<0.05) | Normalised (x10e7) | Normalised (x10e6) |
| **Chr1** | **45,544,716** | **1,456,491,526** | **1,152,426,215** | **6,669,924** | **9,294,083** | **27.14** | 6.37 | **296** | **1524** | 2.57 | 1.32 |
| **Chr2** | 31,106,144 | 997,605,847 | 719,721,432 | 4,406,510 | 6,124,062 | **40.71** | **4.01** | 135 | 749 | **1.88** | **1.04** |
| **Chr3** | 27,353,395 | 875,784,375 | 666,071,915 | 4,266,339 | 5,940,672 | 30.86 | 5.06 | 139 | 943 | 2.09 | 1.42 |
| **Chr4** | 28,079,035 | 898,953,419 | 690,000,886 | 4,033,618 | 5,595,057 | 38.45 | 4.45 | 160 | 761 | 2.32 | 1.10 |
| **Chr5** | **19,979,280** | **641,443,444** | **473,654,461** | **2,697,166** | **3,765,995** | 27.52 | **6.38** | 237 | 846 | **5.00** | **1.79** |
| **Chr6** | 29,294,471 | 937,435,313 | 722,911,288 | 4,204,035 | 5,861,567 | 32.8 | 5.24 | 215 | 1187 | 2.97 | 1.64 |
| **Chr7** | 23,291,025 | 746,953,785 | 582,577,180 | 3,428,200 | 4,768,682 | 35.97 | 4.72 | 175 | 797 | 3.00 | 1.37 |
| **Chr8** | 22,674,453 | 726,961,516 | 553,445,739 | 3,371,858 | 4,687,883 | 37.03 | 4.43 | **124** | **579** | 2.24 | 1.05 |
| **ALL** | | 7,281,629,225 | 5,560,809,116 | 33,077,650 | 46,038,001 | | | 1282 | 7363 | 2.31 | 1.32 |

Suppl. Table_T2: General statistics for the eight chromosome-level pangenome graphs.
In our pangenome graph analysis, several key metrics were employed to characterize genomic variation. The compression factor quantifies the efficiency of the graph representation, calculated as the ratio of total base pairs in the input assemblies to the total base pairs within the constructed graph. The columns labeled “Conserved regions” and “Divergent regions” report the count of genomic segments identified as significantly more conserved or divergent, respectively, relative to the entire pangenome graph and as established by the PanSel tool definition. The two rightmost columns provide a standardized measure of these regional counts by normalizing them against the corresponding chromosome length in the chosen reference assembly. The extreme values (highest and lowest) are highlighted in bold.

These statistics are complemented by several tables :

The details of Panacus values (main manuscript, figure 2) are in Suppl. Table_T3 : Ordered histgrowth in bp all chromosome.xlsx . This Excel file reports the distribution of core (red) and non-core (green) genome content along each chromosome, under two quorum conditions (0 and 1) corresponding to the Panacus execution parameters used in Figure 2 of the main manuscript. The same columns are present in all tabs, and further details on the quorum and coverage settings can be found in the Panacus documentation: https://github.com/codialab/panacus.

Suppl.Table_T4 : Proportion of the different genome types along the 8 chromosome.xlsx
This table provides the different proportions of core, private and accessory genome for every assembly in the graph, and for every chromosome.

Suppl.Table_T5 : list_assemblies_classificiation_sources.xlsx
This table provides detailed information on each genome assembly used in the study, including group assignment, assembly label, haplotype number, species, population classification, sequencing technologies, assembly size, original FASTA file name, assembly method details, data provider or publication reference, and data repository.

Suppl.Table T6 : assemblies_distance_matrix.xlsx
This table contains the kmer distance matrix between all genome assemblies, computed from presence/absence variations across the pangenome. The values reflect the degree of shared genomic content between each pair of assemblies, as used for the clustering analysis in the main manuscript (Figure 1)

### Section 6: Complements on P. armeniaca Transposon library generation

The 11 genome assemblies used in this study — *RougeRH1, Marouch, Rojo_HCUR, Stella, KZ150H1, KR091H1, Meihua, CH240H1, CH250H1, Longwangmao,* and *Pmand CH264* — were selected to represent the major phylogeographic groups identified within the *Prunus armeniaca* complex and related species.

Among them, the first four (*RougeRH1, Marouch, Rojo_HCUR,* and *Stella*) correspond to European cultivars. Their inclusion reflects not only their agricultural relevance but also the known genetic sub-structuring within European apricots, as demonstrated by (Bourguiba *et al.*, 2012), who reported distinct western and central-eastern European clusters. This intra-European divergence justified the use of multiple assemblies from this region.

In addition, two assemblies (*KZ150H1* and *KR091*) represent Central Asian apricots, and one (*Meihua*) represents a Chinese *P. armeniaca*. Three others (*CH240H1, CH250H1,* and *Longwangmao*) correspond to *Prunus sibirica* individuals from China, and the last one (*Pmand CH264*) belongs to *Prunus mandschurica*.

This panel was thus chosen to maximize genetic divergence while maintaining balanced representation across all known genetic groups within and around *P. armeniaca*, forming a robust foundation for pan-genomic and transposon-based comparative analyses.

To support transposon annotation and enrichment analyses across the apricot pangenome, we constructed a curated and non-redundant transposable element (TE) reference library from the 11 genome assemblies included in this study.

- Suppl.table_T7: *CatlibApricot_withoutRedundancy.classif.xlsx*
  This table provides the full curated list of transposable elements identified across the assemblies. For each element, it reports: sequence name, length, strand orientation, classification at multiple taxonomic levels (class, order, Wcode, sFamily), coding status, structural annotation, and a confidence index (CI). This classification enables detailed downstream analysis and comparison across TE categories.

- Suppl.data_D1: *CatlibApricot_withoutRedundancy.fa*
  This FASTA file contains the non-redundant nucleotide sequences of the transposable elements identified across the 11 apricot assemblies. Each sequence appears only once in the dataset, even if detected in multiple assemblies, thereby avoiding duplication and bias in enrichment analyses. The sequences are annotated with detailed metadata, including sequence name, length, strand orientation, TE classification (class, order, Wcode, sFamily), confidence index (CI), and structural or coding features.

### Section 7 : Supplements on variants statistics

⇒ partie stats “SNP/indels générales” (section 7 : varinats général stats)

**Suppl.table_T8 : Structural and SNP Variation Summary.xlsx**

This Excel file contains four sheets summarizing the variation landscape across the pangenome:

1. raw_summary_insertions_deletions_total_with_chr_totals — A global summary of insertions and deletions, including per-chromosome totals.
2. SNPs summary — An overview of single-nucleotide polymorphism distribution.
3. normalized_indel_bin_distribution_per_chromosome — Distribution of indels per size bin and chromosome, normalized by aligned sequence length.
4. normalized_indel_group_contribution_per_bin/chr — The proportional contribution of each phylogeographic group to indel bins, also normalized by alignment length.

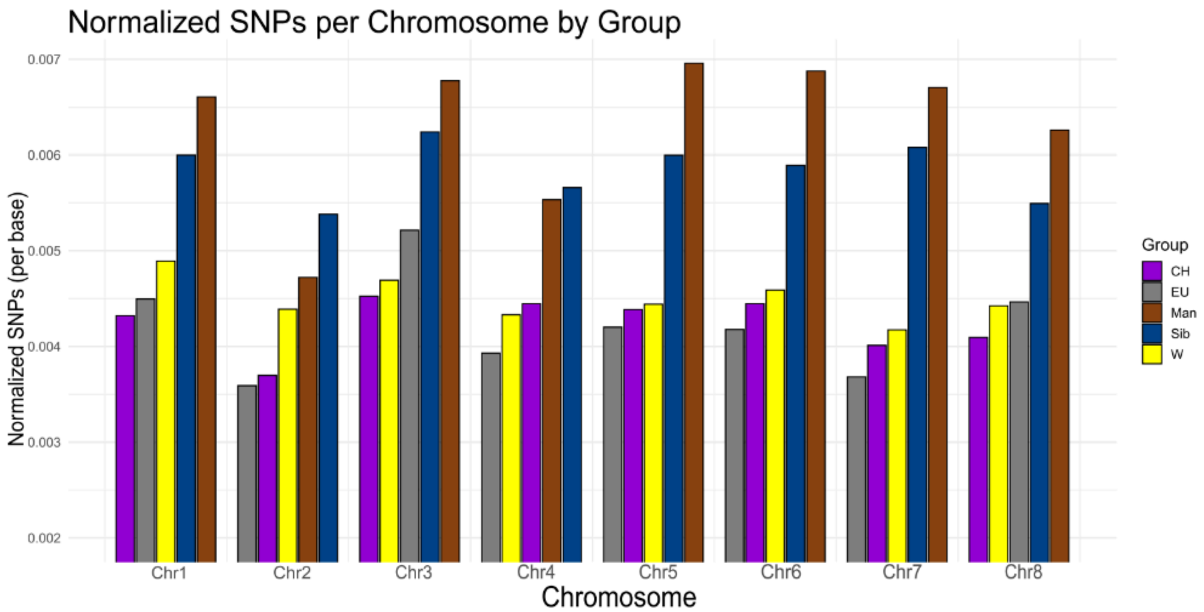


**Figure 3 : SNP frequencies extracted from the graph, relatively to reference Rojo_HCUR**. *Y-axis: SNP frequencies are normalized by chromosome lengths summed from all assemblies of each group.*

### Section 8 : Supplements on transposons analyses

**Suppl.table_T9 : Transposon Distribution and Enrichment Analysis.xlsx**

This file contains six sheets focused on the relationship between indels and transposon content:

1. raw_total_transposons_by_bin_and_chromosome — Absolute counts of transposons overlapping indels, broken down by size bin and chromosome.
2. total_transposons_by_bin_and_chr_normalized — The same data normalized by the total aligned bases per category, providing a measure of density.
3. bootstrap_proportions_top5_orders_by_bins — Bootstrap-derived confidence intervals for transposon order proportions across size bins.
4. pairwise_prop_test_top5_orders_by_bins — Pairwise proportion tests to compare transposon distributions between indel size bins.
5. pairwise_prop_test_top5_orders_by_chr — The same statistical comparisons across chromosomes.
6. prop_enrichment_order_vs_other_bins — An enrichment analysis comparing specific orders to the background distribution across size bins.

These sheets form the basis of the enrichment analyses shown in Figure 3, highlighting how transposon density varies according to indel length and chromosomal location.

##### Indel Size bins

In raw counts, mid-sized bins such as 200–400 bp (8,557 transposons) and 800–1,800 bp (7,100 transposons) have the highest number of insertions, suggesting a concentration of transposons in these indel classes  figure 4A). However, after normalization by the total aligned bases (figure 4B), the 50–200 bp bin becomes the most enriched, with the highest density (~0.00534 transposons per aligned base), indicating that short indels are disproportionately loaded with transposons. Conversely, longer indels (e.g., 5,000–100,000 bp) show lower densities (~$6.56 \times 10^{-4}$), despite having high raw counts, suggesting a more dispersed insertion pattern.

##### Transposon content per chromosomes

When looking at raw transposon counts per chromosome (figure 4C), chr1 leads with 8,431 insertions, followed by chr6 (5,768) and chr2 (5,517). These values largely reflect the size of the chromosomes. However, normalization by chromosome length reveals a different trend (figure 4D): chr8 shows the highest transposon density (~$2.14 \times 10^{-4}$), followed by chr3 (~$2.08 \times 10^{-4}$) and chr6 (~$1.99 \times 10^{-4}$). In contrast, chr5 has both the lowest raw count (3,354) and the lowest normalized density (~$1.77 \times 10^{-4}$), suggesting that it is overall less enriched in transposons.

##### Summary comparison

Raw counts reflect absolute abundance and are strongly influenced by region size.
Normalized densities reveal relative enrichment, highlighting biologically meaningful patterns that are not apparent from raw counts alone. For indels, short bins are the most enriched despite smaller absolute counts. For chromosomes, medium-sized ones (like chr8 and chr3) show the highest density, whereas chr5 stands out as both short and less enriched.

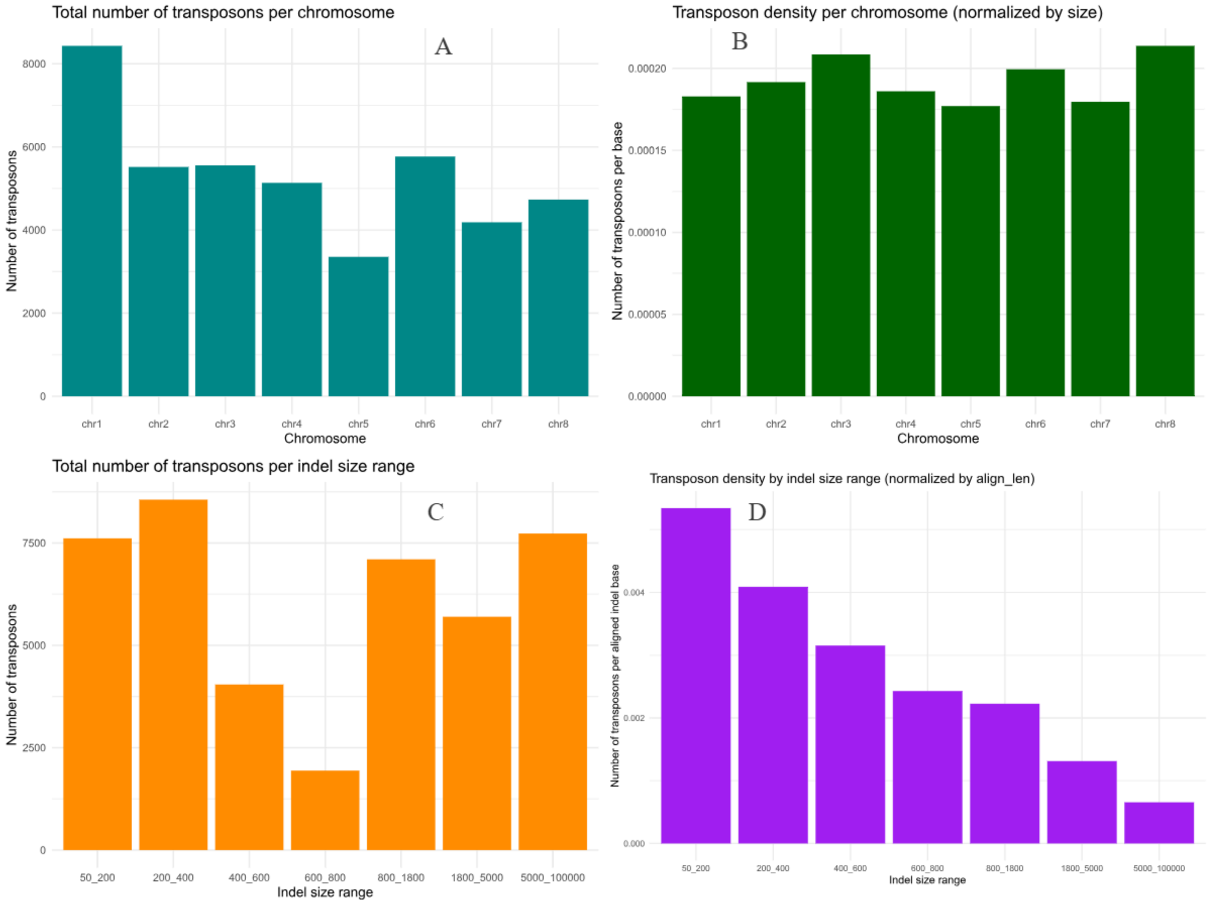


**Figure 4. Transposon distribution across chromosomes and indel size bins.**
*(A) Total number of transposons per chromosome. (B) Transposon density per chromosome, normalized by chromosome size. (C) Total number of transposons per indel size bin. (D) Transposon density per indel size bin, normalized by the total aligned indel length (align_len). While raw counts reflect absolute abundance, strongly influenced by region size, normalized densities reveal relative enrichment, highlighting distinct biological patterns across both chromosomes and indel size bins.*

### Section 9 : Filtering of reads after mapping

**Limitations of Pangenome Graph Tools in Handling SAM Flags**

We observed that current pangenome graph tools, specifically VG tools (v1.57), do not fully support all SAM flags. A notable limitation was identified in the 'vg surject' command; when projecting to references using a GAM file (e.g., from 'vg giraffe' output), information regarding read pairs is not retained. Consequently, pair-based filtering functionality is currently incomplete.

Furthermore, we did not observe any SAM flags associated with secondary or supplementary alignments in the output of 'vg giraffe'. While an option exists to output secondary alignments, preliminary testing indicated it may be non-functional, as its activation drastically reduced the quantity of mapped reads. It is however important to note these observations are specific to the version of VG tools used in this study, and ongoing active development may lead to rapid improvements in their functionality.

To illustrate the aforementioned issue, we performed a comparative analysis of Minimap2 and VG Giraffe using a small subset of samples. This comparison involved different FLAG-based filtering strategies applied to their respective mapping outputs.

- For Minimap2's BAM output, secondary and supplementary alignments were discarded (consistent with their lack of handling by VG tools), and unmapped reads were also excluded. Read pairs with only one mapped mate were retained, achieved by applying the flag filters '-f 4 -F 2304'.
- Similarly, for the VG Giraffe GAM output, subsequently processed by VG Surject to produce a BAM file, secondary and supplementary alignments were discarded (as VG would not set these flags). Unmapped reads were also removed, and read pairs with a single mapped mate were retained using the flag filter '-f 4 -F 2304'.

While the above limitations exist, it is possible to obtain the desired SAM flags by directly projecting during a VG giraffe run onto a pre-selected set of references. This process will lead to proper inclusion of these flags in the resulting BAM output. However, this approach requires executing the 'vg giraffe' command twice, as it currently lacks the functionality to output two distinct formats within a single run. Such a dual execution would incur prohibitive CPU costs, leading us to restrict this particular analysis to a subset of 50 short-read samples.

This small comparative analysis is based on six distinct conditions:

- Minimap2, with no multimapping, requires both mates to be mapped.
- Minimap2, with no multimapping, adhering to the "pair properly mapped" definition (which is software-dependent).
- Giraffe, with projection to all assemblies (ALLREFS), requiring both mates to be mapped.
- Giraffe, with projection to all assemblies (ALLREFS), adhering to the "pair properly mapped" definition.
- Giraffe, with projection exclusively to Rojo_HCUR (1REF), requiring both mates to be mapped.

- Giraffe, with projection exclusively to Rojo_HCUR (1REF), adhering to the "pair properly mapped" definition.

The outcomes of this comparative analysis are presented in the table and plot below. The corresponding SAM flags for each condition are detailed in the legend.

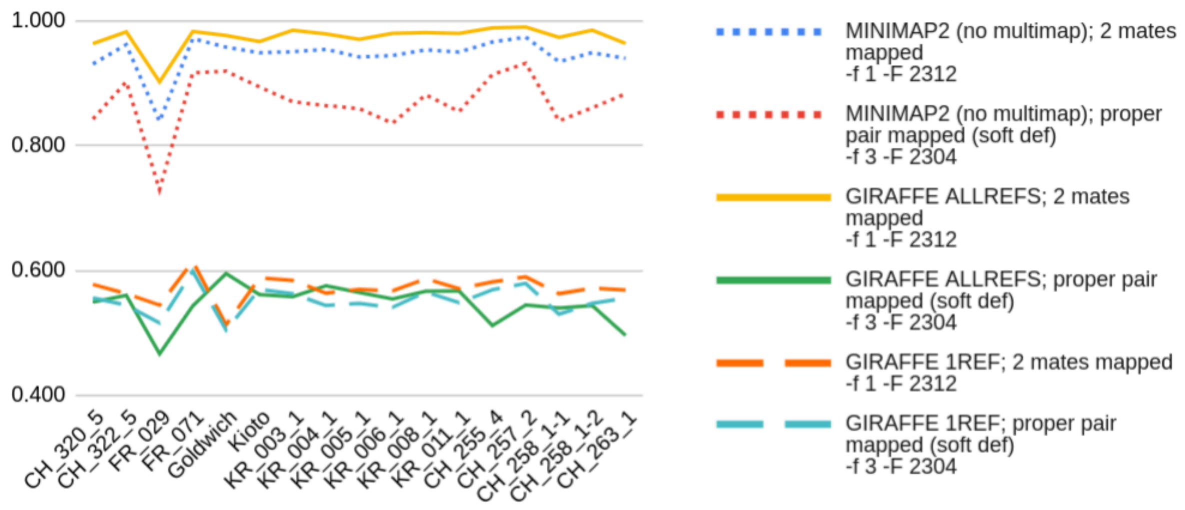


**Figure 5 : Comparative read-pair alignment performance across aligners, reference configurations and flag filters.**

*Blue dotted line — Minimap2, multimapping suppressed, pairs with both mates mapped (-f 1 -F 2312).*

*Red dotted line — Minimap2, multimapping suppressed, pairs flagged as properly mapped (-f 3 -F 2304, definition software-dependent).*

*Yellow solid line — VG Giraffe projected to all assemblies (ALLREFS), pairs with both mates mapped (-f 1 -F 2312).*

*Green solid line — VG Giraffe (ALLREFS), pairs flagged as properly mapped (-f 3 -F 2304).*

*Orange dashed line — VG Giraffe projected exclusively to Rojo_HCUR (1REF), pairs with both mates mapped (-f 1 -F 2312).*

*Light-blue dashed line — VG Giraffe (1REF), pairs flagged as properly mapped (-f 3 -F 2304).*

Note : Counts obtained with -f 1 -F 2312 reflect read pairs in which both mates are mapped and all secondary/supplementary alignments are removed. Counts obtained with -f 3 -F 2304 correspond to the SAM flag “properly mapped” category; this classification is aligner-specific and may include additional internal consistency checks.

Regarding the analysis of unpaired reads, our findings consistently demonstrate that graph-based mapping yields a higher proportion of paired alignments compared to conventional methods when projecting across all available assemblies (as evidenced by the yellow line versus the blue line). However, a significant limitation was observed with 'vg giraffe' under single-reference projection conditions (1REF), where nearly half of the read pairs failed to achieve concordant mapping. Even more unexpectedly, projection to the Rojo_HCUR reference (orange dashed line) resulted in only 50% to 60% of pairs remaining mapped, a proportion substantially lower than what was obtained with Minimap2.

These observations suggest that the current version of VG surject is unadapted to robust handling of BAM-like files and their associated flag-based filtering. This presents a bottleneck in results analysis, as the only alternative is the GAM binary format output by VG giraffe : a binary file that involves advanced programming expertise for effective exploitation.

### Section 10 : MAPQ scores distribution following mapping

MAPQ scores are designed to distinguish high-quality from low-quality mappings. However, the formula used to compute this value is author and software dependent. Thus, direct comparison using thresholds is not valid. See these explanations, for an interesting discussion on this point.

Moreover, as shown in the following figure, the MAPQ distribution generated by Minimap2 and VG Giraffe appears quite different. Strikingly, MAPQ peaks seem to appear every 15 MAPQ units on average, which we hypothesised to be an artifact of normalization by read length (reads from the 325 mapped accessions are 150bp long, on average). Note that this pattern is absent from Minimap2 MAPQ scores.

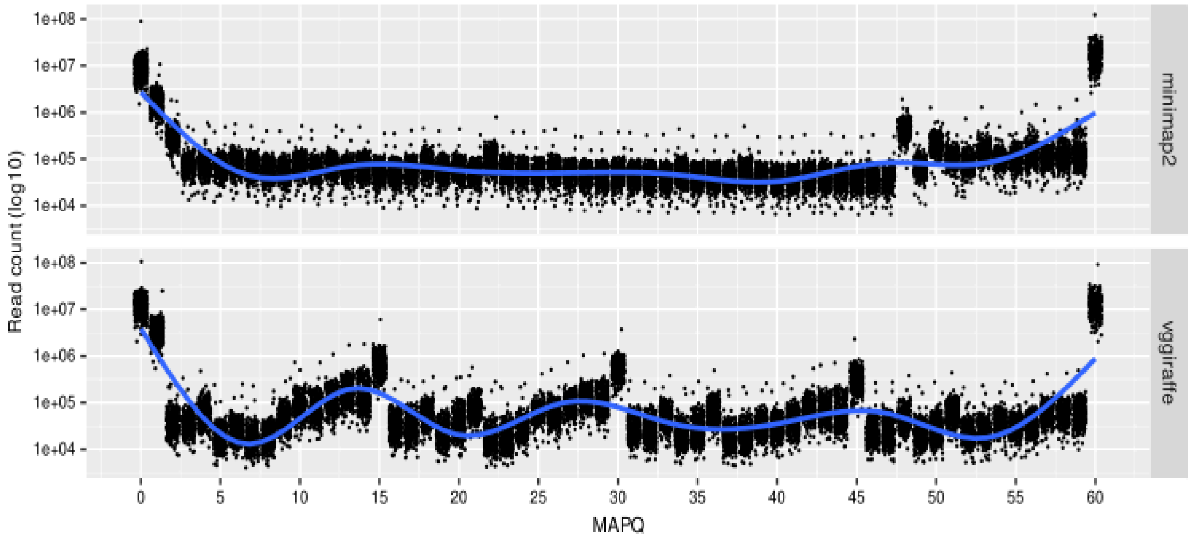


**Figure 6: Distribution of MAPQ scores, using all mappings from all accessions.** *For each MAPQ value, read count per accession are jittered. Blue line corresponds to the fitted Generalized Additive model (GAM) with smoothing parameters selected by restricted Marginal Likelihood (REML) estimation.*

### Section 11 : Computational cost of mapping

While VG tools enable a higher proportion of reads to be mapped, this advantage comes with a substantial computational burden. Across the various mapping configurations tested (detailed in section 8), the CPU cost ranged significantly. Mapping projected to the Rojo_HCUR reference incurred an average of 50 CPU hours per sample, whereas projection to all assemblies escalated the average to 175 CPU hours per sample.

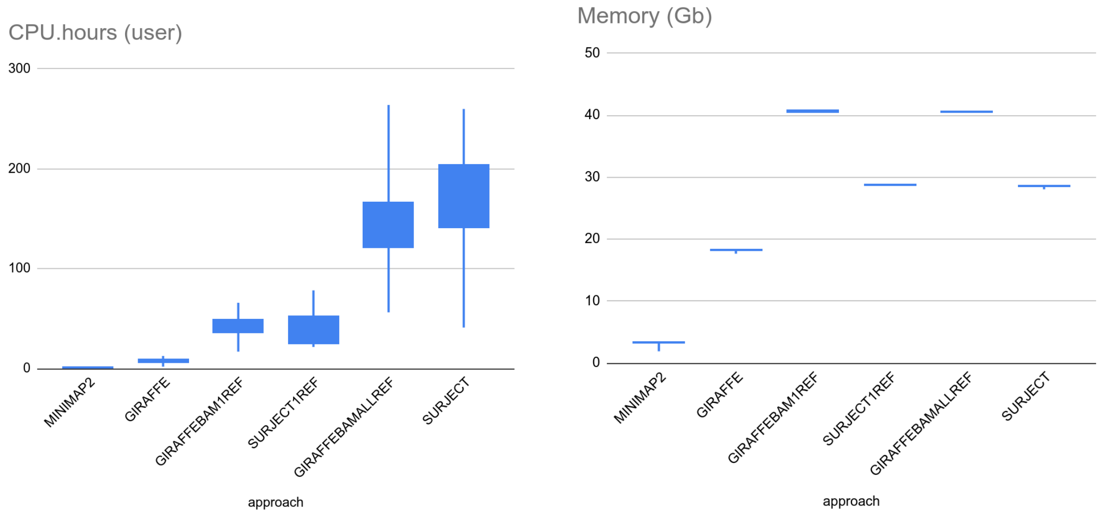


**Figure 6** : CPU and memory cost of the different mapping method (minimap and VG) and their different steps. *Minimap2 corresponds to linear mapping. Giraffe indicates direct graph mapping with internal BAM output and implicit surjection. GiraffeBAM1ref and GiraffeBAMallref use vg giraffe with internal BAM projection to a single reference path (Rojo_HCUR) or all available paths, respectively. Surject1ref and Surjectallref separate the pipeline, with vg giraffe producing GAM files followed by explicit surjection using vg surject to one or all reference paths*

Memory costs remained stable, as they are related to the size of the analyzed pangenome and associated indexes.

### Section 12 : Data relative to DAM Genomic Regions

For the selection of large indels highlighted in Figure 9 from the main manuscript, the coordinates of the corresponding graph loops can be found in Suppl.table_T10. It contains 2 sheets.

- Sheet 1: “positions_in_DAM_loops_assembly” contains said positions.
- Sheet 2: ”transposon_match_per_loop” : For each loop, we report the transposon matches.

To confirm the presence of the DAM region in the *Pmand CH264* assembly, despite its reported absence in the main manuscript, we performed BLAST searches on the raw reads. Results are available in Supplementary Table T11: *Blast_DAM1_to_6_ch_264_4.xlsx*.
